# The Expression of *Pax6* Genes in an Eyeless Arachnid Suggests Their Ancestral Role in Arachnid Head Development

**DOI:** 10.1101/2025.01.29.635487

**Authors:** Isabella Joyce, Austen A. Barnett

## Abstract

**Background:** Many animal lineages utilize *Pax6* transcription factors during eye development. Within Arthropoda, evidence suggests that *Pax6* genes are necessary for the specification of eyes in myriapods, crustaceans, and insects. However, recent data have argued that *Pax6* genes lack a role in the development of the eyes in Chelicerata (=arachnids, horseshoe crabs, and sea spiders). An alternative hypothesis argues that the absence of *Pax6* expression in developing chelicerate eyes could be explained by an earlier role for these genes in patterning eye precursor cells. The arachnid mite *Archegozetes longisetosus* lacks eyes, however it retains two *Pax6* paralogs in its genome. By leveraging these aspects of *A. longisetosus*, we tested the hypothesis that ancestrally chelicerates did not use *Pax6* genes to pattern their eyes but rather used them to pattern the central nervous system. We reasoned that if we observed comparable expression patterns of *Pax6* genes in *A. longisetosus* in comparison to those in arachnids that have retained eyes, then this would support the hypothesis that *Pax6* genes were not ancestrally used for eye specification in chelicerates.

**Results:** We followed the expression of canonical arthropod retinal determination genes to confirm that *A. longisetosus* does not develop vestigial eyes. We found that the expression of the *Pax6* paralogs was consistent with their roles in the development of the ocular region and central nervous system. By co-staining for these genes simultaneously with the conserved head patterning gene *orthodenticle*, we also observed early expression patterns of these genes in the protocerebrum of early *A. longisetosus* embryos that are comparable to those arachnids with embryonic eyes.

**Conclusions:** Our data provide support for the hypothesis that *Pax6* genes were not ancestrally used to pattern chelicerate eyes. The expression patterns of *Pax6* genes in *A. longisetosus* were comparable to those of other arachnids that have eyes. This suggests that the retention of *Pax6* genes in *A. longisetosus* is due to their ancestral, non-eye patterning roles. Further supporting this hypothesis is our observation that *A. longisetosus* does not pattern vestigial eyes. Lastly, our data suggests that the *Pax6* genes, with *orthodenticle*, acted to specify the ancestral arachnid protocerebrum.

## Background

Eyes have likely evolved independently multiple times within animals, underscoring their adaptive advantage in a variety of ecosystems [1,2]. These complex and diverse organs can be found across disparate metazoan lineages, including cnidarians, mollusks, vertebrates, and arthropods (reviewed in [3]). Ancestrally, the arthropods (=insects, crustaceans, myriapods and chelicerates) had a pair of multifaceted compound eyes, as well as multiple simple eyes called ocelli [4,5]. This basic theme is generally conserved across arthropods, albeit with lineage-specific modifications.

Evidence suggests that the arachnid lateral eyes are homologous to the insect compound eyes, whereas the medial eyes are homologous to the ocelli [6]. This ground plan differs from that of the likely sister group of arachnids, Xiphosura (=horseshoe crabs [7] but see [8,9] for an alternate hypothesis).

Instead, horseshoe crabs retain the basal arthropod state of housing both compound eyes and ocelli [4]. Sea spiders (=pycnogonids) also represent a non-arachnid chelicerate group, and these animals bear two pairs of median eyes, but lack lateral eyes [10,11]. At the other extreme of chelicerate visual systems lie groups that have lost their eyes entirely, including many members of Acariformes, the clade comprised of mites [12].

Despite the multiple occurrences of independent eye evolution across animal phyla, the development of these organs usually involves the utilization of Pax6 transcription factors [1]. Within arthropods, including chelicerates, phylogenetic evidence suggests that the last common ancestor of arthropods likely had at least two paralogous *Pax6* genes (see [13]), first identified as *eyeless* (*ey*) [14] and *twin of eyeless* (*toy*) [15] in the fruit fly *Drosophila melanogaster*. These distinct *Pax6* orthologs are used differentially across arthropod groups, obfuscating their ancestral roles in arthropod eye development. For example, in *D. melanogaster,* both *ey* and *toy* are expressed in the eye/antennal imaginal discs, which give rise to the adult eyes (note that *Pax6* input is not necessary for the development of its larval eyes [16]). In these imaginal discs, both *ey* and *toy* have been shown to be necessary for the development of the compound eye [14], whereas only *toy* is necessary for specifying the ocelli [17–19]. In another insect, the beetle *Tribolium castaneum*, both *Pax6* orthologs are required for the formation of the larval eyes, however RNAi of these genes showed only mild effects on the formation of the adult eyes [20]. In crustacean exemplars, the knockout of *ey* in *Daphnia magna* generated eye deformities [21]. In the decapod crustacean *Exopalaemon carinicauda*, the knockout of *ey* resulted in a range of compound eye deformities, whereas the knockout of *toy* had no effect on eye development [22]. Studies into the utilization of *Pax6* orthologs in myriapods is thus far restricted to an expression study in the millipede *Glomeris* marginata, where both *toy* and *ey* are expressed in the optic lobes, the regions that give rise to eyes in other arthropods [23]. These studies are suggestive of a conserved role of *Pax6* orthologs in at least some processes of eye development in the mandibulates, *i.e.*, the clade comprised of insects, crustaceans and myriapods.

The utilization of *Pax6* orthologs in chelicerates, however, is much less clear, as initially demonstrated by studies on horseshoe crabs, Current data suggest that the last common ancestor of Xiphosura underwent three rounds of whole-genome duplications [24–26]. These expansions of the horseshoe crab genome resulted in the retention of three *ey* orthologs and two *toy* orthologs in their genome [26]. Prior to these findings, an expression study in the Atlantic horseshoe crab *Limulus polyphemus* showed that a *toy* ortholog was not expressed in any of the eye anlagen during embryogenesis [27]. This discovery was surprising, as it was the first to suggest that a *Pax6* ortholog does not contribute to eye development in a chelicerate species. This unexpected revelation was further complicated by studies into the utilization of *Pax6* orthologs in the likely sister-clade to Xiphosura, the arachnids.

Within arachnids, the expression of *ey* and *toy* has been studied extensively in spiders. In the spider *Cupiennius salei*, *ey* (=*Cs-pax6a*) is expressed in the developing medial eyes. However, its *toy* ortholog (=*Cs-pax6b*) is only expressed in the anlagen of the optic neuropils that become associated with the medial eyes, and not the eyes themselves [28]. In the spider *Parasteatoda tepidariorum*, neither its *ey* (=*Pt-pax6.1*) nor its *toy* (=*Pt-pax6.2*) orthologs are expressed in any of the developing eyes but are rather expressed in the developing neural tissue that is adjacent to the anlagen of the anterior medial eyes [29,30].

A recent survey of eye development genes across seven spider species also showed no evidence of *Pax6* gene input into the development of their eyes [31]. Another study also showed *Pax6* expression in the developing head region of the opilionid (daddy-longlegs) *Phalangium opilio*. This work revealed that its *Pax6* paralogs are expressed in parts of the brain and its eyes. One paralog, *Po-Pax6a*, is expressed in the lentigenic layer of the eye at later embryonic stages, and both of its *Pax6* paralogs are mainly expressed in the neural tissue of the eye folds [32].

To explain the unexpected expression patterns of *Pax6* genes in spiders, it was hypothesized that the role of *ey* and *toy* in eye development may occur much earlier. This hypothesis suggests that early *Pax6* expression specifies photoreceptor precursor cells prior to their integration into the eye tissue at later stages [29,33,34]. Support for this hypothesis comes from early *Pax6* expression in the anterior rim of the ocular segments of the spider *P. tepidariorum*. In this spider, there were stripes of *Pax6* gene expression in the rim of the ocular region of early germ band stage embryos [34]. The expression of these two genes is associated with stripes of *orthodenticle* gene expression [34], a gene that is associated with early head development in a range of arthropod taxa (see [35] for review and arguments therein). These data are congruent with an alternative hypothesis for the ancestral use of *Pax6* genes in chelicerates, *i.e.*, that *Pax6* genes ancestrally patterned components of the ocular region of the head. The ocular region (note that this is also referred to as the “ocular segment”, however the segmental composition of this region is currently debated [35]) is the anterior-most body region along the antero-posterior axis of arthropods. During arthropod embryogenesis, the ocular region gives rise to the protocerebral portion of the brain, which includes the optic neuropils (see [36]). Outside of chelicerates, there is additional evidence that arthropod *Pax6* genes operate in patterning the ocular region. For example, in *T. castaneum*, both *ey* and *toy* are expressed in the ocular region, and act redundantly to specify the head lobes, or the bilateral outgrowths of the ocular segment that give rise to, among other structures, the larval eyes [37].

To distinguish what ancestral roles *Pax6* genes had in arachnid embryonic development, we followed the expression of these genes in the mite *Archegozetes longisetosus*. *A. longisetosus* is a member of the mega-diverse arachnid clade Acariformes, of which no studies into the expression or role of *Pax6* genes have so far been conducted. Furthermore, *A. longisetosus* has secondarily lost its eyes, a condition that has convergently occurred multiple times within Acariformes (see [38]). Paradoxically, a recent genome sequence revealed that *A. longisetosus* retains orthologs of genes commonly used in arthropod eye development, collectively called the Retinal Determination Gene Network (RDGN), including both *Pax6* gene paralogs, *ey* and *toy* [38]. By leveraging these aspects of *A. longisetosus*, we reasoned that, if ancestrally *Pax6* genes were used in the patterning of the ocular region rather than in the early establishment of eye components, we should see comparable early and late expression patterns of these genes between arachnids that have eyes and *A. longisetosus*. Alternatively, if *Pax6* genes were ancestrally used in the development of arachnid eyes, we would see greatly dissimilar early and late *Pax6* gene expression patterns in comparison to other studied chelicerates.

By following the expression patterns of *Al-ey* and *Al-toy*, we conclude that *ey* likely participated in the establishment of brain compartments, specifically the optic vesicles and the mushroom bodies of the protocerebrum. Furthermore, we show that the expression dynamics of *Al-toy* are consistent with a role in establishing the prosomal shield, a conserved arachnid structure that migrates late in development to cover the brain. Because testing these hypotheses hinges upon *A. longisetosus* truly lacking vestigial or embryonic eyes, we also visualized the expression patterns of orthologs of the RDGN gene components. These results show a lack of RDGN expression in any tissue associated with eyes, which supports the hypothesis that *A. longisetosus* lack vestigial eyes. We also followed the expression of the conserved head patterning gene *orthodenticle* in *A. longisetosus* simultaneously with *Pax6* expression. This provided further support for the role of *Pax6* genes in the development of the early *A. longisetosus* head/anterior region. Taken together, our results support the hypothesis that *Pax6* genes were likely not required for the development of the ancestral arachnid eye, but rather the morphogenesis of the arachnid protocerebrum.

## Methods

### Animal husbandry, embryo collection, and embryo fixation

Mites were reared on a plaster-of-Paris/charcoal substrate in plastic jars to maintain appropriate humidity. Mites were kept in these jars in an incubator at 25°C with wood chips to promote oviposition. Mites were fed with brewer’s yeast daily. Mite embryos were collected and fixed in the same manner described in[39]. Detailed protocols are available from AAB.

### Gene identification and bioinformatic analyses

The *A. longisetosus* orthologs of *wingless, peropsin, rhodopsin, eyes absent, Six3, sine oculis,* and *atonal* were identified previously in [38]. The ortholog of *orthodenticle* was identified in [40], and *dachshund* in [41]. To identify the potential *A. longisetosus* orthologs of *ey, toy, beta-arrestin,* and *myosin-III*, the *D. melanogaster* orthologs of each gene were used as queries for in a tBLASTn screen of the *A. longisetosus* transcriptome [38]. The resulting top hits were transcribed and subsequently aligned with selected metazoan protein sequences using MUSCLE with eight iterations [42]. These alignments were then used with PhyML [43] and the Smart Model Selection (SMS) tool [44] to construct phylogenetic trees. Branch support for these trees were also calculated using the approximate likelihood-ratio test (SH-like) [45] All trees were then edited to make publication-quality images using FigTree (v1.4.3). All phylogenetic statistics are reported in Table S1.

### Hybridization chain reactions and imaging

For all single and double hybridization chain reactions (HCRs), we followed the protocol developed by [46]. Probes specific to each mRNA were developed using the HCR 3.0 Probe Maker software [47]. The mRNA sequences used for probe production were from the transcriptome assembled in [38]. The identifiers of all transcripts used to design probes, as well as their associated HCR amplifiers, can be found in Table S2.

All resulting probes were ordered as oPools from Integrated DNA Technologies at a scale of 50 pmol per oligo. The probe sequences that were used in this study are listed in Tables S3-S17. If a transcript was too small to make the recommended 20 pairs of probes (*i.e., Al-arrestin-2* and *Al-peropsin*), we increased the probe concentration two-fold as recommended by [46]. Control HCRs were performed in parallel but lacked the addition of DNA probes. All HCR buffers and HCR amplifiers were purchased from Molecular Instruments. The amplifier fluorophores were also ordered from Molecular Instruments, and included fluorophores 594, 514, and 647 for use with amplifiers B1, B2, and B3, respectively. All HCR imaging was done on a Zeiss LSM 880 at Lehigh University, Bethlehem, PA. All images were processed in FIJI (v.2.9.0/1.53t), and all figures were assembled using Adobe Illustrator CS6.

## Results

### Development and compartmentalization of the *A. longisetosus* brain

In an effort both to establish a basis for comparable gene expression patterns between *A. longisetosus* and other study arachnids, and to determine if any vestiges of eye development are retained during *A. longisetosus* embryogenesis, we followed the embryonic development of the *A. longisetosus* brain. It is important to note that, unlike most emerging arthropod systems, *A. longisetosus* adults lay eggs at mixed developmental stages. This is because its oviducts serve as brood chambers, and therefore its clutches of eggs often contain embryos at different stages of development [48]. Consequently, the traditional “hours after egg laying” criterion cannot be used for this species. Instead, we use morphology to establish brain/head development-specific stages for the remainder of this paper, e.g. “<u>B</u>rain-<u>D</u>evelopment <u>S</u>tage-1 (=BDS-1).”

The arthropod brain is generally comprised of three cephalic domains. These domains, from anterior to posterior, are the proso-, proto-, and deutocerebrum [35] (however, see [49] for an alternate view). It is the region anterior to the deutocerebrum that the arthropod visual system develops from, and thus this region is often described in totality as the “ocular segment.” The “ocular segment” of chelicerates is also sometimes described as the “pre-cheliceral region” due to its location anterior to the cheliceral/deutocerebral segment. For consistency, we use the term “pre-cheliceral region” to describe the area anterior to the deutocerebral region, *i.e.*, the cheliceral segment, for the remainder of the paper.

As is typical of arthropod brain development, the first stage of brain morphogenesis begins with the pre-cheliceral region of *A. longisetosus* bifurcating into two lateral lobes, often called the “optic lobes” during BDS-1 (Fig 1A-A2). At BDS-2 (Fig. 1B-B2), we observed the appearance of “pits” in the pre- cheliceral region (Fig. 1B2, arrowheads). We take these “pits” to be invaginating neural precursor cells, based on their similarity to structures found in spider head development [28,50–53]. This stage is also characterized by medial boundaries forming around each of the optic lobes, resulting in an antero- posteriorly oriented “groove” between them (Fig. 1B2, dotted line demarks the boundaries of this structure). Also during BDS-2, we observed a pair of anterior-medial grooves, as well as two lateral grooves on each optic lobe. Based on comparative data from the spiders *Cupiennius salei* [28,50,54] and *Parasteatoda tepidariorum* [30,51], and the opilionid *Phalangium opilio* [32,55], we take the two lateral grooves to be the lateral furrows, and the two anterior grooves to be the anterior furrows (Fig. 1C-C2).

**Fig. 1.**
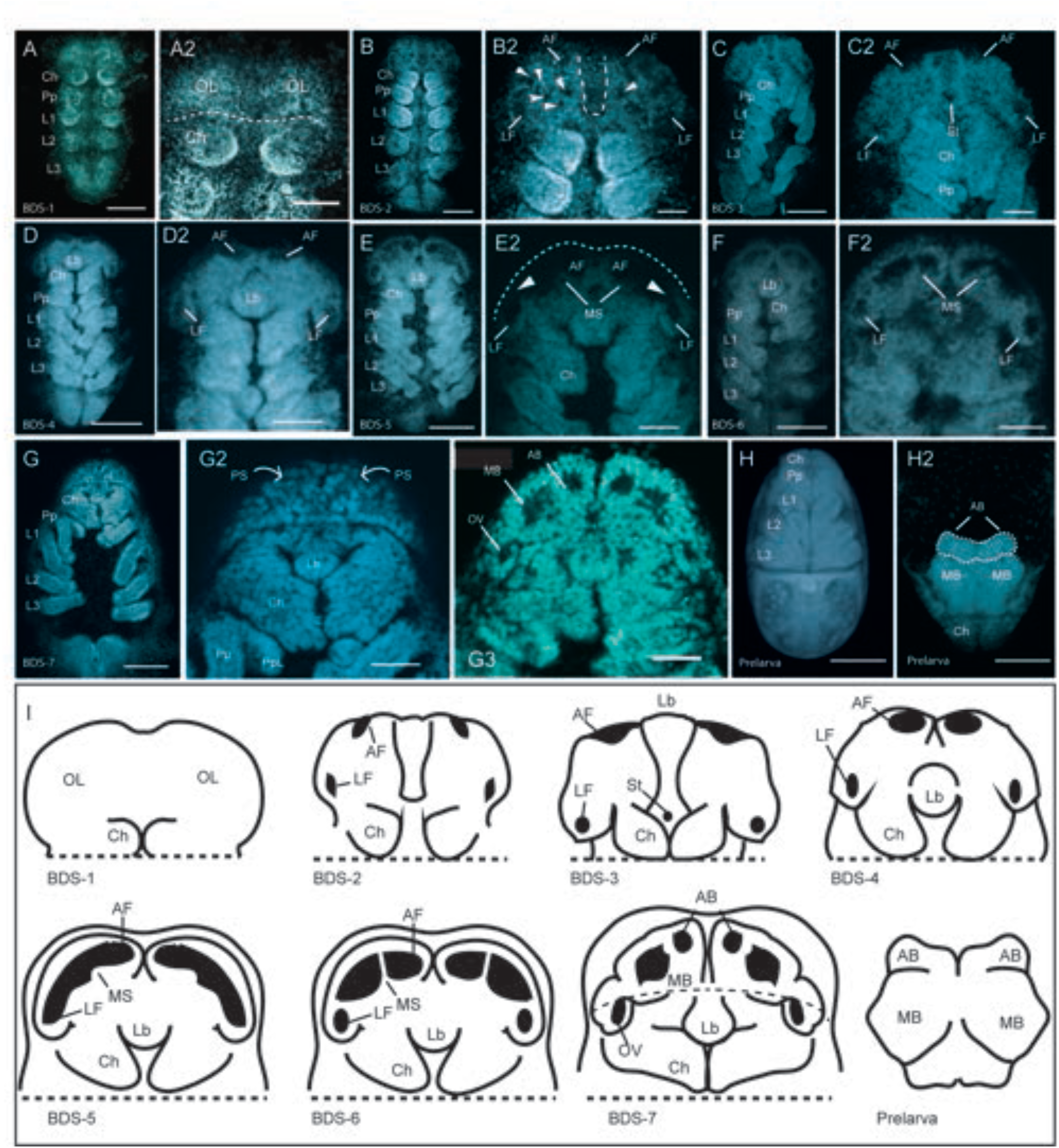
The development and compartmentalization of the *A. longisetosus* pre-cheliceral region**. A** An embryo at <u>B</u>rain <u>D</u>evelopment <u>S</u>tage 1 (BDS-1). **A2** The same embryo shown in **A**, showing the paired optic lobes (OL) of the pre-cheliceral region. The dotted line demarks the boundary between the deuterocerebral region and the pre-cheliceral region. **B** An embryo at stage BDS-2. **B2** shows the same embryo in **B**. The arrowheads point to “pits” in the presumptive neural tissue that are likely neural-precursor cells. Paired lateral furrows (LF) and anterior furrows (AF) are present at this stage, as well as a “groove” medially separating the optic lobes (dotted line). **C** An embryo at stage BDS-3. **C3** A closer image of the same embryo shown in **C**, showing the presence of the anterior and lateral furrows, as well as the newly formed stomodaeal opening (St.). **D** An embryo at stage BDS-4. **D2** A close-up of the head region of the same embryo shown in **D**. The labral halves have fused at this stage, and the labrum (Lb) has migrated posteriorly. **E** An embryo at stage BDS-5, and a close-up of this embryo, **E2**, showing the continuous opening formed by the fusion of the anterior and lateral furrows. The arrowheads mark a continuum between the lateral and anterior furrows, and the dotted line represents the anterior boundary of the embryo that is out of focus. Also at this stage, the medial subdivisions (MS) have begun to send out extensions that will eventually subdivide these continuous tubes. **F** An embryo at stage BDS-6, and **F2** a close-up of the head region of the same embryo. At this stage, the projections of the medial subdivisions have expanded to almost make contact with the lateral edges of the continuous tube of the lateral and anterior furrows. **G** An embryo at stage BDS-7. **G2** shows a ventral confocal 3D projection of this embryo. At this stage, the two halves of the prosomal shield (PS) have migrated ventrally and posteriorly to cover the developing brain region. The arrows show the movement of the prosomal shield halves. **G3** A confocal image of the same embryo in **G-G2** showing a more dorsal Z-slice. Embryos at this stage have subdivided the continuous tubes of the lateral and anterior furrows into the paired arcuate bodies (AB), mushroom bodies (MB), and optic vesicles (OV). **H** A ventral image of the prelarval stage of *A. longisetosus*. In **H2**, the embryo has been rotated so that its dorsum is shown. The brain region has migrated dorsally at this stage, and appears “upside-down” in comparison to the prior stages. The dotted line demarks the fused arcuate body. **I** Schema outlining the deduced morphogenetic events shown in **A- H2.** See text for details. All embryos shown in **A-H2** are oriented with the anterior of the embryo directed towards the top of the page, however please note that in prelarvae, the brain region is inverted as it folds over the head region. **I** Schematics of the aforementioned stages. Note that in the prelarval stage, the brain is “upside-down” in relation to the other stages. Abbreviations are Ch, chelicerae; L1-L3, walking legs 1-3, respectively; Pp, pedipalps. Scale bars in **A, B, C, D, E, F, G, H** and **H2** represent 50 µm. The scale bars in the remaining images represent 20 µm. All embryos shown in the confocal images were stained with DAPI.

BDS-3 is characterized by the deepening of the anterior and lateral furrows into the embryo. The antero-posterior groove that subdivides each ocular lobe was maintained at this stage, and a clear stomodeal opening was present at its posterior terminus (Fig. 1C-C2). BDS-4 follows the fusion of the labral halves, and their subsequent posterior migration (see [56] for details). At this stage, the anterior furrows were more pronounced, and visibly distinct from the lateral furrows (Fig. 1D-D2). At BDS-5, the anterior furrows expanded posterior-laterally and grew in size. Also at this stage, the anterior furrows made a continuous opening with their adjacent lateral furrows (Fig. 1E2, arrowheads mark a continuum between the lateral and anterior furrows). This is noteworthy, as a similar morphogenetic movement has not been observed in either opilionids or spiders, which maintain distinct lateral and anterior furrows during brain morphogenesis [28,30,32,50,51,55]. Also at BDS-5, we observed a field of cells that were growing towards the middle of each anterior furrow (Fig. 1E2). In the spiders *P. tepidariorum* and *C. salei*, two fields of neural precursor cells, called the medial subdivisions, also appear on each of the optic lobes and partially cover the anterior furrow [28,50,51,54]. Due to the similarity of these structures to those in the aforementioned spiders, we take these structures to be homologous to the medial subdivisions (Fig. 1E2). This is notable, as it has been proposed that the medial subdivisions give rise to the optic neuropils of the median eyes of *C. salei* (see Discussion in [28]).

At stage BSD-6, the lateral furrows have closed and have distinct boundaries (Fig. 1F-F2). The medial subdivisions extended anteriorly to contact the opposite sides of each anterior furrow (Fig. 1F2). In the spiders *P. tepidariorum* and *C. salei*, a second group of neural precursor cells, called the lateral subdivisions, migrate and partially cover the lateral furrows [28,50,51]. We did not observe any comparable morphogenetic movements or structures in *A. longisetosus*. This is also of interest, as these lateral furrows are presumed to form the lateral eye optic neuropils in these spiders[28].

In a number of arachnids, the non-neurogenic ectoderm at the anterior rim of the head lobes migrates to cover the developing brain. This “hood-like” structure is often called the prosomal shield [28,30,32,50,51,54,55]. We observed the same structure form in *A. longisetosus*, and its downward migration was mostly complete by BDS-7 (Fig. 1G-G2). Also at BDS-7, we observed that the medial subdivisions divided the anterior furrow into two distinct compartments. Based on observations of comparable stages of spider brain development, we take the anterior-most compartments to be the arcuate bodies (=central complexes) and the immediately posterior compartments to be the mushroom bodies (see [54], their Fig.4 compared to our Fig. 1G3). By using comparable stages in spiders, we also deduce that the lateral furrow forms the homolog to the spider optic vesicle (see[54], their Fig. 4 compared to our Fig. 1G3). We use these terms to describe these structures at this stage and in subsequent stages. In the following prelarval stage, the anterior of the brain “flipped” resulting in the anterior of the brain pointing posteriorly. This movement appears to be conserved in arachnids, as this is also seen in spiders and opilionids, as well as in an extinct chelicerate [8,32,54,57]. In Fig. 1H2, we demarcate the structures by dotted lines that we understand to be the arcuate bodies to highlight this morphological movement.

Taken together, the development of the *A. longisetosus* brain is similar to that of spiders and opilionids, albeit with lineage-specific differences. All groups develop both anterior and lateral furrows. However, these combine to form a continuous “tube” during mid-embryogenesis in *A. longisetosus*. These tubes are then subdivided to form specific compartments in the brain, *i.e.*, the optic vesicles, mushroom bodies, and the arcuate bodies. Surprisingly, the presence of structures associated with eye development in spiders are also present in *A. longisetosus*, despite their lack of eyes. These structures include the optic vesicles, as well as the formation of medial subdivisions that have been suggested to be the precursors of the optic neuropils of the median eyes [28].

### Developmental gene expression shows no evidence of vestigial eyes in *A. longisetosus*

Because of the similarities of *A. longisetosus* brain development to that of spiders, we asked if gene expression patterns could be used to detect embryonic eye primordia that may subsequently degenerate, in a similar manner that was used to discover the vestigial lateral eyes of *P. opilio* [32]. In all studied chelicerates, the eye anlagen originate from the non-neural ectoderm lining the peripheral rims of the head lobes [28,30,32,33]. As the prosomal shield migrates over the neural ectoderm, the eye anlagen presumably locate and connect to their associated optic neuropils (reviewed in [33]). Prior to the migration of the prosomal shield, the eye precursor cells can be identified by their expression of RDGN genes in the lateral, non-neural margins of the head lobes. To detect potential vestigial eye anlagen in *A. longisetosus*, we performed HCRs targeting its orthologs of the RDGN genes *eyes-absent* (*Al-eya*), *sine-oculis* (*Al-so*), *dachshund* (*Al-dac*), *six3* (*Al-six3*), and *atonal* (*Al-ato*). We also targeted the ortholog of *all-trans retinal peropsin* (*Al-peropsin*), which has been used to detect embryonic eyes in the spider *P. tepidariorum* [30]as well as in the opilionid *P. opilio* [32]. Aside from *peropsin,* the only remaining opsin gene that has been retained in the *A. longisetosus* genome is *rhodopsin* [38]. We therefore also used its potential expression patterns to detect possible vestigial embryonic eyes. Lastly, *beta-arrestins* are also important for arthropod photoreception [58], as is the *myosin-III* gene. Both genes have been used to mark the embryonic eyes of chelicerates [32,59]. We therefore targeted these genes to also identify potential vestigial embryonic eyes in *A. longisetosus*.

It is important to note that many of the RDGN component genes are expressed simultaneously in the non-neural and the neural ectoderm in the margins of the head lobes (*e.g.*, [32]). To date, there is no clear marker gene that can be used to distinguish the non-neural and neural ectoderm of the pre- cheliceral region. We were thus careful to make these distinctions to disambiguate the potential misidentification of vestigial eyes. Briefly, if gene expression was observed in the outer-most rims of the head lobes, we took this as evidence for that gene’s potential role in eye development rather that brain development. Below, we describe the resulting expression patterns of these aforementioned orthologs.

### Al-eyes absent expression

In *Drosophila*, the gene *eyes absent* (*eya*) encodes a protein tyrosine phosphatase that acts as a transcriptional co-activator with the products of *sine oculis* and *dachshund* within the RDGN [60]. The use of *eya* in patterning the eyes of arthropods appears to be ancestral, as exemplified by its expression in chelicerate eyes, *i.e.*, spiders and opilionids [28,30–32]. Unlike many of the RDGN genes, only a singleton copy of *eya* has been recovered in all surveyed spider taxa [31]. In the developing head of the spiders *P. tepidariorum* and *C. salei*, *eya* was expressed in the non-neural margins of the head lobes of early embryos. In each lobe, *eya* was detected in two separate domains, *i.e.*, an anterior and posterior domain. As the prosomal shields migrated, *eya* expression was enriched in the edges of the prosomal shield, and these cells were taken to be the primordia of the eyes. Upon the completion of prosomal shield migration, *eya* was expressed in all eye types of *P. tepidariorum* [30]. However, in *C. salei*, *eya* was only expressed in the secondary eyes (*i.e.,* all eye types to the exclusion of the anterior median eyes) [28]. This may be specific to *C. salei*, as a phylogenetic survey of a wide range of spider species showed that their *eya* orthologs are also expressed in all eye subtypes [31].

Outside of spiders, the only other chelicerate in which the embryonic expression of *eya* was surveyed was in the opilionid *P. opilio.* Like spiders, the earliest expression pattern of *Po-eya* was in the lateral margins of the head lobes. *Po-eya* was later expressed in the developing rims of the lateral furrows, as well as in the anterior furrows. This observation is interesting, as it was reported that there was no *eya* expression in the lateral furrows in the spider *P. tepidariorum* [30] nor in *C. salei* [28]. With the aid of high-resolution imaging of *Po-eya* expression, the authors were able to distinguish between non-neural and neural *Po-eya* expression. *Po-eya* expression appeared to be expressed in both the anlagen of the median and vestigial lateral eyes in the non-neural ectoderm as well as in the adjacent neural ectoderm. As the prosomal shield migrated, *Po-eya* expression was observed in the developing median and vestigial lateral eyes [32]. Outside of eye and brain development, the aforementioned studies on spiders and opilionids showed *eya* expression in the labrum, stomodaeum, segmental clusters of the ventral central nervous system, and also in what appears to be the mesoderm of the appendages. Taken together, the data from spiders and an opilionid suggest that *eya* expression can be used to detect both embryonic and potentially vestigial eyes.

Using HCR, we first observed the expression of *Al-eyes absent* (*Al-eya*) at the early stages of prosomal segmentation, when the cheliceral, pedipalpal, and the first two walking leg segments had formed (Fig. 2A-A3). At this stage, *Al-eya* expression was in each of the developing prosomal segments, as well as in a lateral domain in the pre-cheliceral region (arrowhead in Fig. 2A2). This lateral expression domain is similar to the triangular expression domains of *Al-ey* at this stage (see below) and may thus be involved in the formation of the same structure (*i.e.*, the lateral furrows; discussed below). We also observed *Al-eya* in an anterior ectodermal domain, which likely demarks the “groove” that separates the head lobes at BDS-2 (see Fig. 1B-B2). Interestingly, we also observed *Al-eya* expression in the presumptive mesoderm of the developing prosomal segments, as well as in the growth zone of the opisthosoma (Fig. 2A3). This is markedly different from the ectodermal expression of *Al-eya* noted above.

**Fig. 2.**
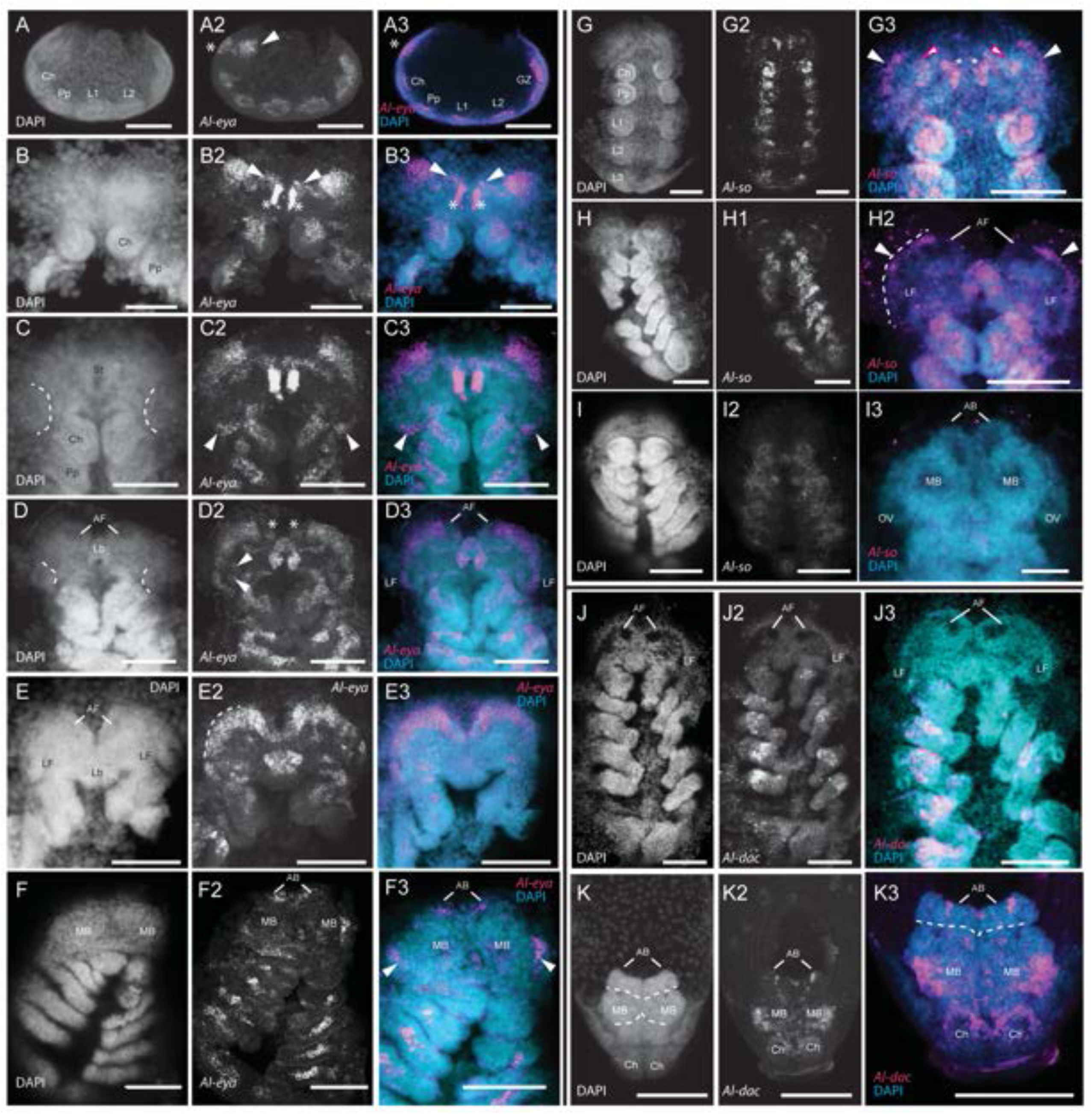
A*l-eyes absent (Al-eya), Al-sine oculis (Al-so),* and *Al-dachshund (Al-dac)* expression. **A-A3** Confocal images of *Al-ato* expression in an early segmental-stage embryo. **A** and **A2** are maximum projections, whereas **A3** is a Z-slice. The asterisks in **A2** and **A3** demark expression in the medial groove, whereas the arrowhead in **A2** points to expression in the lateral pre-cheliceral region. **B-B3** Confocal images of *Al-eya* expression in an embryo at BDS-1. The asterisks demark expression along the lines of the median groove, whereas the arrowheads demark thin lines of expression that appear at this stage. **C- C2** *Al-eya* expression in an embryo at BDS-2. The dotted lines in the DAPI image of **C** delimit the median boundaries of the lateral furrows where *Al-eya* expression appears at their posterior regions (arrowheads). **D-D3** *Al-eya* expression in an embryo at BDS-3. The DAPI image in **D** delimit the median boundaries of the lateral furrows. The asterisks demark *Al-eya* expression in the anterior furrows, and the arrowheads demark *Al-eya* expression in the medial lips of the lateral furrows. **E-E3** *Al-eya* expression in an embryo at BDS-4. The dotted line in **E2** demarks the boundary between the neural and non-neural head ectoderm. **F-F3** *Al-eya* expression in an embryo at BDS-7. The arrowheads in **F3** demark Al-eya expression underneath the prosomal shield in the mushroom bodies. **G-G3** *Al-so* expression in an embryo at BDS-2. Magenta-outlined arrowheads in **G3** demark expression in the medial grooves, and the white arrowheads demark *expression* in two lateral domains of the pre-cheliceral region. **H-H3** *Al-so* expression in an embryo at BDS-3. The dotted line in **H2** delimits the neural/non-neural ectodermal boundary, and the arrowheads demark expression in the lateral neural regions of the head lobes. **I-I2** *Al- so* expression in an embryo at BDS-7. At this stage, *Al-so* expression is not present in the developing head and brain. **J-J3** *Al-dac* expression at BDS-5, showing no expression in the head/brain region. **K-K3** *Al-dac* expression in the brain of a prelarva. All abbreviations are the same as in other figures. All scale bars represent 50 µm.

These expression patterns largely continued into BDS-1 (Fig. 2B-B3), however *Al-eya* was also expressed in two lateral expression domains that connected the bilateral triangular domains in the pre-cheliceral region (arrowheads in Fig. 2B2-B3) to the medial groove-lining domains (asterisks in Fig. 2B2- B3). We take these medial groove-lining domains to be the same ectodermal domain observed in Fig. 2A2 (asterisk).

At approximately BDS-2 (Fig. 2C-C3), the bilateral domains of *Al-eya* expression moved anteriorly. The previously described *Al-eya* expression domains were retained at this stage, however two new bilateral *Al-eya* expression domains appeared. These domains were restricted to the posterior margins of the lateral furrows (arrowheads in Fig. 2C2-C3). *Al-eya* expression was broadly retained in these domains at BDS-3. Intriguingly, however, the two domains of *Al-eya* expression merged into one, lateral domain on each of the head lobes (Fig. 2D-D3). In addition, two domains of *Al-eya* expression appeared on the medial “lip” each lateral furrow (arrowheads mark one pair in Fig. 2D2). *Al-eya* was also expressed in the anterior furrows at this stage (asterisks in Fig. 2D2). Embryos at BDS-4 displayed comparable *Al-eya* expression patterns to the previous stage (Fig. 2E-E3). However, it was at this stage that we were able to observe that *Al-eya* expression was confined to the neural ectoderm of the pre-cheliceral region (dotted outline in Fig. 2E2 demarks the presumed boundary between the neural and non-neural ectoderm). Because the eye anlagen in spiders and opilionids are derived from the non-neural ectoderm, these data support the hypothesis that *A. longisetosus* does not develop vestigial eyes.

Additional data supporting this hypothesis came from the expression of *Al-eya* in BDS-7 embryos (Fig. 2F-F3). At focal planes underneath the prosomal shield, we identified *Al-eya* expression in the lateral margins of the mushroom bodies (arrowheads in Figs. 2F2-F3), as well as in the arcuate bodies (asterisks in Figs. 2F2-F3). In these older embryos we did not detect any *Al-eya* expression in the non-neural prosomal shield that suggested either rudimentary or vestigial eyes.

### Al-sine oculis expression

In *Drosophila*, the Six-family gene *sine oculis* (*so*) encodes a transcription factor that interacts with several members of the RDGN and has been shown to directly form a protein complex with Eya (reviewed in [61]). Like many of the genes in the RDGN, all surveyed spider species retain two paralogs of *so* in their genomes [28,31], however the expression of these paralogs in the development of eyes varies across spiders. In all of the spider taxa investigated in [31], *so1* was shown to be expressed in all of the eye subtypes. A notable exception to this was the usage of *so1* in *C. salei* (=*Cs-six1a*), which was expressed in all of the eye subtypes to the exclusion of the anterior-median eyes [28]. The expression of the second paralog, *so2*, was present in all of the eye subtypes in the majority of the species examined by [31]. However, in the two species of spiders that belong to the clade Synspermiata, *so2* expression was not expressed in any of the eye anlagen. The variability of *so2* usage in eye development was also demonstrated by its expression in only the anterior-lateral eyes of *P. tepidariorum*. Lastly, the *C. salei so2* ortholog (=*Cs-six1b*) was expressed in all of the eye subtypes, to the exclusion of the anterior-lateral eyes [28]. Outside of spiders, the single copy of *so* in the opilionid *P. opilio* was expressed in the medial eyes as well as in the vestigial lateral eyes [32].

Like *P. opilio*, *A. longisetosus* has only a single-copy ortholog of *sine oculis* (*Al-so*). We first detected its expression in BDS-2 embryos (Fig. 2G-G3). *Al-so* was expressed in the presumptive mesoderm of the prosomal appendages, in a similar manner to *Al-eya* expression (see above). BDS-2 embryos showed *Al-so* expression in the medial grooves in a similar manner to *Al-eya* (Fig. 2G3, asterisks). *Al-so* expression was also detected in bilateral domains adjacent to the medial-groove expression domains (magenta-outlined arrowheads in Fig. G3) and also in two lateral domains (arrowheads in Fig. 2G3). At roughly BDS-3, *Al-so* expression remained similar to its expression at BDS-2. We did observe, however, that the *Al-so* expression domains at the lateral edges observed at BDS-2 expanded to line the edge of the neural ectoderm of each head lobe, as well as the lateral lip of each lateral furrow. We did not observe *Al-so* expression in the presumptive non-neural ectoderm that was suggestive of any eye primordia (Fig. 2H-H3). As development progressed into BDS-7, the expression of *Al-so* continued its expression in the mesoderm of the appendages (Fig. 2I-I3). However, in the pre-cheliceral region expression of *Al-so* (?) widely disappeared (Fig. 2I-I3). In summation, these data support the hypothesis that *A. longisetosus* embryos do not develop vestigial eyes during embryogenesis.

### Al-dachshund expression

In *Drosophila*, *Dm-dachshund* (*Dm-dac*) interacts with other components of the RDGN, and *Dm-dac* mutants lack eyes [62,63]. As with most of the RDGN genes, spiders have two paralogs of *dac*. The expression of these paralogs in spider eyes seems to be clade specific; however, in each species surveyed, at least one *dac* paralog is expressed in an embryonic eye [31]. In *P. opilio*, *Po-dac* is expressed in the medial eyes as well as the vestigial lateral eyes. Furthermore, RNAi targeting *Po-dac* results in the absence of the lateral eyes, without affecting the median eyes [32].

In *A. longisetosus*, we performed HCRs targeting its single-copy ortholog (*Al-dac*). As previously reported, *Al-dac* is expressed in the medial domains of the extending embryonic limbs [41]. However, we did not observe *Al-dac* expression in the pre-cheliceral region at any embryonic stage (an example is shown in a BDS-5 embryo in Fig. 2J-J3; earlier embryos not shown), consistent with [41]. We did, however, observe post-embryonic expression of *Al-dac* in the brains of prelarvae (Fig. 2K-K3). In these prelarvae, *Al-dac* was detected in regions we take to be the mushroom bodies, as well as in three small domains in the arcuate bodies (Fig. 2K-K3). Taken together, the lack of *Al-dac* expression in either the neural or non-neural ectoderm of the developing pre-cheliceral region supports the hypothesis that *A. longisetosus* does not develop vestigial eyes.

### *Al-Six3* expression

The six-family transcription factor Six3/Optix has a highly conserved role in demarking the anterior-most region of animal embryos [64]. In arthropods, this region has been proposed to be the prosocerebrum of the brain [35]. In addition to its role of anterior head regionalization, *Six3* is also involved in the formation of animal eyes (*e.g.*, [65]). In *Drosophila*, *Six3* is required for the progression of the morphogenetic furrow of the developing retinas in the eye/antennal imaginal discs [66]. Spiders have two paralogs of *Six3*, and in most spider species, one or both paralogs are expressed in at least one of the eye anlagen, except for the eyes of the spiders *A. geniculata* and *P. phalangioides* [31]. In the daddy-longlegs *P. opilio, Six3* is expressed in the developing median eyes [32].

In *A. longisetosus*, we observed *Al-Six3* expression in an early germ band stage (Fig. 3A-A3). At this stage, *Al-Six3* was detected in the anterior-most region of the embryo, consistent with observations in other animal taxa [64]. At approximately BDS-3, *Al-Six3* was observed in a large anterior domain that spanned and connected the anterior furrows (Fig. 3B-B3). We also detected *Al-Six3* expression in two domains within the neuroectoderm that we take to form the mushroom bodies (arrowheads in Fig. 3B2- B3) as well as in two domains that were in the region of the lateral furrows (arrowheads in Fig. 3B2-B3). As embryogenesis progressed to BDS-4, *Al-Six3* expression was retained in the neuroectodermal anterior furrows (Fig. 3C-C3). The mushroom-body associated expression was also retained (Fig. 3C2- C3, arrowheads). The expression of *Al-Six3* in the lateral furrows of this stage was restricted to the lateral edges of each furrow (Fig. 3C2-C3, asterisks). *Al-Six3* was also present in the labrum, as well as in the extending pedipalpal lobes (Fig. 3C2; note that these are underneath the chelicerae at this stage; see [41,56]). At approximately BDS-6, *Al-Six3* expression remained in the neuroectoderm of the pre-cheliceral region (Fig. 3D-D3). However, the expression of *Al-Six3* in the lateral furrows was markedly reduced and appeared to be restricted to their center (Fig. 3D2-D3, asterisks). Additional differences in *Al-Six3* expression from the previous stages include its expression in the distal tips of the first and second pairs of walking legs (Fig 3D2, arrowheads), as well as stronger expression in the extended pedipalpal lobes.

**Fig. 3.**
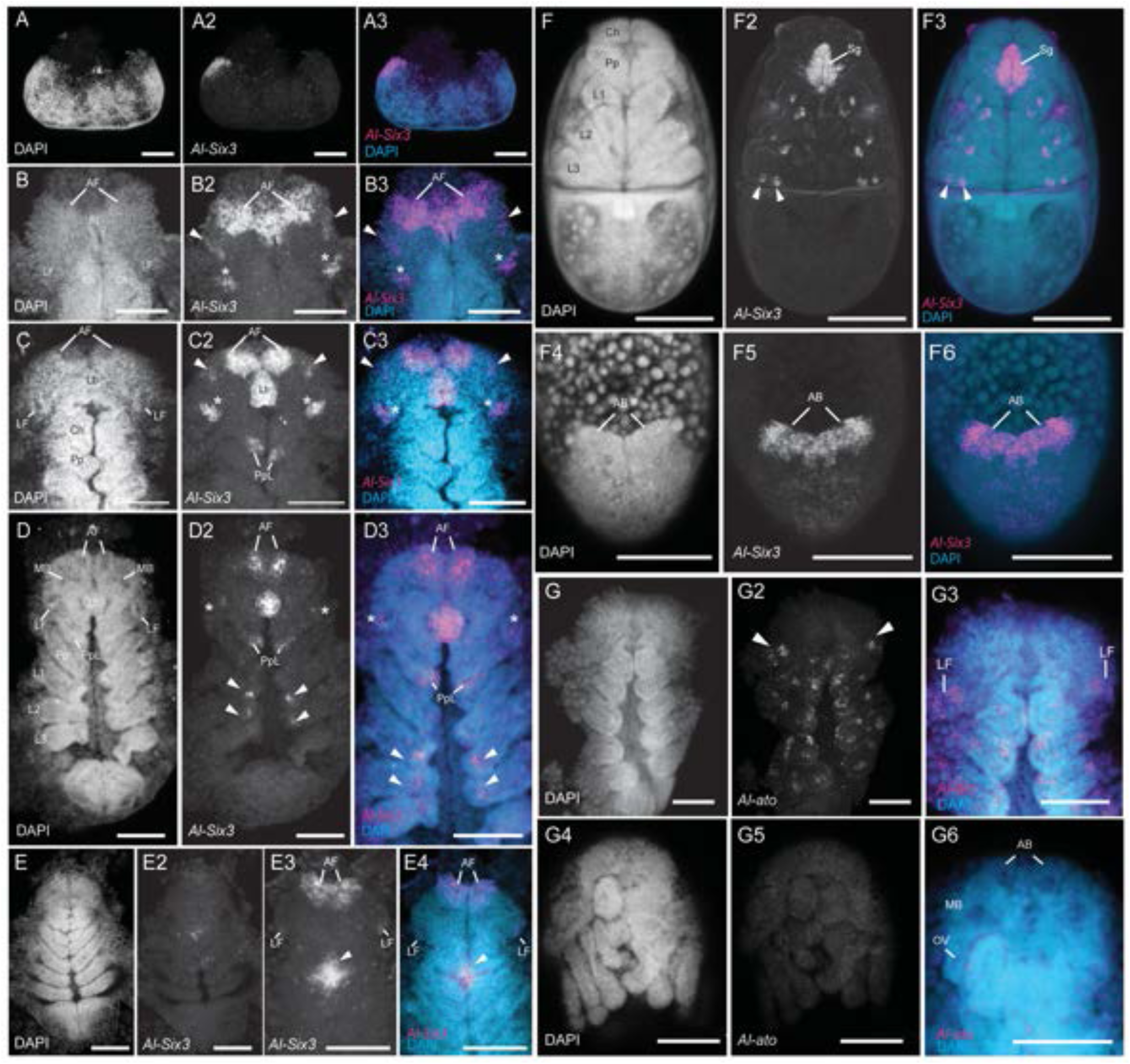
A*l-Six3* and *Al-atonal (Al-ato)* expression. **A-A3** *Al-Six3* expression in an early germ band embryo. **B-B3** *Al-Six3* expression at BDS-3. The arrowheads in **B2-B3** demark expression in the incipient mushroom bodies. **C-C3** *Al-Six3* expression at BDS-4. The arrowheads in **C2-C3** demark expression in the incipient mushroom bodies, and the asterisks demark expression in the lateral furrows. **D-D3** *Al-Six3* expression at BDS-6. Asterisks demark expression in the lateral furrows, whereas the arrowheads demark expression in the tips of the developing legs. **E-E4** *Al-Six3* expression at BDS-7. The arrowheads demark potential expression in the synganglion. **F-F3** *Al-Six3* expression in a prelarva. Note the expression in the synganglion (Sg). **F4-F6** *Al-Six3* expression in the same prelarva, rotated to the dorsum to visualize expression in the brain. **G-G3** *Al-ato* expression in an embryo at BDS-2. The arrowheads in **G2** demark expression in the lateral furrows. **G4-G6** An HCR of *Al-ato* showing lack of expression at BDS-7. All abbreviations are the same as in other figures. All scale bars represent 50 µm.

Also, *Al-Six3* expression in the mushroom bodies disappeared by this stage. In these aforementioned stages, we did not detect *Al-Six3* transcripts in the non-neural head ectoderm. This was made further evident by *Al-Six3* expression at BDS-7, at which point the prosomal shield has migrated over the neural ectoderm (Fig. 3E-E3). At BDS-7, *Al-Six3* expression was completely absent in the non-neural prosomal shield (Fig. 3E2). However, in Z-stacks deeper into the embryos, *Al-Six3* expression was still present in the anterior furrows/ arcuate bodies, as well as in the lateral furrows/optic vesicles (Fig. 3E3-E4). We also observed “patches” of *Al-Six3* expression in the midline of each embryo at this stage (Fig. 3E3-E4; arrowheads) which were absent in control experiments (not shown). In prelarvae, *Al-Six3* expression persisted (Fig. 3F-F6). *Al-Six3* expression in prelarvae was observed in two “spots” of expression in each of the walking legs (Fig. 3F2-F3; arrowheads demarcate two such spots in a third walking leg).

Additionally, *Al-Six3* was ubiquitously expressed in a structure whose position and shape suggest that it is the synganglion [67]. *Al-Six3* was also observed in the arcuate bodies at this stage, as well as in punctate domains in the anterior brain (Fig 3F4-F6). Taken together, our *Al-Six3* expression data also support the hypothesis that *A. longisetosus* does not have vestigial eyes.

### Al-atonal (Al-ato) expression

In *Drosophila*, the product of *atonal* (*Dm-ato*) is activated by the products of *Dm-so* and *Dm-eya* to initiate photoreceptor development (reviewed in [61]. In spiders, the *ato1* paralog seems to have a conserved expression domain in the anlagen of all eye subtypes, with the only exception being the spider *Segestria senoculata*, a member of the Synspermiata [31]. The same study also provided evidence that supports the hypothesis that the *ato2* paralog was ancestrally expressed in the primary eye primordia.

We detected the earliest expression of the singleton *atonal* ortholog in *A. longisetosus* (*Al-ato*) in BDS-2 embryos (Fig. 3G-G3). In these embryos, *Al-ato* was expressed in “clusters” of cells in the developing prosomal appendages, in a similar manner to both paralogs of *atonal* in the spider *P. tepidariorum* [29]. In the pre-cheliceral region, *Al-ato* expression was observed in two clusters of cells in each optic lobe, in regions straddling each presumptive lateral furrow within the neural ectoderm (Fig 3G2, arrowheads). We did not observe any *Al-ato* expression in the pre-cheliceral region that would be indicative of vestigial eye formation, *i.e.,* in a similar manner to spider *ato1* expression in the non-neural ectoderm of the head. In fact, *Al-ato* expression at this stage was most similar to the expression of *the ato2* paralog of the spider *P. tepidariorum* [29]. *Pt-ato2* was shown to be expressed in the pre-cheliceral region in two neuroectodermal clusters similar to our observations of *Al-ato* (see Fig 5I in[29]). We did not observe any changes in *Al-ato* expression throughout development, and its expression was never subsequently observed in new locations of the pre-cheliceral region (not shown). This trend ended in later BDS-7 embryos, at which all *Al-ato* expression ceased (Fig. 3G4-G6). Because spider *ato1* orthologs are expressed in the eye primordia of the prosomal shield at comparable stages [31], our data provide further evidence that *A. longisetosus* lack vestigial eyes.

**Fig. 4.**
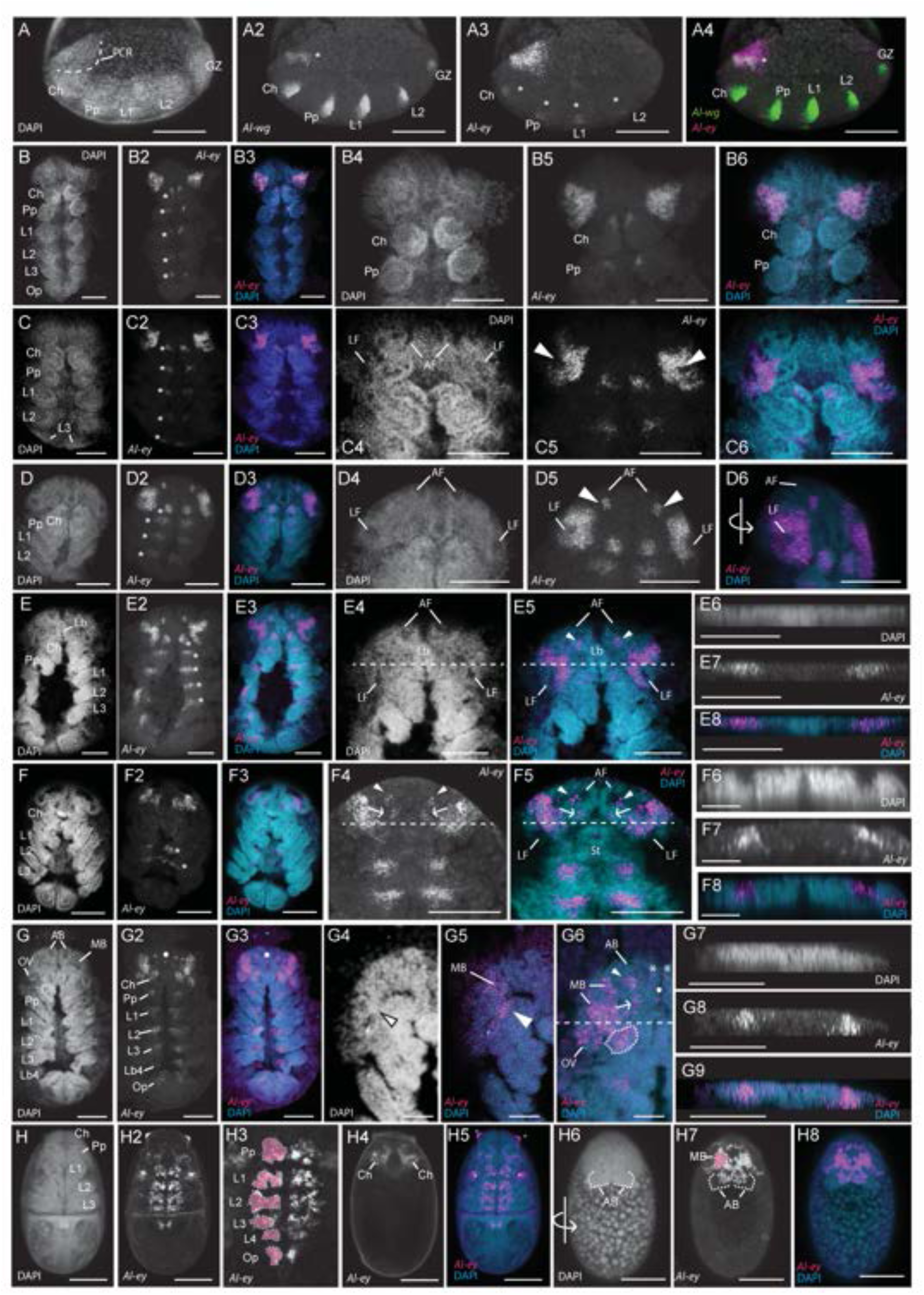
A*l-eyeless* (*Al-ey*) expression. **A-A4** Confocal images of a double hybridization chain reaction (HCR) in an early, prosomal-segmentation stage embryo targeting both *Al-wingless* (*Al-wg*) and *Al-ey*. All images of this embryo are oriented to show its lateral side, and its anterior is directed towards the left of the page. **A** DAPI nuclear counterstain image of this embryo, showing the location of the pre-cheliceral region (PCR), outlined in a dotted line. **A2** *Al-wg* is expressed in each of the developing prosomal segments at this stage, as well as in the segmental growth zone (GZ) and in a stripe of expression in the pre-cheliceral region (asterisk). **A3** *Al-ey* is expressed in this same embryo in a triangular-shaped domain of expression, and also in paired, clusters of cells in each of the prosomal segments (asterisks). **A4** A merged confocal image of this embryo showing *Al-wg* expression (green) concurrently with *Al-ey* expression (magenta). Note the co-expression of both of these genes in the pre-cheliceral region (asterisk). **B-B6** An embryo at a stage approximately between BDS-1 and BDS-2. **B** DAPI channel. **B2** *Al- ey* expression in the same embryo, showing the retention of *Al-ey* expression in triangular domains in the pre-cheliceral region, and also in paired clusters of expression in each of the prosomal segments (asterisks). **B4-B6** Confocal images of the same embryo, zoomed in to highlight the morphology of the pre-cheliceral region **(B4)** DAPI channel. **(B5)** *Al-ey* expression in this region. **(B6)** Both channels merged. **C-C6** Confocal images of a single embryo at BDS-2. **C** Nuclear counterstain with DAPI. **C2** Expression of *Al-ey* in this embryo, showing the retention of *Al-ey* expression in paired domains of the pre-cheliceral region, and also the paired expression in clusters of each prosomal segment (asterisks). **C3** Merged image of both the DAPI counterstain (blue) and *Al-ey* expression (magenta). **C4-C6** Confocal images of this embryo zoomed-in to show the appearance of the anterior and lateral furrows **(C4**, AF and LF, respectively). *Al-ey* expression “clears” from the lateral furrows at this stage (arrowheads, **C5**). **C6** Merged confocal image of DAPI (blue) and *Al-ey* expression (magenta). **D-D6** Confocal images of an embryo at stage BDS-3. **D** Nuclear counterstain with DAPI. **D2** Confocal image of *Al-ey* expression in this embryo, showing the retention of *Al-ey* expression in the pre-cheliceral region and in the paired clusters of the prosomal segments (asterisks). **D3** merged image of DAPI (blue) and *Al-ey* expression (magenta). **D4-D6** Zoomed-in confocal images of this embryo. At this stage, the anterior and lateral furrows are more distinct **(D4). D5** *Al-ey* expression is “ring-like” at this stage, as it surrounds the deepening lateral furrow. Two additional clusters of expression are also present at this stage (arrowheads), just posterior to each anterior furrow. **D6** A 3D-projection of this embryo, showing the merged DAPI (blue) and *Al-ey* (magenta) channels. The embryo has been rotated (see rotated arrow for orientation) to show the absence of *Al-ey* expression in the lateral furrows. **E-E5** Confocal images of a single embryo at stage BDS-4. **E** DAPI nuclear counterstain. **E2** *Al-ey* expression in this embryo, showing the retention of *Al-ey* in the pre- cheliceral region and in the paired, segmental clusters of the prosomal segments (asterisks). **E3** Merged image of DAPI (blue) and *Al-ey* expression (magenta). **E4** Zoomed-in image of the same embryo showing the DAPI nuclear counterstain of the pre-cheliceral region. **E5** *Al-ey* expression (magenta) merged with the DAPI counterstain (cyan) showing the “opening” of the *Al-ey* expression domains around the lateral furrows (see text for details). Arrowheads point to the *Al-ey* expression domains at the posterior of the anterior furrows. Dotted lines in **E4-E5** mark the position of the orthogonal slices shown in **E6-E8**. **E6** Confocal orthogonal slice through the antero-posterior axis of this embryo. **E7** *Al-ey* expression in the same orthogonal slice. **E8** Merged image of the DAPI (cyan) and *Al-ey* (magenta) confocal channels in the same orthogonal slice. **F-F5** Confocal images of an embryo at stage BDS-5. **F** DAPI nuclear counterstain of this embryo. **F2** *Al-ey* expression in this embryo. Note that the paired domains in the prosomal segments are retained at this stage, however only those of the second and third walking legs are visible (asterisks). **F3** Merged confocal image of this embryo showing the DAPI counterstain (cyan) and *Al-ey* expression (magenta). **F4** *Al-ey* expression in a Z-slice towards the dorsum of the same embryo. **F5** Merged confocal image of the same embryo, showing the DAPI counterstain (cyan) and *Al-ey* expression (magenta). In both **F4** and **F5**, arrowheads point to *Al-ey* expression in clusters associated with the anterior furrows, and the arrows point to newly-appeared clusters of *Al-ey* expression posterior to these. Dotted lines in **F4-F5** mark the position of the orthogonal slices shown in **F6-F8. F6** DAPI image of this orthogonal slice. **F7** *Al-ey* expression in this orthogonal slice, showing the internalization of *Al-ey* positive cells migrating inward. **F8** Merged DAPI and *Al-ey* channels in this orthogonal slice. **G-G6** Confocal images of an embryo at late BDS-6/early BDS-7. **G** DAPI image of this embryo. **G2** *Al-ey* expression in this embryo. **G3** Merged DAPI and *Al-ey* expression channels in this embryo. Note the appearance of paired clusters of *Al-ey* expression in the fourth walking leg segment bearing the fourth limb buds (Lb4) and also in paired clusters in the opisthosoma (Op). Furthermore, note the appearance of a new, central cluster of cells expressing *Al-ey* just above the labrum (arrowheads in **G2** and **G3**). **G4** DAPI image of this embryo, zoomed-in to show the structure of the pre-cheliceral region at this stage. The arrowhead points to the closing boundary separating the mushroom body from the arcuate body. **G5** Merged DAPI (cyan) and *Al-ey* (magenta) channels, showing expression in this closing boundary (arrowhead), and in the surface of the region above the mushroom body (MB). **G6** Dorsal Z-slice of this embryo, showing the internalized *Al-ey* expression in the pre-cheliceral region (see text for details). The horizontal, dotted line demarks the region of the orthogonal slices shown in **G7-G9**. **G7-G9** Orthogonal confocal slices through the pre-cheliceral region showing the internalization of *Al-ey* positive cells. **G7** DAPI channel. **G8** *Al-ey* expression. **G9** Merged DAPI channel (cyan) and *Al-ey* channel (magenta). **H-H8** *Al-ey* expression persists in the prelarval stage. **H** DAPI stain. **H2** *Al-ey* expression. **H3** *Al-ey* expression in the same prelarval. The paired, segmental *Al-ey* expressing clusters have become more complex. The left-most clusters are highlighted in pink and outlined. **H4** Z-slice showing the more dorsal cheliceral expression of *Al-ey*. **H5** Merged DAPI (cyan) and *Al-ey* expression (magenta) channels. Asterisks denote artefactual cuticle staining. **H6-H8** Confocal images of the same embryo, rotated to show the dorsally-migrated brain/pre-cheliceral region at this stage. **H6** DAPI stain; the arcuate bodies (AB) are outlined with a dotted line. **H7** *Al-ey* expression in this region, showing its expression in the mushroom bodies (MB) and the arcuate bodies. The left mushroom body is outlined and highlighted in pink. **H8** Merged DAPI (cyan) and *Al-ey* expression (magenta) channels. Scale bars represent 50 µm in all images, except in **F6-F8** and **G4-G6**, where they represent 20 µm. All other abbreviations are the same as in other figures.

**Fig. 5.**
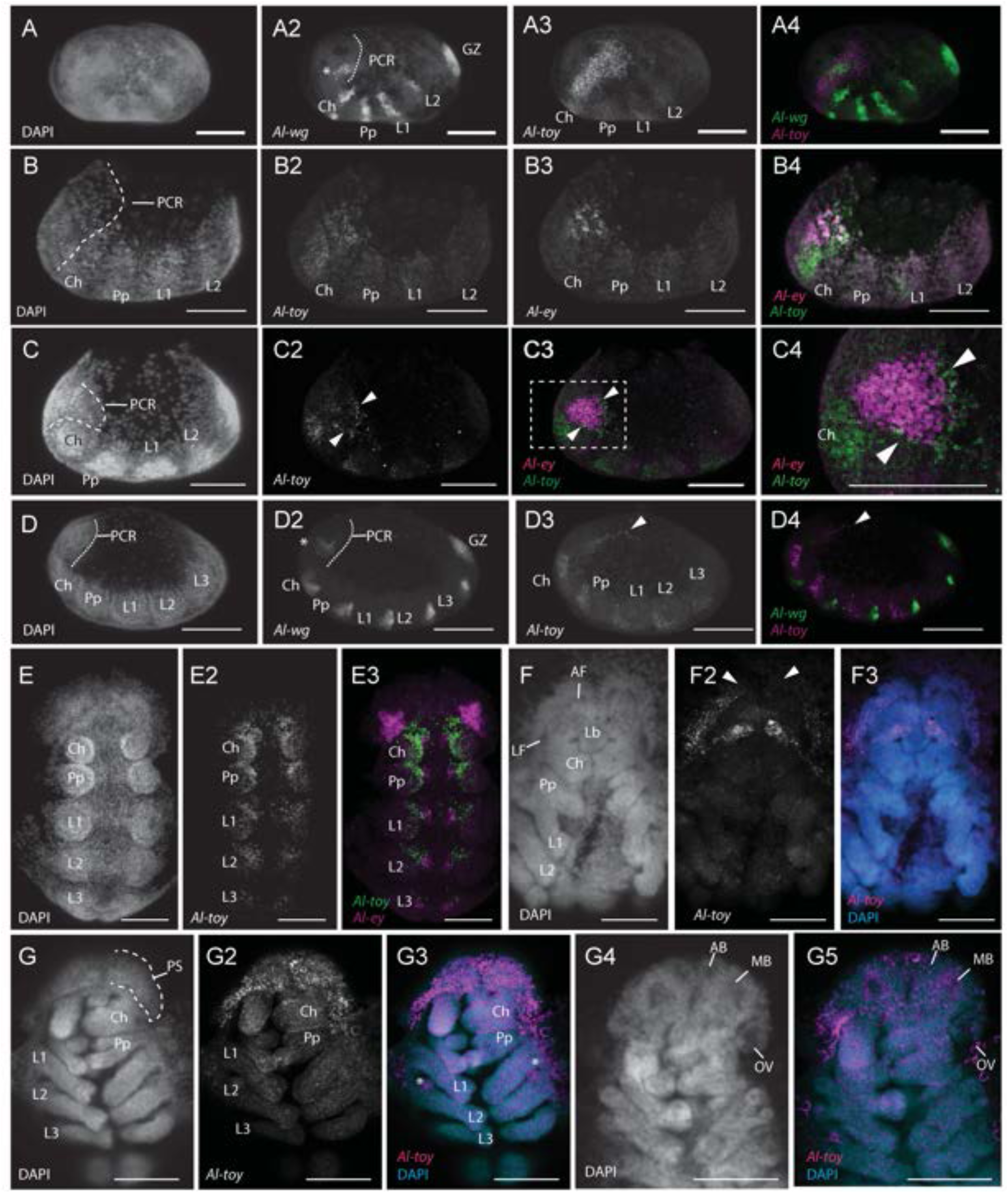
A*l-twin of eyeless* (*Al-toy*) is dynamically expressed throughout development. **A-A4** An early blastoderm/ prosomal segmentation stage embryo. **A** DAPI nuclear counterstain of this embryo. **A2** *Al-wg* expression in this embryo, showing its expression in stripes of the first four prosomal segments, in a stripe in the pre-cheliceral region (asterisk), and in the growth zone (GZ). The pre-cheliceral region (PCR) is outlined. **A3** Confocal image of *Al-toy* expression in this embryo. *Al-toy* is expressed in a broad domain in the cheliceral segment and its extension into the pre-cheliceral region. *Al-toy* was also weakly expressed in the developing prosomal limb buds. **A4** A merged confocal image showing simultaneous *Al-wg* (green)and *Al-toy* (magenta) expression. **B-B4** A slightly older blastoderm stage embryo. **B** DAPI counterstain of this embryo. The pre-cheliceral region is outlined in a dotted line. **B2** *Al-toy* expression in a broad anterior domain. **B3** *Al-ey* expression in the same embryo in a triangular domain in the pre- cheliceral region. **B4** *Al-toy* (green) and *Al-ey* (magenta) expression overlap at this stage. **C-C4** *Al-toy* and *Al-ey* co-expression at an early, four limb bud-stage embryo. **C** Nuclear DAPI counterstain of this embryo. The pre-cheliceral region is outlined. **C2** *Al-toy* is expressed in the anterior of each limb bud, and also in boundary-like domain (arrowheads). **C3** This *Al-toy* boundary (green) “outlines” *Al-ey* (magenta) expression. The dotted-lined box represents the region zoomed into in **C4,** which shows the mutually- exclusive expression domains of both *Pax6* orthologs. The arrowheads demark the *Al-toy* “boundary” type expression pattern. **D-D4** Confocal images of an embryo just prior to BDS-1. **D** DAPI counterstain of this embryo with the pre-cheliceral region outlined. **D2** *Al-wg* expression. **D3** *Al-toy* expression in the anterior of the prosomal limb buds, and in a thin domain in the pre-cheliceral region (arrowhead). **D4** Merged image of Al-wg and Al-toy expression. The arrowhead demarks the thin *Al-toy* expression domain in the pre-cheliceral region. **E-E3** Confocal images of an embryo at stage BDS-1. **E** DAPI counterstain of this embryo. **E2** *Al-toy* expression in the developing limb buds. Note the absence of expression in the pre- cheliceral region. **E3** Co-stain of both *Al-toy* (green) and *Al-ey* (magenta) expression. Note that both genes are not co-expressed at this stage. **F-F3** Confocal images of an embryo approximately at stage BDS-6. **F** DAPI counterstain of this embryo. **F2** *Al-toy* is expressed in the lateral non-neural ectoderm, in paired clusters above the labrum (asterisks), and in lines of expression connecting these to the lateral ectoderm (arrows). Arrowheads mark the anterior pre-cheliceral non-neural ectoderm that lacks *Al-toy* expression. **F3** Merged confocal image of the DAPI (cyan) channel and the *Al-toy* (magenta) channel. **G- G5** Confocal images of an embryo approximately at stage BDS-7. **G** DAPI counterstain of this embryo. The dotted line outlines one half of the prosomal shield (PS). **G2** *Al-toy* is expressed in migrating prosomal shield. **G3** Merged confocal images of DAPI (cyan) and *Al-toy* (magenta). The asterisks denote expression in the Claparede’s organs. **G4** DAPI counterstain of a Z-slice deeper into the embryo. **G5** Merged confocal images of DAPI (cyan) and *Al-toy* (magenta) in this Z-slice. *Al-toy* is notably expressed in the medullae of the mushroom bodies (MB). Embryos in **A-D4** are oriented with their anterior poles directed towards the left of the page. Embryos in the remaining images are oriented with their anterior poles directed towards the top of the page. All scale bars represent 50 µm. All other abbreviations are the same as in other figures.

### The expression of genes associated with downstream eye development does not support the presence of vestigial eyes in *A. longisetosus*

To ensure that we thoroughly tested for the absence of vestigial eyes in *A. longisetosus*, we targeted genes downstream of the canonical arthropod RDGN genes. In the opilionid *P. opilio* and the spider *P. tepidariorum*, the opsin gene *peropsin* is expressed in true embryonic eyes, as well as embryonic rudiments of the eyes of the opilionid after the completion of the migration of their prosomal shields [30,32]. Therefore, to detect the possible presence of rudimentary eyes in *A. longisetosus*, we performed HCRs targeting the *A. longisetosus* ortholog of *peropsin* (*Al-peropsin*). This experiment showed no expression of *Al-peropsin* at any stage of embryonic development, in the brain or otherwise (not shown). Alongside *Al-peropsin*, *Al-rhodopsin-7* was identified as the only other opsin retained in the *A. longisetosus* genome [38]. *rhodopsin-7* genes have been implicated in circadian rhythm photoreception in various taxa (reviewed in [68]). To test for the possibility that this gene may be expressed in developing, vestigial eyes, we performed HCRs targeting this gene’s expression. We also did not detect any *Al- rhodopsin-7* expression at any developmental stage in the pre-cheliceral region (not shown).

Beta-arrestins are utilized in photoreceptor specification, and their expression patterns have been recently used to identify the vestigial eyes of *P. opilio* [32]. By scouring the published *A. longisetosus* transcriptome [38], we identified three candidates for beta-arrestin orthologs. To verify these potential orthologs, a phylogenetic reconstruction was performed, which placed the transcript TRINITY_GG_5120_c51_g1_i7 in the same clade as Dm-Kurtz with high support (aLRT=0.90). The transcript TRINITY_GG_4713_c203_g1_i1 was placed in a clade with Dm-Arrestin-2 (aLRT=0.99), and the transcript TRINITY_GG_3318_c59_g1_i3 was placed in a clade with Dm-Arrestin-1 (aLRT=0.99) (Fig. S2). An HCR targeting all three genes showed no expression at any BDS in the pre-cheliceral region (not shown).

The expression of the *myosin-III* gene (known as *ninaC* in *D. melanogaster*) is expressed in the larval and adult eyes of the horseshoe crab [59], and the paralog *Po-myoIII-2* was used to detect the vestigial eyes of *P. opilio* [32]. By using several chelicerate and arthropod NinaC/Myosin-III proteins as queries, and by subsequently performing phylogenetic analyses of our possible hits, we were unable to detect any potential *myosin-III* orthologs in the *A. longisetosus* genome or transcriptome. This is interesting, as NinaC proteins in *D. melanogaster* are expressed in the photoreceptor cells, and their mutational abrogation results in photoreceptor defects [69]. Therefore, the absence of a *myosin-III/ninaC* in the eyeless mite is interesting for future studies into how natural selection targets photoreceptor genes in species undergoing eye loss or reduction. In summation, the expression of the RDGN genes as well as the absence of the expression of opsins and *beta-arrestin* together support the hypothesis that *A. longisetosus* lack embryonic or vestigial eyes.

### The identification and phylogenetic assessment of the *A. longisetosus Pax6* orthologs

We searched the recently published *A. longisetosus* genome and transcriptome [38] for putative *Pax6* orthologs using tBLASTn [70] and *D. melanogaster* Pax6 protein sequences as queries. This approach allowed us to identify two *Al-ey* transcripts that we refer to as *Al-ey.1* (TRINITY_GG_2648_c164_g1_i2) and *Al-ey.2* (TRINITY_GG_2648_c164_g1_i1), respectively (Fig. S1A). Both transcripts mapped to the same genomic locus (tig00005243_pilon), suggesting that they are isoforms rather than paralogs. The *Al- ey.1* isoform is 2,268 bps long, whereas the slightly longer *Al-ey.2* isoform is 2,354 bps long. Although *Al- ey.2* is longer, this difference is due to its lengthened 3’UTR. Surprisingly, *Al-ey.2* has a shorter coding sequence in comparison to *Al-ey.1, i.e.*, 670 bps and 1,401 bps, respectively. Therefore, the predicted amino-acid sequence of *Al-ey.1* has a 117 C-terminal extension that *Al-ey.2* lacks. By using Splign [71] to align these transcripts to the genome, we found that the differences between these two isoforms are a result of the differential use of exons towards the 5’ end of the gene. The *Al-ey.1* isoform is spliced from ten exons, whereas *Al-ey.2* is spliced from nine. The 3’ end of *Al-ey.1* is constructed from three exons, *i.e.*, exons 8-10. *Al-ey.2,* however, lacks exons 8 and 9, and instead uses an alternative exon upstream from exon 10. We denoted this exon as “Exon 8.5” due to its position between exons 8 and 9 (Fig. S1A). Using the above methodology, we also identified a single putative *Al-toy* transcript (TRINITY_GG_5245_c530_g1_i1; Fig. S1B). This 1,324 bp transcript is comprised of five exons, and maps to a distinct genomic locus (tig00005236_pilon) from that of the *Al-ey* transcripts.

The deduced protein sequences of the *Al-ey* and *Al-toy* isoforms were then evaluated with the NCBI Conserved Domain Database tool [72]. Both isoforms contain the three characteristic domains found in Pax6 proteins; the paired domain, the octapeptide-like domain, and the homeodomain (Fig. S1A- C) (see [73]). It was recently shown that arthropod Eyeless proteins contain a diagnostic lysine at position 64 in the linker region of the paired domain. This differs from arthropod Toy sequences, which instead have an arginine at this site [13,33]. The amino acid sequences of both *Al-ey* transcripts contain this diagnostic lysine residue (Fig. S1). Furthermore, the deduced Al-Toy amino acid sequence has the characteristic arginine at that site (Fig. S1C). These results support our hypothesis that the retrieved *Pax6* sequences are *de facto* distinct *eyeless* and *toy* orthologs.

To further test the hypotheses that these transcripts represent *bona fide* orthologs of *toy* and *ey*, we performed a maximum-likelihood phylogenetic assessment (PhyML) [43] of the deduced amino acid sequences of these transcripts with those from other metazoan *Pax6* orthologs (Fig. S1D). We also used the amino acid sequence of a putative *Pax2/5/8 A. longisetosus* ortholog (TRINITY_GG_4424_c37_g1_i1) as an outgroup in conjunction with other selected metazoan *Pax2/5/8* orthologs. This phylogenetic interrogation placed *Al-ey* and *Al-toy* in their predicted clades to the exclusion of *Al-Pax2/5* with high support (*i.e.*, aLRT scores of 0.98 for *Al-ey* and 0.96 for *Al-toy;* Fig. S1A). Taken together, our results support the identity of these transcripts as distinct singleton *eyeless* and *toy* orthologs.

### *Al-ey* expression

We initially performed HCRs simultaneously targeting *Al-ey* with the segmentation gene *Al-wingless* (*Al- wg*), which has been used as a marker for early segmentation stages in a variety of arthropods (reviewed in [74]). Using this methodology, we detected the earliest expression of *Al-ey* during the prosomal segmentation stage preceding BDS-1, at which the segments of the first four prosomal segments had been delineated (*i.e.*, the cheliceral, pedipalpal and first two walking leg segments; Fig. 4A-A4). *Al-ey* expression was observed in two paired, triangular-shaped domains within the pre-cheliceral region (Fig. 4A3). Embryos of this stage have an additional domain of *Al-wg* expression in the pre-cheliceral region (Fig. 4A2, asterisk), which has also been observed in a number of chelicerates during head/brain development [75]. Our double HCRs of *Al-ey* and *Al-wg* revealed that *Al-ey* expression in the pre-cheliceral region encompasses this *Al-wg* domain (Fig. 4A4).

At this early stage, we also observed paired clusters of expression in each of the developing prosomal segments (Fig. 4A3-A4; asterisks in 2A3). These expression domains of *Al-ey* were also observed at a later stage between BDS-1 and BDS-2 (Fig. 2B-B6) in the nascent tissue of the ventral nerve cord (Fig. 2B2, asterisks). This segmental expression of *Al-ey* was present in all subsequent stages of *A. longisetosus* embryogenesis, leading up to the prelarval stage.

At BDS-2, we observed the “clearing” of *Al-ey* expression in the newly formed lateral furrows (Fig. 4C-C6; arrowheads in C5; note the left lateral furrow in C4 is obfuscated by the surrounding tissue). This expression pattern was maintained in BDS-3, with *Al-ey* expression present in a ring-like domain surrounding the deepening lateral furrows (Fig. 4D-D6). We also detected additional expression in two clusters of cells just posterior to the anterior furrows (Fig. 4D5, arrowheads).

At BDS-4, the ring-like domain of expression was transformed into a “C-shaped” expression pattern, leaving *Al-ey* expression in the ventral-most cells of the periphery of the lateral furrows (Fig. 4E- E8). This appeared to be the result of the ring of expression observed in BDS-3 “breaking” at its dorsum. Additionally, *Al-ey* expression was maintained in the previously aforementioned clusters of cells posterior to the anterior furrows (Fig. 4E5, arrowheads). In arachnids, the process of brain development involves the internalization of neural tissues (*e.g*.,[54]). Therefore, we also asked if, and when, a comparable internalization occurs in *A. longisetosus* through visualizing orthogonal views of *Al-ey* stained embryos. Using this method, we observed that at BDS-4, the bilateral *Al-ey* positive cells were still embedded within the surrounding tissue (Fig. 4E6-8).

At late BDS-5, we identified *Al-ey* expression in clusters of cells on the ventral margins of the region of the “opened” lateral furrow, as well as the insides of this combined lateral and anterior furrows (Fig. 4F-F8). This suggests that the *Al-ey* expressing cells in the periphery of the lateral furrow at BDS-3 and 4 internalized as the lateral and anterior furrows form a continuous tube. To explore this further, we also imaged *Al-ey* expression along an orthogonal plane in a similar region to that shown in Figs. 2E6-8. This revealed the presence of *Al-ey* expressing cells surrounded by the edges of the “tube” made by the fusion of the lateral and anterior furrows (Fig. 4F6-8). *Al-ey* expression was also retained in two small domains at the posterior-lateral region of each anterior furrow (Fig. 4F4-F5; arrowheads). Additionally, another pair of small domains appeared at this stage posterior to the initial pair (Fig. 4F4-F5; arrows).

At late BDS-6/early BDS-7, the *Al-ey* expression patterns became more complex. *Al-ey* expression was maintained in segmental clusters in the developing central nervous system (CNS), however additional clusters appeared in the fourth walking leg segment as well as in the opisthosoma (see [39,76] for explanations on the divergent posterior segmentation in this species). Within the developing brain region, *Al-ey* expression was mostly internalized, however some external (*i.e.*, surface-level) expression did remain. Of note, a small cluster of *Al-ey* positive cells emerged in the center of the pre-cheliceral region, just anterior to the labrum (Fig. 4G2, G3 and G6, dots). External expression was also found at the closing border of the continuous anterior and lateral furrows at the site the lateral furrows’ anterior border (Fig. 4G4-5, arrowheads). Additionally, we observed external *Al-ey* expression in the region above the incipient mushroom bodies (Fig. 4G4-5). Further towards the dorsal Z-axis, we observed *Al-ey* expression in the posterior-lateral region of the arcuate bodies (Fig. 4G6, arrowhead). We take these to be the same *Al-ey* expressing cells that we observed in a similar location at BDS-5 (Fig. 4E5 and F5). The two clusters of *Al-ey* expression just posterior to these were also retained at this stage (Fig. 4G6, arrow; compare to Fig. 4F4 and F5, arrows). Two additional clusters of *Al-ey* expression were likewise seen in the medial portion of each arcuate body (Fig. 4G6, asterisks). *Al-ey* positive cells were also found in the newly compartmentalized optic vesicles, as well as inside of the incipient mushroom body region of the closing tubes (Fig. 4G6). An orthogonal view along the frontal plane revealed that these *Al-ey* expressing cell clusters of the mushroom bodies took on a triangular shape, with their vertices pointing ventrally (Fig. 4G7-9). This was in contrast to the shape of these clusters in BDS-5, and may indicate a pattern of internal migration via changes in cell shape, (*e.g.*, apical constriction) as the continuous lateral furrow/anterior furrow tubes close.

We also detected *Al-ey* expression in the post-embryonic prelarval stage, where its segmental expression was maintained from earlier stages (Fig. 4H-H8). *Al-ey* expression in the segmental clusters of the CNS became more complex, likely reflecting the differentiation of the neural cells expressing this gene. It is also important to note that the segmental CNS expression of the cheliceral segment moved to a more dorsal position. *Al-ey* was also expressed in the dorsal-most region of the brain, occupying the same space as the arcuate body, and also in larger and more anterior paired expression domains, which we take to be the mushroom bodies (Fig. 4H6-H8).

Taken together, our results indicate that *Al-ey* expression is expressed in the developing lateral furrows/optic vesicles, the mushroom bodies, and the components of the anterior furrow/arcuate body during embryogenesis. Furthermore, *Al-ey* expression persists in the CNS of post-embryonic stages. We did not observe *Al-ey* expression at these stages in the tissues taken to be the precursors to the embryonic eyes, *i.e.*, the non-neural ectoderm of the head lobes.

### *Al-toy* expression

As with *Al-ey*, we initially co-stained embryos for *Al-toy* expression simultaneously with the segmentation gene *Al-wingless* (*Al-wg*). We detected the earliest expression patterns of *Al-toy* during a similar early blastoderm stage as shown in Fig. 4A-A4, *i.e.*, when the first four prosomal segments had been delineated by *Al-wg* (Fig. 5A-A4). Also at this stage, the aforementioned pre-cheliceral stripe of *Al-wg* was present. *Al-toy* was expressed in a broad domain that extended from the anterior to the posterior of the embryo. The anterior of this domain appeared to be restricted to the cheliceral segment, and its posterior domain broadened into the pre-cheliceral region where it covered the pre-cheliceral *Al-wg* stripe (Fig. 5A3-A4). *Al-toy* was also expressed weakly in the developing limb buds (Fig. 5A3).

In spiders and daddy-longlegs, the *Pax6* orthologs are often expressed at the same embryonic stage, and appear to have specific early-stage expression domains [28–32,34]. We therefore performed HCRs simultaneously targeting both *Al-ey* and *Al-toy* to resolve when both orthologs are potentially co-expressed during pre-cheliceral development. We observed the co-expression of *Al-ey* and *Al-toy* in embryos at the same stage shown in Fig. 5A (Fig. 5B). At this early germ-band stage, we observed *Al-toy* expression in the pre-cheliceral region (Fig. 5B2), indicative of its slight anterior migration from the cheliceral segment (see Fig. 5A-A4). Low-level *Al-toy* expression was also present in the developing prosomal appendages. Interestingly, the triangular *Al-ey* expression domain (first shown in a slightly later stage in Fig. 4A-A4) completely overlapped with this *Al-toy* domain (Fig. 5B3-4B4). This is comparable to the co-expression of both *toy* and *ey* in the spider *P. tepidariorum* during its stage 8.1 and 8.2 [34], and thus likely represents a conserved feature for these genes in arachnids.

Subsequently, when the prosomal limb-buds have begun to grow more distinct, the overlap between the *Pax6* genes decreased (Fig. 5C-C4). The expression of *Al-ey* was retained in its anterior, triangular domain. However, *Al-toy* was expressed at the margins of the pre-cheliceral region, *i.e.*, at the anterior cheliceral segment boundary, and at the lateral boundaries marking the lateral edges of the presumptive ocular lobes (Fig 5C2 and C3; arrowheads). Co-staining with *Al-ey* revealed that this pre-cheliceral expression of *Al-toy* encompassed the triangular domains of *Al-ey* expression, with *Al-toy* expression being “cleared” from the region of *Al-ey* expression (Fig. 5C3 and C4). This is remarkably similar to the expression of *toy* and *ey* in the spider *P. tepidariorum*, specifically at its stage 9.1 [34]. In this species, *ey* is also expressed in a triangular domain at its stage 9.1. As in *A. longisetosus*, its *toy* ortholog appears to also encompass the *ey* expression domain [29,34]. The high degree of similarity of these *Pax6* expression patterns *in A. longisetosus* compared to spiders supports our hypothesis that arachnid *Pax6* orthologs are acting early in the development of the pre-cheliceral region, and not the development of the eyes.

*Al-toy* expression was also retained in the first four prosomal limb buds; however, it was restricted to the anterior of each developing limb (Fig. 5C-C3). This observation is interesting, in that *toy* expression was not observed in the developing limb buds of the spider *C. salei* [28]. However, in the spider *P. tepidariorum*, *toy* may be expressed in a similar manner to our observations of *Al-toy*. The previous expression reports of the *P. tepidariorum toy* ortholog primarily focused on its expression in the developing head. However in [29], there does appear to be some *toy* expression in the prosomal appendages (see their Fig. 5J). Furthermore, a recent study into RDGN genes in the opilionid *P. opilio* did not reveal *toy* expression in its appendages either [32]. Therefore, the role of *toy* in the developing arachnid limb may be a labile feature of arachnid evolution. Nevertheless, further taxonomic sampling is needed to test these hypotheses.

At the onset of BDS-1, *Al-toy* expression was maintained in the anterior portion of the first four pairs of limb buds, and also appeared in the third walking leg buds (Fig. 5D-D4). We also observed *Al-toy* expression in a thin domain trailing from anterior to posterior from the pre-cheliceral region. Based on the position of this trailing domain, as well as its position near the pre-cheliceral *Al-wg* stripe (Fig. 5D2-D4), we take this to be the remnant of *Al-toy* expression seen in the pre-cheliceral region of the preceding stages (Fig 5D3 and 3D4; arrowheads). At late BDS-1, all pre-cheliceral expression of *Al-toy* was absent, however *Al-toy* expression in the developing limbs persisted (Fig. 5E-E4). We noticed that the expression of *Al-toy* in the developing limbs was reminiscent of the segmental expression of *Al-ey* in the embryonic midline (Fig. 3C-C3). We therefore asked to what extent *Al-toy* was co-expressed with *Al-ey* at this stage. Our double HCR targeting both transcripts revealed that these paralogs were not segmentally co- expressed, with *Al-toy* being restricted to the limb buds, and *Al-ey* being restricted to the CNS (Fig. 5E3).

*Al-toy* expression was not detected in subsequent stages (not shown) until approximately BDS-6 (Fig. 5F-F3). This is notable, as *toy* orthologs are expressed at comparable stages in spiders [28–31] and opilionids [32]. At BDS-6, *Al-toy* expression was absent from the developing appendages*. Al-toy* transcripts were detected, however, in the lateral boundaries surrounding the optic lobes, as well as in the dorso-lateral margins of the lateral furrows. In spiders, these lateral margins have been described as the non-neural ectoderm that eventually migrates to form the prosomal shield (e.g., [28,50,51]). *Al-toy* was also expressed in a pair of domains just above the labrum (Fig. 5F2, asterisks), and also in a line of cells connecting these domains to the lateral optic lobe domains (Fig. 5F2, arrows). *Al-toy* expression was also absent from the anterior-most region of the optic lobes (Fig. 5F2, arrowheads). Comparable expression patterns are not seen for *toy* in either spiders [28–31] or opilionids [32].

At approximately BDS-7, *Al-toy* expression was ubiquitous in the prosomal shield (Fig. 5G-G3), confirming our hypothesis that *Al-toy* expression in the previous stage (i.e., Fig. 5F-F3) was in the non-neural ectoderm of the head lobes. This is striking, as neither *Pax6* ortholog is expressed in the developing prosomal shield in spiders. However, in the opilionid *P. opilio*, its *toy* ortholog was expressed in the leading margin of the migrating prosomal shield ([32]; their Fig. S2). Thus, our observations may represent a lineage-specific use for *toy* in mites. We additionally observed *Al-toy* expression in the Claparede’s organs. These organs are modified coxal extensions of the second walking legs that act to aid in water uptake in *A. longisetosus* larvae (see [56] for notes on their development). Also, deeper into the embryo, we observed punctate expression of *Al*-*toy* in the developing brain, and also its expression in the interior medullae of the mushroom bodies (Fig. 5G4-G5). We did not observe *Al-toy* in any subsequent stages, including the prelarval stage, following BDS-7 (not shown).

Together, our observations show a large degree of divergence in *Al-toy* expression in comparison to *toy* expression patterns in other studied arachnids. These differences include its expression in the developing prosomal appendages, its absence in the pre-cheliceral region following early stages, its ubiquitous expression in the late prosomal shield, and its expression in an acariform mite-specific appendicular structure, the Claparede’s organ. As was the case with *Al-ey* expression, we did not observe *Al-toy* expression in any of the eye-generating tissues.

### Early co-expression of the Al-Pax6 paralogs and the head-patterning gene orthodenticle

The similarities in *Pax6* expression between mites and spiders during the development of the pre- cheliceral region, coupled with the absence of vestigial eye primordia in *A. longisetosus*, suggests that these genes likely did not participate in eye development in the ancestral arachnid. However, they do suggest an ancestral role in the development of the arachnid pre-cheliceral region and its non-eye derivatives. As mentioned previously, in many chelicerates investigated, there appears to be no *Pax6* ortholog expression in any of the eye primordia, with the exceptions being *eyeless* expression in the anterior-median eyes of *C. salei* [28], and the late expression of *Po-Pax6a* in the median eyes of *P. opilio* after the migration of the prosomal shield [32]. To explain the widespread absence of *Pax6* expression in the embryonic spider eyes, or their associated neuropils, it was hypothesized that the cellular precursors to eye primordia or photoreceptors may be specified or triggered by the early expression of *Pax6* genes prior to their differentiation in later developmental stages. Another proposed explanation is that the *Pax6* genes play no role in spider eye development, but they instead play a role in the patterning other aspects of the pre-cheliceral region [29,31,33].

In a wide array of arthropod exemplars, *orthodenticle* orthologs are co-expressed with *Pax6* genes in the protocerebral region of the brain (see [35] and arguments therein for a summary). We reasoned that, if we observed early *Al-ey* and/or early *Al-toy* co-expression with *Al-otd,* this would be indicative of a role for these *Pax6* genes in specifying the protocerebrum instead of eyes, as *A. longisetosus* appears to lack eye primordia (see above). Alternatively, if *Pax6* genes ancestrally specified cells destined to be associated with the eyes during the early stages of development, we should see an absence of *Pax6* expression, and their potential co-expression with *orthodenticle*, at these early stages in the eyeless mites. We chose to directly compare our observations to those of spiders, as the earliest expression patterns for *otd* in opilionids has thus far been reported for stages 8 and later [32], at which the pre-cheliceral region has already been established and has become morphologically complex. It is also important here to note that the lineage leading to spiders underwent a whole-genome duplication [77], which has resulted in at least two *otd* orthologs in spiders [31]. In the spider *P. tepidariorum*, *orthodenticle-1* is necessary for the development of the spider pre-cheliceral region [78]. Also, a recent single-cell RNAseq study showed that *Pt-otd1* is expressed with the *Pax6* orthologs in cell clusters marking the pre-cheliceral region [34]. We therefore performed HCRs simultaneously targeting the *Pax6* genes and the singleton *A. longisetosus orthodenticle* ortholog (*Al-orthodenticle; Al-otd*) in early germ band mite embryos.

Following the transition from radial to bilateral symmetry in *P. tepidariorum* embryos, *Pt-otd1* is expressed in the posterior-most boundary of the pre-cheliceral region, adjacent to the cheliceral segment [78,79]. Similarly, the earliest expression of *Al-otd* that we observed was in early germ-band embryos, where it was expressed in a continuous domain in the embryonic anterior, as well as in the cells incipient ventral nerve cord (=VNC, Fig. 4A-A2). This anterior *Al-otd* domain overlapped with the bilateral *Al-ey* expression domain at this stage, specifically at the lateral margins of *Al-otd* expression (Fig. 6A3, A5, A7, A8, and A11). Furthermore, *Al-otd* was expressed in the same cells as *Al-toy* at this stage, however its co-expression with *Al-otd* was more extensive than that of *Al-ey* (Fig. 6A4, A6, A9, and A12). This early expression of *Al-toy* extended more medial-ventrally than that of *Al-ey*, where it overlapped *Al-otd* expression in all but the medial domains of *Al-otd* (Fig. 6A4 and A6). Taken together, these early expression patterns of *Al-ey, Al-toy*, and *Al-otd* are similar to those observed in the spider *P. tepidariorum* at roughly its Stage 8 during the early development of the pre-cheliceral region[34,78].

**Fig. 6.**
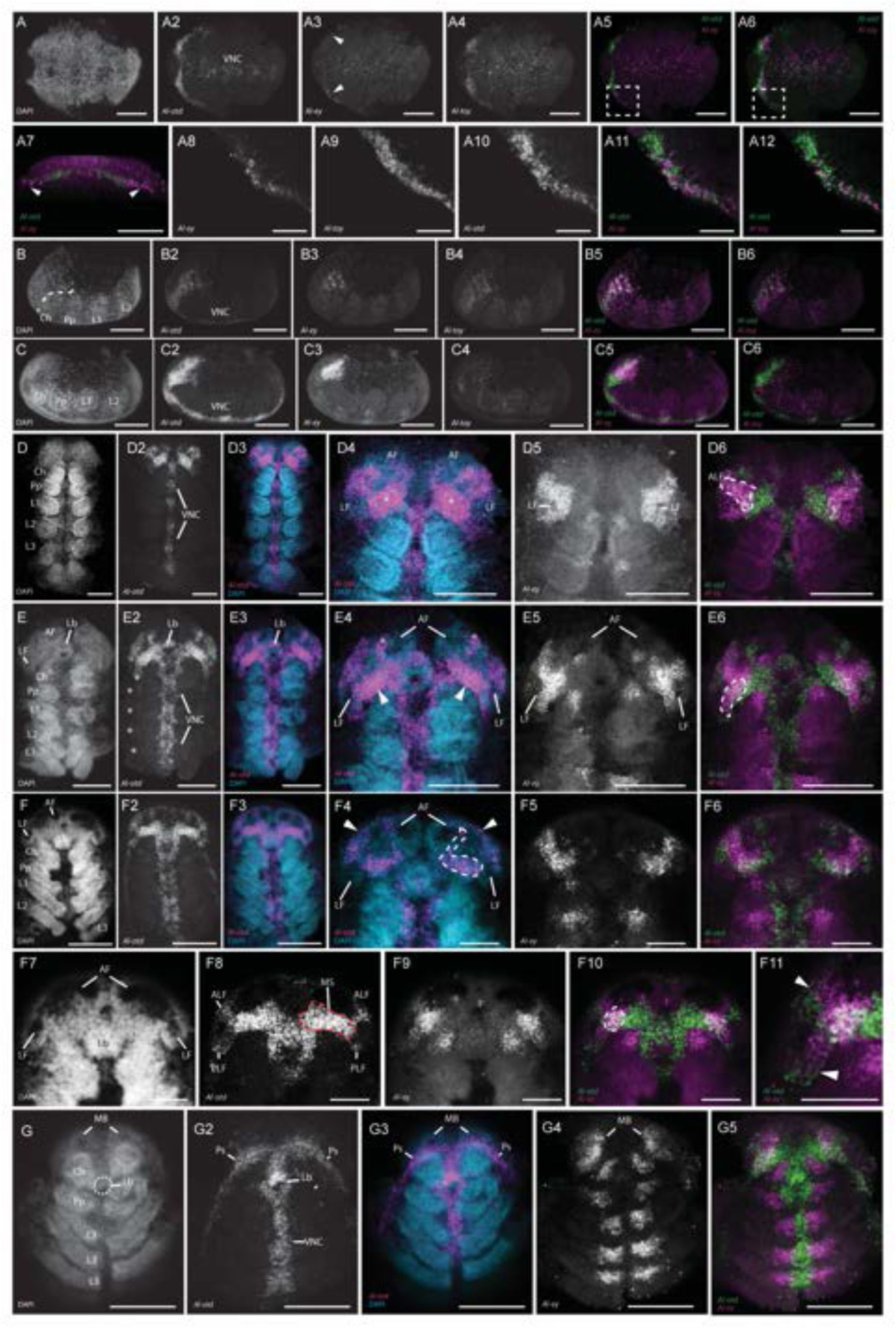
The co-expression of the *Al-Pax6* orthologs with the head-patterning gene *orthodenticle*. **A-A6** *Al- Pax6* co-expression with *Al-orthodenticle* (*Al-otd*) in an early, pre-segmental embryo. The ventral portion of the embryo is shown. **A** DAPI counterstain of this embryo. **A2** *Al-otd* expression in this embryo in a continuous stripe of expression in the embryonic anterior (left of the page). Also, *Al-otd* is weakly expressed in cells of the incipient ventral nerve cord (VNC). **A3** *Al-ey* expression is restricted to two, lateral domains in the anterior rim of the embryo. **A4** *Al-toy* is also expressed in two, paired domains in the anterior of the embryo. **A5** Merged image of the *Al-otd* (green) and *Al-ey* (magenta) channels. **A6** Merged image of the *Al-otd* (green) and *Al-toy* (magenta) channels. Dotted boxes in these images represent the fields of view shown in A8-A12. **A7** Three-dimensional projection of the same embryo, rotated to show the frontal-most expression of *Al-otd* (green) and *Al-ey* (magenta; arrowheads) in the anterior. The embryo is oriented with the ventral towards the top of the page. **A8-A10** Single channel confocal images of *Al-ey*, *Al-toy*, and *Al-otd*, respectively, in the region of the dotted boxes outlined in A5 and A6. **All** Confocal slice of the region of the embryo outlined in the dotted box in A5. **Al2** Confocal slice of the region of the embryo outlined in the dotted box in A6. A11-A12 show that both *Al-Pax6* orthologs are co-expressed with *Al-otd* in this region. **B-B6** *Al-Pax6* and *Al-otd* expression in an early, three prosomal segmented staged embryo. **B** DAPI nuclear counterstain. **B2** *Al-otd* expression in the developing head. Expression also persists in the VNC. **B3** *Al-ey* expression. **B4** *Al-toy* expression. **B5** Merged confocal channels of *Al-otd* (green) and *Al-ey* (magenta) expression. **B6** Merged confocal channels of *Al-otd* (green) and *Al-toy* (magenta) expression. Note that this is the same embryo shown in Fig. 3B-B4. **C-C6** *Al-Pax6* and *Al-otd* expression in an early, limb-bud stage embryo. **C** DAPI nuclear counterstain. **C2** *Al-otd* expression. **C3** *Al-ey* expression. **C4** *Al-toy* expression. **C5** Merged confocal channels of *Al-otd* (green) and *Al-ey* (magenta) expression. **C6** Merged confocal channels of *Al-otd* (green) and *Al-toy* (magenta) expression. **D-D6** *Al-ey* and *Al-otd* expression in an embryo at approximately stage BDS-2. **D** DAPI nuclear counterstain. **D2** *Al-otd* expression. **D3** Merged confocal channels of *Al-otd* (magenta) and the DAPI counterstain (cyan). **D4** The same merged image shown in D3, zoomed in on the developing pre-cheliceral region. Asterisks demark the “blocks” of *Al-otd* expression that surrounds the incipient stomodaeum. LF= the sites of the developing lateral furrows. **D5** Image showing *Al-ey* expression in this embryo. **D6** The same embryo, however the channels showing *Al-otd* expression (green) and *Al-ey* expression (magenta) have been merged. The dotted line outlines the co- expression of these genes in the anterior lateral furrows. **E-E6** *Al-ey* and *Al-otd* expression in an embryo at approximately stage BDS-4. **E** DAPI nuclear counterstain. **E2** *Al-otd* expression. The asterisks demark the horizontal lines of expression at the proximal-most boundary of each prosomal appendage. **E3** Merged confocal channels of *Al-otd* (magenta) and the DAPI counterstain (cyan). **E4** The same merged image shown in E3, zoomed in on the developing pre-cheliceral region. Arrowheads mark the “blocks” of *Al-otd* expression. Asterisks mark expression in the lateral margins of the anterior furrows. **E5** Image showing *Al-ey* expression in this embryo. **E6** The same embryo, however the channels showing *Al-otd* expression (green) and *Al-ey* expression (magenta) have been merged. The dotted line outlines the co- expression of these genes in the ventral-most portion of the lateral furrows. **F-F11** *Al-ey* and *Al-otd* expression in an embryo at approximately stage BDS-5. **F** DAPI nuclear counterstain. **F2** *Al-otd* expression. **F3** Merged confocal channels of *Al-otd* (magenta) and the DAPI counterstain (cyan). **F4-F6** Confocal slices of the same embryo, however the slices were taken more dorsally in the embryo. **F4** The same merged image shown in F3, zoomed in on the developing pre-cheliceral region. Arrowheads point to *Al-otd* expression in the margins of the head lobes in the presumptive non-neural ectoderm. *Al-otd* expression in one the medial subdivisions is outlined with a dotted line. **E5** Image showing *Al-ey* expression in this embryo. **F6** The same embryo, however the channels showing *Al-otd* expression (green) and *Al-ey* expression (magenta) have been merged. **F7-11** The same embryo, however the images were taken at a more “surface-level”, or ventral position of the embryo in comparison to F4-F6. **F7** DAPI counterstain at this position. **F8** *Al-otd* expression at this level, showing its expression relative to the anterior lateral furrows (ALF) and the posterior lateral furrows (PLF). MS=expression in the medial subdivision. The right-most MS is outlined in a red, dotted line. **F9** *Al-ey* expression at this position. **F10** Merged image showing *Al-otd* expression (green) and *Al-ey* expression (magenta). Note the co- expression of these genes in the lateral component of the medial subdivisions. The left one is outlined in a dotted line. **F11** The same image shown in F10, however it is zoomed into one of the lateral furrows. Arrowheads point to the boundaries between *Al-ey* and *Al-otd* expression. **G-G5** *Al-ey* and *Al-otd* expression in an embryo at approximately stage BDS-7. **G** DAPI nuclear counterstain. The dotted outline shows the position of the labrum which has been obscured by the movements of the pre-cheliceral region at this stage. **G2** *Al-otd* expression in this embryo, showing its expression in the lateral margins of the two halves of the prosomal shield (PS). **G3** Merged confocal channels of *Al-otd* (magenta) and the DAPI counterstain (cyan). **G4** Image showing *Al-ey* expression in this embryo. **G5** The same embryo, however the channels showing *Al-otd* expression (green) and *Al-ey* expression (magenta) have been merged. Embryos shown in B-C6 are oriented with their anterior pointing towards the left of the page, and their dorsal regions towards the top of the page. All subsequent images are oriented so that the anterior of the embryos are directed towards the top of the page. All scale bars represent 50 µm, except as follows: A8- A12, 10 µm; F4-F11, 20 µm. All other abbreviations are the same as in other figures.

**Fig. 7.**
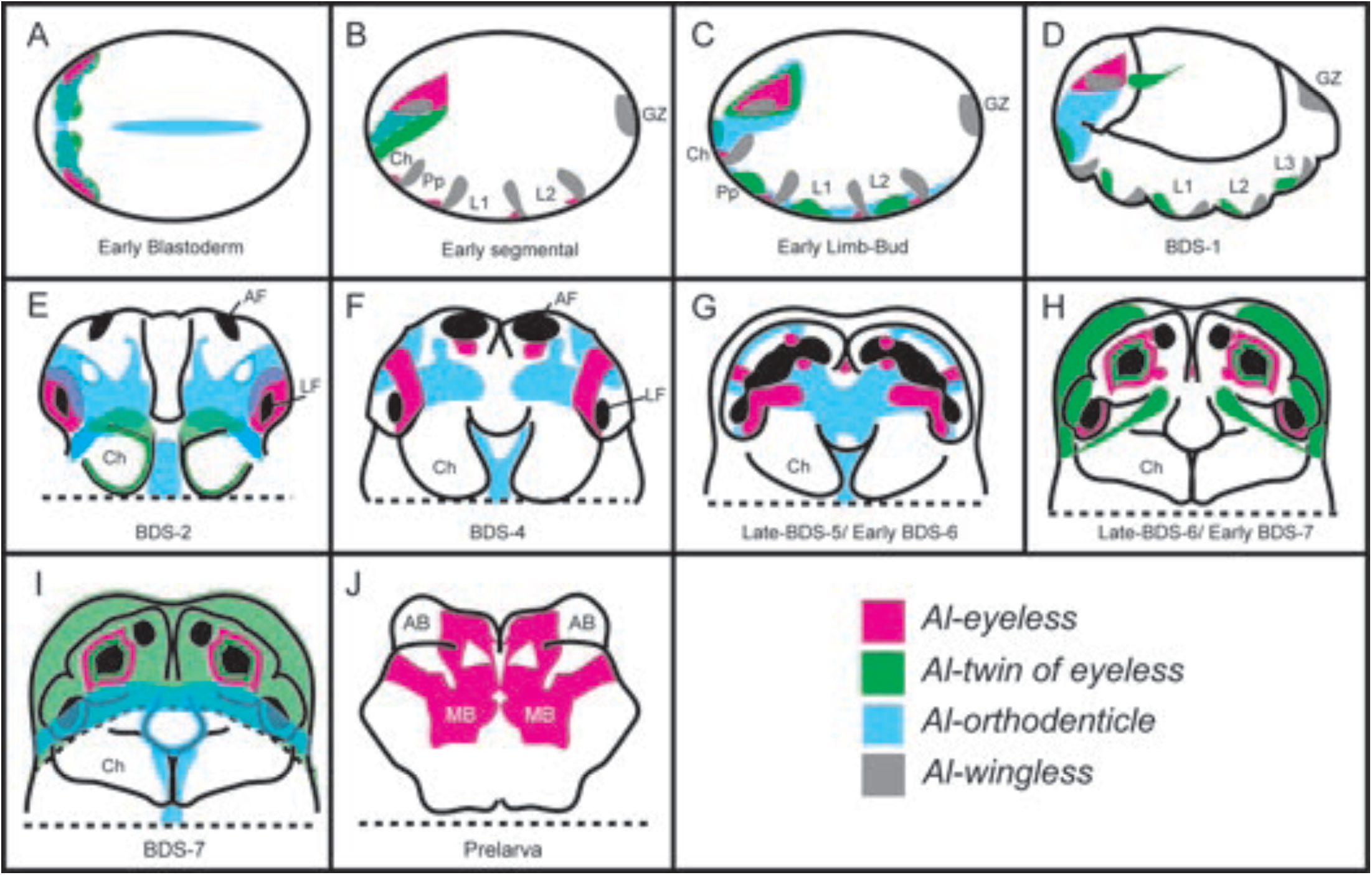
Summary drawings showing the relative expression patters of the *Al-Pax6* orthologs and *Al-otd* throughout the development of the pre-cheliceral region. See text for details.

In a subsequent germ-band stage, at which the first three prosomal segments had been delineated (*i.e.*, the cheliceral, pedipalpal and first two walking leg segments), *Al-otd* expression was retained in the incipient ventral nerve cord, as well as the pre-cheliceral region (Fig. 6B2). *Al-ey* was co-expressed with *Al-otd* at this stage in its lateral, triangular domains of expression (Fig 6B3). *Al-toy* was also co-expressed with *Al-otd* at this stage, however *Al-toy*’s expression domain was much broader and fully encompassed the domain of *Al-otd* where it extended more posteriorly in the pre-cheliceral region (Fig. 6B4 and B6).

As show in Fig. 3, *Al-toy* expression dramatically decreases in the pre-cheliceral region following its initial expression. Approximately at the stage when this decrease begins, and when the prosomal limb buds became more distinct, *Al-otd* retained its expression domain in the pre-cheliceral region where it was still co-expressed with *Al-ey* (Fig. 6C2, C3, and C5). However, *Al-toy* was not observed to be co-expressed with *Al-otd* at this stage (Fig. 6C6).

We argue that these early expression patterns are likely indicative of a role for both *Pax6* genes and *Al-otd* acting together to specify the neural field of the incipient protocerebrum. Evidence for this includes the fact that eye development in all chelicerates studied occurs after the protocerebral region has been specified, and also that the eye precursor cells form in the non-neural cells of the peripheral head lobes. Thus, the co-expression of *Pax6* genes and *otd*, as seen in spiders, is likely independent of their expression in the incipient eyes. Therefore, our results are more consistent with a role of *Pax6* genes in specifying the protocerebrum, and that this role was inherited from the last common ancestor of spiders and mites.

### Late expression of *Al-otd* suggests the ancestral role of orthodenticle in arachnids

It was also shown that both the duplicated spider *otd* paralogs, as well as the singleton *otd* ortholog in the opilionid *P. opilio*, are expressed in the developing brain and eyes at developmental stages following the establishment of the head [28,30,32,34]. We thus utilized these data to ask how the spider *otd* paralogs were potentially subfunctionalized, or neo-functionalized, in the lineage leading to spiders.

We began these observations at BDS-2, when the morphology of the pre-cheliceral region becomes more complex (note that *Al-otd* is expressed broadly at BDS-1 in a manner similar to Fig. 4C5; not shown). At this stage, we detected *Al-otd* expression in the pre-cheliceral region as well as its retention in the developing ventral nerve cord (Fig. 6D-D3). Within the pre-cheliceral region, *Al-otd* was expressed in the anterior and posterior margins of the nascent lateral furrows. Note that *otd2* is also expressed in the lateral furrows of the spider *P. tepidariorum* [30,31], as is *otd* in the opilionid *P. opilio* [32]. *Al-otd* was also expressed in connected domains that span the central to the lateral pre-cheliceral region (Fig. 4D4). We also detected *Al-otd* expression in two “blocks” of cells surrounding the site of the future stomodaeum (Fig. 4D4, asterisks) in a similar manner to *otd* in *P. opilio* [32] and *otd2* in spiders [31].We likewise co-stained these embryos with *Al-ey* expression, and found that *Al-otd* and *Al-ey* expression overlap in the anterior of the lateral furrows (note that *Al-toy* expression is absent in the pre- cheliceral region at this stage). However, they do not overlap at the posterior portion of the lateral furrows (Fig. 6D6, the left anterior portion of the lateral furrow, ALF, is outlined).

At approximately BDS-4, *Al-otd* expression was still present in the developing ventral nerve cord (Fig. 6E-E2). Faint *Al-otd* expression was also detected in horizontal lines of expression at the proximal-most boundary of each prosomal appendage (Fig. 6E2, asterisks). This appendicular expression may be homologous to that of *Pt-otd2* expression at later stages (*i.e.*, stages 12 and 13; see Figs. 11H-I in [30]) and also to *otd* expression in opilionid *P. opilio* ( [32], their Fig. S2). Within the pre-cheliceral region, *Al- otd* expression was detected in the developing labrum (Fig. 6E-E3), in a similar manner to spider *otd2* orthologs [28,31] and to the expression of *otd* in *P. opilio* [32]. *Al-otd* expression was subsequently retained in the “blocks” of cells surrounding the stomodaeum (Fig. 6E4, arrowheads), and was also detected in the lateral margins of the anterior furrows (Fig. 6E4, asterisks). By co-staining for *Al-ey* expression, we were able to detect its co-expression with *Al-otd* expression in the ventral-most portion of the lateral furrows (Fig. 6E6, dotted outline).

At BDS-5, the expression of *Al-otd* in the ventral nerve cord remained, and the proximal appendicular expression domains became more pronounced. Furthermore, each of these domains appeared to combine to become continuous with one another on their respective half of the embryo (Fig. 4F-F3) similar to *otd* expression at stages 12-15 in *P. opilio* [32]. Also in a similar manner to the spider *otd2* orthologs [28,31], and the *P. opilio otd* ortholog [32], we additionally detected *Al-otd* expression in the margins of the head lobes in the presumptive non-neural ectoderm (Fig 6F4, arrowheads). Also, *Al- otd* expression surrounded the periphery of the fused labrum (Fig. 6F-F3).

Recall that at this stage, the medial subdivisions send out “extensions” of cells that will separate the anterior furrows from the nascent mushroom bodies (Fig. 1E-E2). We detected *Al-otd* expression in these medial subdivision extensions (Fig. 6F4; the right-most medial subdivision’s extension is outlined). Because of their proximity to the persisting “blocks” of *Al-otd* expression surrounding the labrum at this stage, we take the “blocks” of *Al-otd* expression in BDS-2 and BDS-4 to be the medial subdivisions. We also detected the co-expression of *Al-ey* and *Al-otd* within a population of cells in the lateral halves of these medial subdivisions (Fig. 6F8-11; one of these populations is outlined in F10). These patterns are similar to those of spider *otd.2* expression [31] and to *P. opilio otd* expression [32], suggesting a high degree of conservation of *otd* expression in these tissues.

*Al-otd* expression was then detected at late BDS-5/early BDS-6 in the ventral-most half of the lateral furrows, and its expression was the highest in their anterior and posterior poles. Because *Al-ey* at this stage is also expressed in the ventral-most half of the lateral furrows (see Fig. 4F-F3), we asked to what extent *Al-otd* and *Al-ey* are co-expressed in these “open” lateral furrows. By co-detecting *Al-ey* with *Al-otd*, we found that their expression is mutually exclusive in the anterior lateral furrows, with *Al-otd* expression located more “inwardly” in the lateral furrow that *Al-ey* expression (Fig 6F10-11; arrowheads in F11 point to the boundaries between *Al-ey* and *Al-otd* expression). These expression patterns are interesting, as they may point to a coordinate-type system to establish polarity, or a mechanism of regionalization, in the lateral furrows. Lastly, both *Al-ey* and *Al-otd* were co-expressed in the ventral-lateral cells that are adjacent to the lateral furrows (Fig 6F10-11; these cells are outlined in the left side of the embryo in F10).

At approximately BDS-7, when the prosomal shield halves had migrated and fused, we observed *Al-otd* expression in the margins of the fused prosomal shield. This expression domain was continuous to the lateral appendicular expression domains of the previous stages (Fig. 6G-G3). A similar colorimetric staining pattern was observed for *otd2* orthologs in the spider species studied in [31], however the authors described these as the results of artefactual cuticle staining. Interestingly, we did not observe similar staining to that shown in Fig. 6G-G5 in our control experiments, nor in any other HCR experiments of other genes. Furthermore, *P. opilio otd* is expressed in a similar pattern at stages 12-15 [32].

Therefore, we take this expression pattern to be a conserved mode of *otd* expression amongst arachnids.

*Al-otd* expression was also present in the ventral nerve cord and the periphery of the labrum at this stage (Fig. 6G-G3). Within the pre-cheliceral region, we did not observe *Al-otd* in any of the tissues in which it was expressed in the previous stages. This was shown via the co-staining of *Al-otd* with *Al-ey*, which is expressed largely in the mushroom bodies at this stage (Fig. 6G4-G5). We did not observe expression of *Al-otd* at any subsequent pre-larval or larval stages (not shown).

In terms of eye development, one *P. tepidariorum otd* ortholog, *Pt-otd2*, is expressed late in development in the anterior median eyes [30,31]. In the spider *C. salei*, both of its *otd* paralogs are expressed in tissues associated with the eyes. Specifically, *Cs-otxa* (=*otd1*) is expressed in the vesicles of the posterior-lateral eyes, whereas *Cs-otxb* (*=otd2*) is expressed in all sets of lateral eyes as well as the posterior median eye vesicles [28]. In a recent, comprehensive study of diverse spider taxa, it was shown that in all of the spider species studied, *otd2* orthologs are expressed exclusively in the anterior-median eyes, with the notable exception of a lack of any eye *otd* expression in *Pholcus phalangioides*[31]. Additionally, a recent study on *P. opilio* revealed that its *otd* ortholog is expressed in all of its eye primordia [32]. We did not observe similar expression patterns of *Al-otd* in the aforementioned HCR experiments, further confirming the absence of vestiges of eyes in *A. longisetosus* during embryonic development. In summation, our results, coupled with those observed in *P. opilio* [32], suggest that *Al-otd* expression is most similar to spider *otd2* ortholog expression in the pre-cheliceral region [31]

## Discussion

### The morphogenesis of the arachnid head and brain in light of *A. longisetosus*

Modern studies into the development of the arachnid head have largely focused on the spiders *P. tepidariorum* (e.g., [51] and *C. salei* (e.g.,[50,54]). These studies, in conjunction with a recent studies on the development of the opilionid *P. opilio* [32,55], have revealed potential synapomorphic features of arachnid head development. These synapomorphies include the appearance of the anterior and lateral furrows, followed by the migration of the non-neural prosomal shield over the pre-cheliceral region.

Despite the conservation of these features *in A. longisetosus*, our data show major morphogenetic divergences between mites and the aforementioned arachnids in the embryonic pre-cheliceral region.

First, we observed differences in the timing of the appearance of the lateral and anterior furrows.

In the aforementioned spider exemplars, the lateral furrows form first, followed by the anterior furrows [50,51]. This may also be the order of appearance in *A. longisetosus*, however with our methods, we were only able to visualize the appearance of both pairs of furrows simultaneously at BDS-2. In *P. opilio*, it is also unclear as in which order these furrows appear, as they also seem to appear simultaneously (see Fig. 8C in [8]).

Another point of divergence between spiders and *A. longisetosus* can be seen in the later morphogenesis of the anterior and lateral furrows. In spiders, the division of the anterior furrows into the mushroom and arcuate bodies occurs through the migration of the medial subdivisions expanding into the anterior furrows, which results in the compartmentalization of the arcuate and mushroom bodies [50,51,54]. Our observations suggest that this aspect of brain morphogenesis is conserved between mites and spiders, with one major exception. In the aforementioned spider species, the anterior furrows do not appear to make continuous grooves with the lateral furrows at any stage. In *A. longisetosus*, however, the lateral and anterior furrows become continuous with one another after their initial appearances as distinct structures (see Fig. 1). These continuous grooves are then subsequently subdivided to form the arcuate bodies, mushroom bodies, and the optic vesicles, with the arcuate and mushroom bodies delimited by extensions of the medial subdivisions. A secondary, and as of yet unnamed, group of cells forms later to separate the posterior of the mushroom bodies from the anterior of the lateral furrows.

One last point of divergence was seen in the development of the lateral furrows. In spiders, the lateral furrows are further subdivided through the expansion of another grouping of paired elevated neural tissues called the lateral subdivisions. These lateral subdivisions subdivide the lateral furrows into the lateral and medial optic vesicles [28–32,54]. Our observations of *A. longisetosus* brain compartmentalization did not show evidence of any lateral subdivisions. Interestingly, it has been proposed that it is these lateral vesicles give rise to the optic ganglia of the lateral eyes of spiders [54]. It is therefore tempting to attribute the absence of the lateral subdivisions in *A. longisetosus* to their absence of eyes. However, to test this hypothesis, the compartmentalization of the brains of acariform mites that have retained their eyes needs to be studied.

Taken together, our results suggest that presence of paired anterior and lateral furrows, as well as the migration of the prosomal shield are conserved aspects of arachnid brain/pre-cheliceral region development. Despite this, the subsequent subdivisions of these furrows may be lineage-specific. An alternative explanation for our observations could be that the mode of brain compartmentalization in *A. longisetosus* is highly derived within Acariformes. To clarify this, more studies into the brain development of members of this hyper-diverse clade are needed. Also, given that *A. longisetosus* lacks eyes, the presence of the lateral furrows that give rise to the optic vesicles needs an explanation. The simplest explanation is that these compartments of the brain do not only give rise to the optic neuropils as they do in spiders, and they therefore may contribute to other important components of the brain. Alternatively, these lateral furrows could be vestiges of the optic neuropils of true acariform eyes. Cellular lineage tracing experiments, which are not yet available for this species, are needed to further explore these hypotheses.

### *Pax6* genes and arachnid eyes

The ancestral role of *Pax6* gene expression in the development of chelicerate eyes has thus far puzzled researchers (reviewed in [33]). An early expression study of a single *toy* ortholog in the horseshoe crab *Limulus polyphemus* showed no clear expression in any of its eye anlagen [27]. Since this study, it has been revealed that *L. polyphemus* has at least five *Pax6* orthologs, two of which that are likely *toy* orthologs and three of which that are likely *eyeless* orthologs [13,26]. It may still be shown that some of these orthologs are indeed expressed in the developing horseshoe crab eyes. However, recent data from spiders have shown no expression of any *Pax6* gene in any of its eye anlagen [29–31,34,80], except for the expression of *eyeless* in the anterior-median eyes of *C. salei* [28]. Furthermore, a recent study into the RDGN genes of *P. opilio* showed that both *Pax6* genes are expressed in the developing median eyes[32]. These observations have thus far neither supported nor falsified the hypothesis that *Pax6* genes did not play an ancestral role the development of chelicerate eyes and may instead suggest that a lack of *Pax6* expression in spider eyes may be specific to that clade.

An alternative explanation has been proposed, which posits that the expression of *Pax6* genes observed during early spider development act to specify the fates of eye photoreceptor cells prior to their incorporation into the eyes during subsequent morphogenetic events. Using our eyeless mite, as well as its expression patterns of the early head-patterning gene *orthodenticle*, we tested this hypothesis. Our data show similarities to the early expression patterns of *eyeless* and *toy* of spiders in the pre-cheliceral region. We therefore argue that these data are more consistent with an ancestral role of *Pax6* genes acting with *otd* to specify the neural cells of the protocerebrum in arachnids. It will be interesting to see if this holds true throughout in the remaining arachnid groups, including other non-spider members of Arachnopulmonata, as well as other members of Acariformes.

### *Al-ey* expression in comparison to other arachnids

Our data show both shared and derived modes of *eyeless* expression in arachnids. We found that early *Al-ey* expression is similar to that of spiders, specifically in comparison to early *P. tepidariorum eyeless* expression [30,31,34]. In both taxa, *eyeless* is expressed early in an anterior domain in the pre-cheliceral region. Furthermore, our data highlights other conserved aspects of arachnid *eyeless* expression. We observed its expression in the nascent mushroom bodies and the optic vesicles. *eyeless* expression was similarly observed in spiders [28,30,31,34], and may also be expressed in these structures in *P. opilio* [32].

By evaluating *eyeless* expression patterns reported for other taxa, we conclude that in spiders, in an opilionid and in *A. longisetosus* (Fig. 2), *eyeless* is subsequently expressed in cells associated with the lateral furrows. Because the lateral furrows likely develop into the optic vesicles in spiders [54], the usage of these optic vesicles in arachnids lacking eyes should be a key focus of study to further our understanding of both the arachnid brain, its development, and the function of *eyeless* in patterning these structures. Our results cannot falsify the hypothesis that our observed *Pax6* expression patterns are vestigial, *i.e.*, relictual features of eye development. We are currently exploring methods to abrogate gene expression in *A. longisetosus*, however we are thus far limited to gene expression surveys. Once methods to test for gene functions are in place, we plan to knock down both *Pax6* genes to test this hypothesis directly. If *Pax6* gene expression is indeed vestigial in *A. longisetosus*, we would expect to see no morphogenetic anomalies in *Pax6*-depleted embryos. Related to this, we cannot thus far falsify the hypothesis that the lateral furrows/optic vesicles are themselves vestigial. Because we do not yet know if, or exactly what, other non-visual roles of the optic vesicles may be, more functional neural studies into these compartments of the arachnid brain are necessary before making this conclusion.

### *Al-toy* expression in comparison to other arachnids

Our data also suggests both conserved and derived aspects of *toy* expression amongst arachnids. To date, early expression data for *toy* in arachnids is limited to the spider *P. tepidariorum* [34]. In both this spider and in *A. longisetosus*, *toy* is expressed in a broad anterior domain in the pre-cheliceral region. Following this pattern, *Al-toy* expression deviates dramatically from its ortholog’s expression in other arachnids. For instance, *Al-toy* is expressed in the developing prosomal appendages, in a manner not seen in other studied arachnids. Also, *Al-toy* expression disappears from the pre-cheliceral region at intermediate stages of development. This is in stark contrast to opilionids and spiders, in which *toy* expression persists in the pre-cheliceral region at comparable stages [31,32]. We also observed ubiquitous expression of *Al-toy* in the migrating prosomal shield at late stages of development. In the opilionid *P. opilio*, both *Pax6* orthologs appear to be ubiquitously expressed in this structure at later stages [32]. In spiders, neither *Pax6* ortholog appears to be expressed ubiquitously in the prosomal shield [31]. It is tempting to conclude that these deviations in the use of *toy* in *A. longisetosus* is due to their absence of eyes. However, since *toy* does not contribute to eyes in any studied arachnid, the differential use of *toy* in *A. longisetosus* necessitates future functional studies.

### Eyes in Acari: future directions

Members of Acari (mites and ticks) display a wide degree of morphological diversity, owing to their occupancy of numerous ecological niches [81,82]. The monophyly of Acari is currently still contested (e.g., [7,83,84]), however it is generally agreed that Acari is comprised of two internally-monophyletic groups, the Parasitiformes (*e.g.,* ticks) and Acariformes (mites). Within Acariformes, the number, position, and the types of eyes present are extremely diverse, also owing to their ecological diversity (see [81,82,85]for review). Examples of this diversity include the retention of both lateral and median eyes, with varying numbers of each (*e.g.*, the mite *Heterochthonius gibbus* has one median eye and a pair of lateral eyes [86]), or the parallel loss of all eyes independently in a number of acariform groups (reviewed in [81,82]). Notwithstanding, it has been hypothesized that the ground plan for acariform mites is the presence of two median eyes and two pairs of lateral eyes, a condition that was inferred in an early study [87]. However, it has been cautioned that determining the pleisiomorphic condition of acariform eyes is a complex problem considering the morphological disparity between acariform sub-clades [85].

Nonetheless, given the wide degree of eye diversity in mites, coupled with the emergence of new developmental data on the expression and function of RDGN genes in arachnids, mites are a clade of extreme interest in terms of exploring the developmental evolution of arachnid eyes.

## Conclusions

Eye loss is extensive across mite species, however paradoxically, the eyeless mite *A. longisetosus* retains two *Pax6* paralogs in its genome. These observations, coupled with recent data suggesting that there is no role for these genes in the development of spider and horseshoe crab eyes, allowed us to test the hypothesis that ancestrally chelicerates did not use *Pax6* genes to pattern their eyes. By following the expression of genes canonically associated with arthropod eye development, we found support for the hypothesis that *A. longisetosus* does not develop vestigial eye tissues. Following this, we showed that both early and late *Al-ey* and *Al-toy* expression patterns are highly similar to those of studied chelicerate taxa that do have eyes. An alternative hypothesis, *i.e.,* that chelicerates do use *Pax6* genes to pattern early photoreceptor cells prior to their incorporation into the eyes, is not supported by our findings. We reasoned that, if this hypothesis were supported, we would have observed either an absence of early *Pax6* gene expression, and/or highly derived *Pax6* gene expression in comparison to other studied chelicerates. Our findings do not support this hypothesis but rather suggest that these genes were used in the development of the ancestral chelicerate brain/central nervous system. Our results also support the hypothesis that ancestrally, *Pax6* genes worked with *orthodenticle* to specify the neural cells of the protocerebrum independent of *orthodenticle’s* role in eye specification. Lastly, our results suggest that one *Pax6* ortholog, *eyeless*, was likely used ancestrally in chelicerates to specify the paired structures of the chelicerate brain called the “optic vesicles” as well as the mushroom bodies.

Studies into the function of *Pax6* genes in model, and emerging model, chelicerates are needed to further test these hypotheses. The potential roles of *Pax6* genes observed in the median eyes of the spider *C. salei* [28], and the median eyes of the opilionid *P. opilio* [32] , also need further investigation to test the hypotheses that these patterns are due to lineage-specific adaptations, or due to developmental systems drift. Furthermore, studies into the use of *Pax6* genes are needed in members of Pycnogonida (sea spiders), as they are the likely sister-group to the remaining chelicerate taxa. If *Pax6* gene expression were also absent in their embryonic eyes, this would further support the hypothesis that *Pax6* genes played no role in the ancestral development of chelicerate eyes.

## Acknowledgements

This work was produced by IJ as part of her undergraduate Honors Thesis. We would therefore like to specially thank her thesis committee members, Joseph Leese, Dia Beachboard, and Daniel Proud. We would also like to thank the members of the Barnett Lab, as well as Prashant Sharma, Michael Layden, and Richard Thomas for their helpful comments on this manuscript.

## Declarations

### Ethics approval and consent to participate

Not applicable.

### Consent for publication

Not applicable.

### Availability of data and materials

All original confocal images are available upon request to AAB. The transcriptome sequences are publicly available from [38]. All probe sequences for the hybridization chain reactions are available in Supplementary Tables S3-S17.

### Competing interests

The authors declare no competing interests.

### Funding

Funding for this work was provided by the DeSales University Berg Endowment, and funds from the DeSales University Center for Teaching Excellence Innovation Grant awarded to AAB.

### Author contributions

IJ and AAB developed the study design and experiments. IJ and AAB both performed the wet-lab experiments, performed the confocal microscopy, as well as the bioinformatic analyses. Both authors wrote the original draft. All authors edited and approved the final version.

---

Fig. S1 The architecture and phylogenetic interrogation of *A. longisetosus eyeless* and *twin of eyeless* sequences. (**A**) Schematics showing the deduced genomic architecture of *Al-eyeless*, as well as the architecture of its spliced transcripts. In the genomic schema, the exons are represented as grey boxes, and the dotted lines represent splicing patterns. The dotted lines on the top of the map correspond to splicing patterns of the *Al-ey.1* isoform. The blue box below exons 8 and 9 represents exon 8.5, which is used in the Al-ey.2 isoform. Below the genomic schema are schematics representing the architecture of *Al-ey.1* and *Al-ey.2*. These show how each exon corresponds to the resulting mRNA landmarks, including the untranslated regions (UTRs), the paired domains (PD), the octapeptide-like domains (OP-Like), and the homeodomains (HD). (**B**) Schematics showing the deduced genomic architecture of *Al-toy*, as well as the architecture of its spliced transcript. All annotation styles and abbreviations are the same as in **A**. (**C**) Alignments of the deduced amino acid sequence of the paired domains of both Al-Ey and Al-Toy with selected arthropod Pax6 orthologs. The linker region bisects the paired domain into PAI and RED sub- domains. The linker region of Al-Ey, as well as that of other arthropod orthologs, contains a distinct arginine (R) at position 64 (highlighted in red). The linker region of Al-Toy and other arthropod Toy orthologs contains the diagnostic lysine (K) at this position (also highlighted in red). (**D**) The resulting tree from a maximum-likelihood phylogenetic analysis of selected arthropod Ey, Toy, and Pax2/5 proteins. The clade with green-highlighted branches represents a monophyletic Toy clade, and the clade with blue branches represents the monophyletic Ey clade. *A. longisetosus* Pax sequences are denoted with red labels. The node colors correspond to approximate likelihood ratio test (aLRT) scores (see legend). All labels have their GenBank accession numbers in parentheses.

Fig. S2 Phylogenetic analysis of putative *A. longisetosus* Arrestin protein orthologs. The tree is transformed as a cladogram to improve visualization. The red clade highlights Kurtz orthologs, the green clade highlights Arrestin-1 orthologs, and the blue clade highlights Arrestin-2 orthologs. The node colors correspond to approximate likelihood ratio test (aLRT) scores (see legend). All labels have their GenBank accession numbers in parentheses.

## Notes

### Competing Interest Statement

The authors have declared no competing interest.

