## Supplemental Tables for "The Expression of *Pax6* Genes in an Eyeless Arachnid Suggests Their Ancestral Role in Arachnid Head Development"

**Table S1:** Statistics for the phylogenetic trees. See text for details.

| Ortholog Search | SMS Substitution Model, BIC | SMS Decoration, BIC | Tree Log-Likelihood |
|---|---|---|---|
| ***eyeless (ey), twin of eyeless (toy)*** | Q.insect, 63856.57 | R+F, 131422.34 | -63856.57 |
| ***beta-arrestin*** | Q.insect, 82423.62 | R+F, 82423.62 | -414303.87 |

**Table S2:** The transcript IDs and amplifier sequences used to construct each HCR probe. Note that for *Al-toy,* two probe sets were constructed.

| Gene | Transcript | HCR Amplifier |
|---|---|---|
| ***Al-arrestin-1*** | TRINITY_GG_3318_c59_g1_i3 | B3 |
| ***Al-arrestin-2*** | TRINITY_GG_4713_c203_g1_i1 | B1 |
| ***Al-ato*** | TRINITY_GG_4863_c1996_g1_i1 | B2 |
| ***Al-dac*** | TRINITY_GG_5120_c350_g1_i2 | B2 |
| ***Al-ey*** | TRINITY_GG_2648_c164_g1_i2 | B1 |
| ***Al-eya*** | TRINITY_GG_4744_c173_g1_i10 | B2 |
| ***Al-krz*** | TRINITY_GG_5120_c51_g1_i7 | B2 |
| ***Al-myoIII*** | TRINITY_GG_3318_c34_g1_i4 | B1 |
| ***Al-otd*** | TRINITY_GG_5990_c97_g1_i2 | B3 |
| ***Al-peropsin*** | TRINITY_GG_4858_c50_g1_i1 | B3 |
| ***Al-rhodopsin*** | TRINITY_GG_6262_c55_g1_i2 | B3 |
| ***Al-Six3*** | TRINITY_GG_5990_c859_g1_i1 | B2 |
| ***Al-so*** | TRINITY_GG_5990_c254_g1_i5 | B3 |
| ***Al-toy*** | TRINITY_GG_5245_c530_g1_i1 | B1, B2 |
| ***Al-wg*** | TRINITY_GG_4863_c585_g1_i7 | B2 |

**Table S3:** Probe pairs designed for *Al-arrestin-1* HCRs (B3 initiator)

| Pair | Initiator | Spacer | Hybridzation | Hybridzation | Spacer | Initiator |
|---|---|---|---|---|---|---|
| **1** | GTCCCTGCCTCTATATCT | TT | TTTAATAGACTTGAGTTAGCATTGC | AAGCTATGAAAAAGTTGTCGGACGA | TT | CCACTCAACTTTAACCCG |
| **2** | GTCCCTGCCTCTATATCT | TT | TCACTAGTAAGGGAAGAGTTACCTC | ACATTATTGGAGAGGGTGTCGTACG | TT | CCACTCAACTTTAACCCG |
| **3** | GTCCCTGCCTCTATATCT | TT | TGGAAAAGTGATGTGTAGGGTTAGA | AAAAATAGGAATGTCTTGTATGACC | TT | CCACTCAACTTTAACCCG |
| **4** | GTCCCTGCCTCTATATCT | TT | TAGTGATGGAACATTACAGCGACGA | AATGATAGCGACATATCCCGTTCGG | TT | CCACTCAACTTTAACCCG |
| **5** | GTCCCTGCCTCTATATCT | TT | ATCCGTTCAGCTCTTCAACACTGTC | CTACTTAAGTATGATATTTTCACTG | TT | CCACTCAACTTTAACCCG |
| **6** | GTCCCTGCCTCTATATCT | TT | TCACCAAATTTGAACGGAAAAGTCC | TTGGTTTATGTCTCAATTGTGATGC | TT | CCACTCAACTTTAACCCG |
| **7** | GTCCCTGCCTCTATATCT | TT | AGTATAGAGTCAGGTATAGCGTAGA | AGTGACAGACAGCCTAAAGAGTAAA | TT | CCACTCAACTTTAACCCG |
| **8** | GTCCCTGCCTCTATATCT | TT | ACAGAGGCATTGGTGGAAAGTTTGT | CATGTCAGCTACCAGTTACCACCGA | TT | CCACTCAACTTTAACCCG |
| **9** | GTCCCTGCCTCTATATCT | TT | CAACCGTTTACAGCGCTATTAAAAC | TCAGGATTTTTACAGCAAAAGCTCA | TT | CCACTCAACTTTAACCCG |
| **10** | GTCCCTGCCTCTATATCT | TT | CTCATCCAGCCCTTAATGGTTCGAC | CCTTTAAATTGAATACTTCAGACTC | TT | CCACTCAACTTTAACCCG |
| **11** | GTCCCTGCCTCTATATCT | TT | ATTTATGAGACCAAAGACCAAGTGC | CCAAACAAACAAAACGATAAAAAGG | TT | CCACTCAACTTTAACCCG |
| **12** | GTCCCTGCCTCTATATCT | TT | GCTAATCAACAAAATTATAACCAGC | CCATTACAACCAAAGTTACGAACCA | TT | CCACTCAACTTTAACCCG |
| **13** | GTCCCTGCCTCTATATCT | TT | AGACGAGCGCACAAACCATCAGACC | CATTGAATACGGAAGCAGTTTGAGT | TT | CCACTCAACTTTAACCCG |
| **14** | GTCCCTGCCTCTATATCT | TT | AAATAGTGATTCTCCATCACAGACA | GAGAATATTTGCAGAACAATCACTT | TT | CCACTCAACTTTAACCCG |
| **15** | GTCCCTGCCTCTATATCT | TT | GAAAGAAATCGAAACCAATGGTTGG | GACAATTTCGATTATGAGTCAACTC | TT | CCACTCAACTTTAACCCG |
| **16** | GTCCCTGCCTCTATATCT | TT | AAAGTTGATTTGAGAAACTTGCAAG | TCGCATTGACTTTGGGACTGCAGCT | TT | CCACTCAACTTTAACCCG |
| **17** | GTCCCTGCCTCTATATCT | TT | CTTTGAGACATCGATAAAATGGAAT | GGTCCGACAGATTTAGATTTCTCTT | TT | CCACTCAACTTTAACCCG |
| **18** | GTCCCTGCCTCTATATCT | TT | TGAATATACGGCTCATATTTGGGTC | TTACTTTTGTTAGTTCGTCATCTGC | TT | CCACTCAACTTTAACCCG |
| **19** | GTCCCTGCCTCTATATCT | TT | TCTTCTGGTTTTCCACCTTGAAGTA | TTGATCAGGCGATTGGCCGCCTCAA | TT | CCACTCAACTTTAACCCG |
| **20** | GTCCCTGCCTCTATATCT | TT | GCTTGTAATGATTCTCTTTCCTGAG | TTGATCAACTTTTCCAGAACCATTA | TT | CCACTCAACTTTAACCCG |

**Table S4:** Probe pairs designed for *Al-arrestin-2* HCRs (B1 initiator)

| Pair | Initiator | Spacer | Hybridzation | Hybridzation | Spacer | Initiator |
|---|---|---|---|---|---|---|
| **1** | GAGGAGGGCAGCAAACGG | AA | CATAAGAGATAACAATACCAATTGC | CACCGACGTATAGCCTGACGCGAAT | TA | GAAGAGTCTTCCTTTACG |
| **2** | GAGGAGGGCAGCAAACGG | AA | TAGATCAGTATCTGTTTCTCTGAGC | ATCAGATTGTGAGAATGTGGTTGAA | TA | GAAGAGTCTTCCTTTACG |
| **3** | GAGGAGGGCAGCAAACGG | AA | TTGTTTTGTTGTGCCGATGGGATCA | CCGTTTAATGCAATCCCTCTTTTGT | TA | GAAGAGTCTTCCTTTACG |
| **4** | GAGGAGGGCAGCAAACGG | AA | CATTTACAGGGTATCCATCACGAGA | GATAGACTTTAGAGAGTGATGCGCC | TA | GAAGAGTCTTCCTTTACG |
| **5** | GAGGAGGGCAGCAAACGG | AA | CTTCTGTTCGTTGAATGGCGGAAAC | TACTATAATGACCACTTATGAAATT | TA | GAAGAGTCTTCCTTTACG |
| **6** | GAGGAGGGCAGCAAACGG | AA | GCCGTGAAAGTATAGTTCTCGATCG | TATTATAACGTTAACCGATATGACA | TA | GAAGAGTCTTCCTTTACG |
| **7** | GAGGAGGGCAGCAAACGG | AA | CTGAACATAAAACTCTTACTGATCG | CTCACTTCAAGATTCAATTTGCCGG | TA | GAAGAGTCTTCCTTTACG |
| **8** | GAGGAGGGCAGCAAACGG | AA | CTAATAGCCATTGAAACTGAGTCGG | ATTGATTGCTTAACGAACTGTAGTT | TA | GAAGAGTCTTCCTTTACG |
| **9** | GAGGAGGGCAGCAAACGG | AA | CATATTCTATTCCAAGAGGTGGTCC | CTTTATCAGTTACATACAAGATCAG | TA | GAAGAGTCTTCCTTTACG |
| **10** | GAGGAGGGCAGCAAACGG | AA | AGACGGCGGGGCGTTATGCGGCAAT | ATCTTCAGGACCAGGTTGTATAGTG | TA | GAAGAGTCTTCCTTTACG |
| **11** | GAGGAGGGCAGCAAACGG | AA | AGTTTTTGCACGAGTCGTTCCTGAA | AATGTGAAAGGAATAGCATTGGGAC | TA | GAAGAGTCTTCCTTTACG |
| **12** | GAGGAGGGCAGCAAACGG | AA | ATTTCGTCTTCTTCTCTTCCATATC | AATTGTCGTGAAAAGTTGAGTCCCA | TA | GAAGAGTCTTCCTTTACG |
| **13** | GAGGAGGGCAGCAAACGG | AA | ATCTTTAAGATATTGTTGGTCTACC | GGCTGTGATTTGACCAAATAACTGT | TA | GAAGAGTCTTCCTTTACG |
| **14** | GAGGAGGGCAGCAAACGG | AA | TCTTGATGATCACCAAAATCTCGAT | ACAACTCCATTGAGTGGATCACAAT | TA | GAAGAGTCTTCCTTTACG |

**Table S5:** Probe pairs designed for *Al-atonal* HCRs (B2 initiator)

| Pair | Initiator | Spacer | Hybridzation | Hybridzation | Spacer | Initiator |
|---|---|---|---|---|---|---|
| **1** | CCTCGTAAATCCTCATCA | AA | ATAATTGAAGCCATGAAACAAAGCC | TCAGAGTAGTAATTGAGCGTTTCAT | AA | ATCATCCAGTAAACCGCC |
| **2** | CCTCGTAAATCCTCATCA | AA | ATTTTGATAGCTTTCTATCGTCTCC | AGCTCTGCGCCATTTGGAGTGTTTC | AA | ATCATCCAGTAAACCGCC |
| **3** | CCTCGTAAATCCTCATCA | AA | AAAGGCAATATTTAGAGAATGCATT | CGAAGGCACAACTTCTCGAAGTCGA | AA | ATCATCCAGTAAACCGCC |
| **4** | CCTCGTAAATCCTCATCA | AA | CGTCTTTTCTTAACAATGACTTGAG | CGTCTTTCTCTAGCATTAGCCGCTA | AA | ATCATCCAGTAAACCGCC |
| **5** | CCTCGTAAATCCTCATCA | AA | CGGTGAGCTTTCCTCCACTTTGCTC | TTCATATTTACTTTTAGACTTGTTT | AA | ATCATCCAGTAAACCGCC |
| **6** | CCTCGTAAATCCTCATCA | AA | TAGTCATCGCTGTCCAAAGAGCCAA | CTTTTCCACACCATATCATCCAACT | AA | ATCATCCAGTAAACCGCC |
| **7** | CCTCGTAAATCCTCATCA | AA | CATTAGGTGATTGTAATGATGAGCC | ATGATTTCTGTGATCCGGACATCTT | AA | ATCATCCAGTAAACCGCC |
| **8** | CCTCGTAAATCCTCATCA | AA | ATTCGAGGCGTTTGAGATGACATCG | TAGCTGAAATCTGACTTCTGTCGTA | AA | ATCATCCAGTAAACCGCC |
| **9** | CCTCGTAAATCCTCATCA | AA | GAGGAACAATCGACGTAAGAGTTAG | TCTGGTGTCGACACTGGAGTACAAT | AA | ATCATCCAGTAAACCGCC |
| **10** | CCTCGTAAATCCTCATCA | AA | AATCAAAAAGTGGAAAAGATTGATC | TGGTCTCATTGTCGGAAAGCGAAGA | AA | ATCATCCAGTAAACCGCC |
| **11** | CCTCGTAAATCCTCATCA | AA | TGAAAGAACACCCAACGAAGAGCTT | AAGATGAGTTTGTGTTGGCGAAACT | AA | ATCATCCAGTAAACCGCC |
| **12** | CCTCGTAAATCCTCATCA | AA | GTTATTGTAGCGGATGGCACAACAC | TACTATTGTATACTATAAGGCATTG | AA | ATCATCCAGTAAACCGCC |
| **13** | CCTCGTAAATCCTCATCA | AA | GTGAAGTGGACCAAAGAGACTCAGC | TAAACACGCAATACTTGTTTTAGTT | AA | ATCATCCAGTAAACCGCC |
| **14** | CCTCGTAAATCCTCATCA | AA | AGACAAAGCGCAAAGCCTTCTCATC | TATATGGGACTATAGCTCCACCATA | AA | ATCATCCAGTAAACCGCC |
| **15** | CCTCGTAAATCCTCATCA | AA | CCGAAGGCAACGTCGAGTTTCAAAT | GCGGAGATCCAATCCACGCGCTTTA | AA | ATCATCCAGTAAACCGCC |
| **16** | CCTCGTAAATCCTCATCA | AA | TTTGTGTGAGACTGAGTAACAGGTC | CAACGTATAAATTCAATTCTCTGTC | AA | ATCATCCAGTAAACCGCC |
| **17** | CCTCGTAAATCCTCATCA | AA | TCATAGACTCATTATTATTGGGCGC | TTTCAGTGAAGAGCATTTCAAATGT | AA | ATCATCCAGTAAACCGCC |
| **18** | CCTCGTAAATCCTCATCA | AA | TGTGCTTGCCTGATGAGGTTATACG | TCAGTTTAATTAAATAGTCAAGACT | AA | ATCATCCAGTAAACCGCC |
| **19** | CCTCGTAAATCCTCATCA | AA | AGGTATTCGATGTTTGGGAGAGAAC | TGTTTTGCTGAACAACAACAACAAA | AA | ATCATCCAGTAAACCGCC |
| **20** | CCTCGTAAATCCTCATCA | AA | GTCCTATCACCAACTCTAATAACAT | AACATATCGTCGGGTCATTGCTATA | AA | ATCATCCAGTAAACCGCC |

**Table S6:** Probe pairs designed for *Al-dachshund* HCRs (B2 initiator)

| Pair | Initiator | Spacer | Hybridzation | Hybridzation | Spacer | Initiator |
|---|---|---|---|---|---|---|
| **1** | CCTCGTAAATCCTCATCA | AA | TCCGTTTGTTGTTGTTGCTGTTGCG | TAATTGCAAAACTTTTAATCTACTG | AA | ATCATCCAGTAAACCGCC |
| **2** | CCTCGTAAATCCTCATCA | AA | CTTGTTGGTTTTGAGTGATTTGCTG | GGGGAGTCGGTGACGATTTCAGAGA | AA | ATCATCCAGTAAACCGCC |
| **3** | CCTCGTAAATCCTCATCA | AA | CTGCTGCTGATGTATCTCGTGCTCC | ATTCTGTGTATTGTGTGATATCTGT | AA | ATCATCCAGTAAACCGCC |
| **4** | CCTCGTAAATCCTCATCA | AA | CAAATGCTCTCTCATTTCTCGCTCC | TTTTTGTTCTTCATTCAGTTGTTTT | AA | ATCATCCAGTAAACCGCC |
| **5** | CCTCGTAAATCCTCATCA | AA | AGTTCAGCTTTTTGCAAATGCAATT | TCAAGTTGTGCCGCCAACTCAGCCC | AA | ATCATCCAGTAAACCGCC |
| **6** | CCTCGTAAATCCTCATCA | AA | CAATTGCCAAAAGCCCCTCAATATT | TTTGTTGTTGTCTAGCATTATGAGC | AA | ATCATCCAGTAAACCGCC |
| **7** | CCTCGTAAATCCTCATCA | AA | TTGTTGAGTTTGCTGTTGCTGTGGA | CAGCAATGTTTCCAAACAATGTTCC | AA | ATCATCCAGTAAACCGCC |
| **8** | CCTCGTAAATCCTCATCA | AA | TTGCGGTTTCACTCGCACCCCGTGT | GAGGAGTTGTCGAACTGTTTCGATT | AA | ATCATCCAGTAAACCGCC |
| **9** | CCTCGTAAATCCTCATCA | AA | GCTGACAGGGTTGTCGCTATTGTTA | ACACGATATACCGTTAAATCCAGAA | AA | ATCATCCAGTAAACCGCC |
| **10** | CCTCGTAAATCCTCATCA | AA | TCGTCGTCATCATCAGTAGTGTCAT | CTATTATTACTTTCCACGTCGTCTT | AA | ATCATCCAGTAAACCGCC |
| **11** | CCTCGTAAATCCTCATCA | AA | TGCGCTGCTGGTGTTACTGCTGCTG | ATTATTATTATTGTTTGTATTGTTG | AA | ATCATCCAGTAAACCGCC |
| **12** | CCTCGTAAATCCTCATCA | AA | GTTACGAGAGCTTAAGTTAAGTGCG | GTTATTTCCACTACTATTGACATTA | AA | ATCATCCAGTAAACCGCC |
| **13** | CCTCGTAAATCCTCATCA | AA | TGTTGCTGTGAAAGTAACCATAAAT | GAGTTCGGGACATGTTGTTGTTGCT | AA | ATCATCCAGTAAACCGCC |
| **14** | CCTCGTAAATCCTCATCA | AA | CCAGAACCTTCCGATGTGGCGTTGT | GTTGGATTGAAATCTGAGGCCACGG | AA | ATCATCCAGTAAACCGCC |
| **15** | CCTCGTAAATCCTCATCA | AA | CTGGAGGCGGACTTGCATTTGAAAA | CATTGCTTCCGTGATGCGAAGCTAT | AA | ATCATCCAGTAAACCGCC |
| **16** | CCTCGTAAATCCTCATCA | AA | TGGTGCGGTTCCAGCGCTGAAGTGT | CACAAGTTGTTGCTGTTGCTGAGGA | AA | ATCATCCAGTAAACCGCC |
| **17** | CCTCGTAAATCCTCATCA | AA | TTCGCTGCTAGCATTGCTGCTGCGG | AAGAAAGGAAATGCGGCCGAATAAC | AA | ATCATCCAGTAAACCGCC |
| **18** | CCTCGTAAATCCTCATCA | AA | TATTAAGACTGCTATGCGTTTGCTT | CGGCACTTGTGTTACCACTACTATT | AA | ATCATCCAGTAAACCGCC |
| **19** | CCTCGTAAATCCTCATCA | AA | ATCATCAGAGAAACGGGGCTTTTTA | GTCTATATTTTCACTGGCATAGTCA | AA | ATCATCCAGTAAACCGCC |
| **20** | CCTCGTAAATCCTCATCA | AA | TTGTGTTGCTGTGGCCCTGAGGTGA | GTTGGGTATCCGAGATAGGGCACAA | AA | ATCATCCAGTAAACCGCC |
| **21** | CCTCGTAAATCCTCATCA | AA | TGACTGAACTTCTCTTGGGTGGACG | ATGATGGGACTGGGCTAACAATGCC | AA | ATCATCCAGTAAACCGCC |

**Table S7:** Probe pairs designed for *Al-ey* HCRs (B1 initiator)

| Pair | Initiator | Spacer | Hybridzation | Hybridzation | Spacer | Initiator |
|---|---|---|---|---|---|---|
| **1** | GAGGAGGGCAGCAAACGG | AA | AATACGTGGCCAATAATTGGAAGCG | TACAAATTAATCAATGAATGGTCAC | TA | GAAGAGTCTTCCTTTACG |
| **2** | GAGGAGGGCAGCAAACGG | AA | GGAACTGATACTCCCGGAGATATCA | TCACTGTTCTGACCGGGCACTTGCA | TA | GAAGAGTCTTCCTTTACG |
| **3** | GAGGAGGGCAGCAAACGG | AA | CAGCGAACTGTCCGTTCACTGTTTG | CAGTAGAAGTCGTTGAGTTGTTGAC | TA | GAAGAGTCTTCCTTTACG |
| **4** | GAGGAGGGCAGCAAACGG | AA | TGGACGCGAATAACTTCCGAGTGTT | GTGAACCTGTGCTGCTGACGGACAC | TA | GAAGAGTCTTCCTTTACG |
| **5** | GAGGAGGGCAGCAAACGG | AA | AACATACATGAATAGGAAGAGGCAG | GGATCATAATTACGTCCGGGTGGTG | TA | GAAGAGTCTTCCTTTACG |
| **6** | GAGGAGGGCAGCAAACGG | AA | AAGAGTTAGGGGTCATATTGTTGGC | CCGCTGCAGCCGCTTGTTGTTGAAG | TA | GAAGAGTCTTCCTTTACG |
| **7** | GAGGAGGGCAGCAAACGG | AA | GTAAGTGTCAGCCATTGATGCCATC | CGAATATGACGACATTCCCATTGAA | TA | GAAGAGTCTTCCTTTACG |
| **8** | GAGGAGGGCAGCAAACGG | AA | GCAAAACCCGAATTGATTGGAAGCC | TGTGGAATTGTCGGATATAATGAAT | TA | GAAGAGTCTTCCTTTACG |
| **9** | GAGGAGGGCAGCAAACGG | AA | GCCTGCTCTGCAGCCCTCCTTTGAT | GTGTTATGGCTGTGATGGACGACTG | TA | GAAGAGTCTTCCTTTACG |
| **10** | GAGGAGGGCAGCAAACGG | AA | CGCGTCTATTGGAGAACCAAACCTG | TCAACTTTTCTTCACGTCTCCATTT | TA | GAAGAGTCTTCCTTTACG |
| **11** | GAGGAGGGCAGCAAACGG | AA | CTCAGGCAAATCGATTTTGGCCGCG | AATATGATCGTATGCCTGAATTCTA | TA | GAAGAGTCTTCCTTTACG |
| **12** | GAGGAGGGCAGCAAACGG | AA | TGAGTGCGCTCGAACTCTTTTTCTA | CGCTCTCGAGCGAACACATCGGGAT | TA | GAAGAGTCTTCCTTTACG |
| **13** | GAGGAGGGCAGCAAACGG | AA | TCCTATTTCGCTGTAATTTTCTCTT | CTTCGATCTGTTCGGGTGTAAACGA | TA | GAAGAGTCTTCCTTTACG |
| **14** | GAGGAGGGCAGCAAACGG | AA | AGAGCCGTCGTTGTGAGACGAATCT | TCTAAGTCTCGCTGCGGAATCGCCT | TA | GAAGAGTCTTCCTTTACG |
| **15** | GAGGAGGGCAGCAAACGG | AA | GTGGTCGGCGGCGGAACTGTAGGCA | TTTTTATGAGAATTTCCATTTTCAA | TA | GAAGAGTCTTCCTTTACG |
| **16** | GAGGAGGGCAGCAAACGG | AA | TCTGACCGTTAAGAATACGCAACTT | GATACCAGGGTGACGGTCGCGGCCA | TA | GAAGAGTCTTCCTTTACG |
| **17** | GAGGAGGGCAGCAAACGG | AA | GGTTTGTTGTTCTTTTTGTGCAGCA | ATATACAGCATCGGCTGCCTGTGAG | TA | GAAGAGTCTTCCTTTACG |
| **18** | GAGGAGGGCAGCAAACGG | AA | ACAGACGGTACGTTGTCATTAGTAC | TTACGAAGAACTCGATTGATTGACG | TA | GAAGAGTCTTCCTTTACG |
| **19** | GAGGAGGGCAGCAAACGG | AA | TTTCCCAGGCAAATATTGAGGGACA | CGCCTTCTGACAAAAGTCTGTCCCT | TA | GAAGAGTCTTCCTTTACG |
| **20** | GAGGAGGGCAGCAAACGG | AA | AACATCAGATGTGGCCACACGTGGT | GCGCTTGTACTGCGCTATTTTCGAG | TA | GAAGAGTCTTCCTTTACG |
| **21** | GAGGAGGGCAGCAAACGG | AA | CCGGTCTCGTAATATCGGCCCAATA | CCACCGATAGCTCGCGGACGTATTG | TA | GAAGAGTCTTCCTTTACG |
| **22** | GAGGAGGGCAGCAAACGG | AA | GAATGCGCGATATATCGCAAGGTCG | TAGACACGCAACCATTTGATACCTG | TA | GAAGAGTCTTCCTTTACG |
| **23** | GAGGAGGGCAGCAAACGG | AA | TCACGTAAACACCGCCCAATTGATT | TGGTCGAGTCCGGTAGAGGACGACC | TA | GAAGAGTCTTCCTTTACG |
| **24** | GAGGAGGGCAGCAAACGG | AA | GCAGTGTTCCTGCTGTGATAATACT | ACCCGAGTGACCTTTGTGGGGCATT | TA | GAAGAGTCTTCCTTTACG |
| **25** | GAGGAGGGCAGCAAACGG | AA | CCAGTTGATTTACACCAGTCTTATT | GACACCACTGATGCCGTTGTGATCA | TA | GAAGAGTCTTCCTTTACG |
| **26** | GAGGAGGGCAGCAAACGG | AA | CGATCAATAGATTGAACACGCATTG | ATGGCCAACTATTATCAAACACTAT | TA | GAAGAGTCTTCCTTTACG |
| **27** | GAGGAGGGCAGCAAACGG | AA | AATATAAGGCCATTCAAATGCAATC | AATAAAGATCAGGGCGTATCACAGA | TA | GAAGAGTCTTCCTTTACG |
| **28** | GAGGAGGGCAGCAAACGG | AA | GATTAATATATAGCACTTTGAGCCG | TCACTATAGAGTTCATAACTAAGTC | TA | GAAGAGTCTTCCTTTACG |
| **29** | GAGGAGGGCAGCAAACGG | AA | GAAGTAAAACAGAGCCAAACAGACG | TATATGTTGTATTATAGCAGATGGA | TA | GAAGAGTCTTCCTTTACG |

**Table S8:** Probe pairs designed for *Al-eya* HCRs (B2 initiator)

| Pair | Initiator | Spacer | Hybridzation | Hybridzation | Spacer | Initiator |
|---|---|---|---|---|---|---|
| 1 | CCTCGTAAATCCTCATCA | AA | AGACTTCACAAGTTTGTGGCGTTAG | GTTACACCACATTTTCAAGTTTCAA | AA | ATCATCCAGTAAACCGCC |
| 2 | CCTCGTAAATCCTCATCA | AA | AACATCTTGATCACTTGAAAGCGTT | AACTCAATACTAGTCGCCAAGCAAT | AA | ATCATCCAGTAAACCGCC |
| 3 | CCTCGTAAATCCTCATCA | AA | TTTGAAGTACTAATAAAGTGGCAAG | AAATCGCGTCAGTTCAGACCAAAAA | AA | ATCATCCAGTAAACCGCC |
| 4 | CCTCGTAAATCCTCATCA | AA | AATTGTTCAGCTTATAATCAATTTG | GGTCGCAACACGTCAACTAATACAG | AA | ATCATCCAGTAAACCGCC |
| 5 | CCTCGTAAATCCTCATCA | AA | ACTCCTTCTTCACGAAAATTCTGCG | CAAATGAATGTCAGCAAATTGTTTC | AA | ATCATCCAGTAAACCGCC |
| 6 | CCTCGTAAATCCTCATCA | AA | GAGGCCACACCGAATATCATTACTT | CTGTAAATGTTTTCGATCGCAAAGT | AA | ATCATCCAGTAAACCGCC |
| 7 | CCTCGTAAATCCTCATCA | AA | GCCATTTTTCTGAGCCAGTCTACAC | ATTTCTTTGATGCGTCTATATCTAA | AA | ATCATCCAGTAAACCGCC |
| 8 | CCTCGTAAATCCTCATCA | AA | CTTCCAATTGCATGCCTAAGTCATA | GGTTTTCAGCGAATCTGTTGAGCAG | AA | ATCATCCAGTAAACCGCC |
| 9 | CCTCGTAAATCCTCATCA | AA | TTGGAGTTTCCGTCTTTATAGGGGA | TTTTTGGCGCTATAGGTATAGCATT | AA | ATCATCCAGTAAACCGCC |
| 10 | CCTCGTAAATCCTCATCA | AA | GGCATGGCTGAAACCACCATATGTG | AGTGGAATAGTAGGGCGAATAACCA | AA | ATCATCCAGTAAACCGCC |
| 11 | CCTCGTAAATCCTCATCA | AA | TGGCGATGGATTCAAATAAGCGGCC | AGTATTAGTGGTAAAAGTTGTACTA | AA | ATCATCCAGTAAACCGCC |
| 12 | CCTCGTAAATCCTCATCA | AA | AAGGACTTTGATATGCAGCCTGCAT | CTGCAGCAGCAGCTGAACCGGCGCT | AA | ATCATCCAGTAAACCGCC |
| 13 | CCTCGTAAATCCTCATCA | AA | CCCAACTGTGATGATAATCTGTTGG | CCACTTTTACATCGTTTCCACACAT | AA | ATCATCCAGTAAACCGCC |
| 14 | CCTCGTAAATCCTCATCA | AA | TCTTGTCAGCGAATGGACTCGAAAC | GAGATGAAGATATAGTATTGCCTTT | AA | ATCATCCAGTAAACCGCC |
| 15 | CCTCGTAAATCCTCATCA | AA | CTTCGTCTAAATGTTGCTGTGACTT | TGTCCGAACTTAACAGGTCAGACGA | AA | ATCATCCAGTAAACCGCC |
| 16 | CCTCGTAAATCCTCATCA | AA | GATTAATGTTGAGCAGTAACATACC | CCTTAAAAAGTTTGATCGCAGCAAA | AA | ATCATCCAGTAAACCGCC |
| 17 | CCTCGTAAATCCTCATCA | AA | TGCTCTCATTTATTTATTGCAAGCG | AGCGATGCATATGCATAATCAAACA | AA | ATCATCCAGTAAACCGCC |
| 18 | CCTCGTAAATCCTCATCA | AA | CAACTACTAATTCTCATCAATGGGG | TTTTACATTTTACACCCTCAAACTC | AA | ATCATCCAGTAAACCGCC |
| 19 | CCTCGTAAATCCTCATCA | AA | TCTTATTGCTCTACAAGTCCTTCGA | CTTTCTTAAGCGCTATAACGAGACA | AA | ATCATCCAGTAAACCGCC |
| 20 | CCTCGTAAATCCTCATCA | AA | CTCCAAATCGACTTCCCTTTCGCTT | TGCTCTTATGCTTGATTGAAGGAAG | AA | ATCATCCAGTAAACCGCC |

**Table S9:** Probe pairs designed for *Al-krz* HCRs (B2 initiator)

| Pair | Initiator | Spacer | Hybridzation | Hybridzation | Spacer | Initiator |
|---|---|---|---|---|---|---|
| 1 | CCTCGTAAATCCTCATCA | AA | TTGGGCTTTTCTTGTTGGGCTTTTA | TGGGCTTTTCTTGTTGGGCTTTTCT | AA | ATCATCCAGTAAACCGCC |
| 2 | CCTCGTAAATCCTCATCA | AA | GGTATCCTCACATAGAACAGCTCTC | TTAAATACAGATATACCTAAAGACA | AA | ATCATCCAGTAAACCGCC |
| 3 | CCTCGTAAATCCTCATCA | AA | CAGTCTCTCCTCTTAGTCTTAGTCG | GGATTCAAAATTCAATACTTTATGC | AA | ATCATCCAGTAAACCGCC |
| 4 | CCTCGTAAATCCTCATCA | AA | TCTCTTCGGGTTTAGGATGCATCAG | CTACTTTTGAGGCATTTATGTTATT | AA | ATCATCCAGTAAACCGCC |
| 5 | CCTCGTAAATCCTCATCA | AA | CTTCATGTTTAAGTTGACCGTCAAG | CAATTGTTGAAGACGCTAAGTTTGT | AA | ATCATCCAGTAAACCGCC |
| 6 | CCTCGTAAATCCTCATCA | AA | TGTACATTAACTGCAATATCTTCTC | ACAGTTCGGTTCGAATTATTAGCGA | AA | ATCATCCAGTAAACCGCC |
| 7 | CCTCGTAAATCCTCATCA | AA | CTGAGTTTCTTTTATGAGGTTTTTC | AAACCACTTTCCTGATCGCTAATCG | AA | ATCATCCAGTAAACCGCC |
| 8 | CCTCGTAAATCCTCATCA | AA | CTAATTTCTTTATCAGCCTTTCTTG | CAAAGAAGAACGGAAAAGCATTGGA | AA | ATCATCCAGTAAACCGCC |
| 9 | CCTCGTAAATCCTCATCA | AA | AATCTGGATCTATCAGTACAACGCC | GACCAAATACTTTACGATCTTTTAT | AA | ATCATCCAGTAAACCGCC |
| 10 | CCTCGTAAATCCTCATCA | AA | TATGAAACGTTTCGTTCAAACTGTC | TCTGACATTAAGAGAAGCTATGCTT | AA | ATCATCCAGTAAACCGCC |
| 11 | CCTCGTAAATCCTCATCA | AA | TATGGCTTAATCACCAGTCAAATCG | CTAACTTATCCATCACTTCTTTCTT | AA | ATCATCCAGTAAACCGCC |
| 12 | CCTCGTAAATCCTCATCA | AA | TAAAACTGGAACTCAAGTCGAAGGC | TTTTATGCCACGAGAATGCATTTTA | AA | ATCATCCAGTAAACCGCC |
| 13 | CCTCGTAAATCCTCATCA | AA | TAGTCCAAGACATCATTTACGAGAA | GGCCTTAGAACAGTCGCGACTCATT | AA | ATCATCCAGTAAACCGCC |
| 14 | CCTCGTAAATCCTCATCA | AA | TCCCTCTATCCGTATTACAACTTAC | CCGTACATGACATCGCGTGTGAACA | AA | ATCATCCAGTAAACCGCC |
| 15 | CCTCGTAAATCCTCATCA | AA | AATTTGACTCACGTCCTGAATACTC | CAGCCACAAACACGGTTTTGGTGGT | AA | ATCATCCAGTAAACCGCC |
| 16 | CCTCGTAAATCCTCATCA | AA | AAGAATTGCATTAAAGCAACGATCG | GCTGTTAGTAAAAAGTGAATGATAG | AA | ATCATCCAGTAAACCGCC |
| 17 | CCTCGTAAATCCTCATCA | AA | GGCAGTGCTGCCGCTACTCTTCTGA | CTTGCTTTGTCATTGGTCTCCATTC | AA | ATCATCCAGTAAACCGCC |
| 18 | CCTCGTAAATCCTCATCA | AA | GGCATTTGTTATGTCATATACAACC | CCATTTGGACGTTTGATGGAATGAA | AA | ATCATCCAGTAAACCGCC |
| 19 | CCTCGTAAATCCTCATCA | AA | CTGTTCTGTCTTCTAAATACATAGT | CTGCAGTGTCCCATAATTGTAATCG | AA | ATCATCCAGTAAACCGCC |
| 20 | CCTCGTAAATCCTCATCA | AA | CAACACTTTGTTCGCCCAGAAATAC | TAAAACGAGTAATTAATGACGTCTT | AA | ATCATCCAGTAAACCGCC |

**Table S10:** Probe pairs designed for *Al-otd* HCRs (B3 initiator).

| Pair | Initiator | Spacer | Hybridzation | Hybridzation | Spacer | Initiator |
|---|---|---|---|---|---|---|
| **1** | GTCCCTGCCTCTATATCT | TT | TGATAAGAGTGCATTTTGTCATTAG | CGAATATGATCGGGATGTCATGAAA | TT | CCACTCAACTTTAACCCG |
| **2** | GTCCCTGCCTCTATATCT | TT | GGACTTATTATCCGGATTATATTCC | ATTTCACAACACTTGAAACTTCCAG | TT | CCACTCAACTTTAACCCG |
| **3** | GTCCCTGCCTCTATATCT | TT | TTAATTGGTGGTGTCCTAGCGGTCA | CAGTCGTTAGGCGCAGAGATTCCTC | TT | CCACTCAACTTTAACCCG |
| **4** | GTCCCTGCCTCTATATCT | TT | CATGTGTAGACATTACAGGATTCAT | TCTGATGGCCAGTAGGAGCATGTAA | TT | CCACTCAACTTTAACCCG |
| **5** | GTCCCTGCCTCTATATCT | TT | ATTGAGAGCAGATGAGCCCATAGTG | TGAGCTCATAGTTGGGCCACTCATT | TT | CCACTCAACTTTAACCCG |
| **6** | GTCCCTGCCTCTATATCT | TT | ATTGTCGGCATATGATAATCCATAT | ACAGCACCTAACTGTGTGTGATGAG | TT | CCACTCAACTTTAACCCG |
| **7** | GTCCCTGCCTCTATATCT | TT | AACTTTGAGGTGGATAGCAAGAGGC | CATAATGATATGCTGATGCAGGTCC | TT | CCACTCAACTTTAACCCG |
| **8** | GTCCCTGCCTCTATATCT | TT | GTAGGCAGCAGTTGCCCTCTGCATA | AGCACTGCTTGCCATTTGGTGGTGA | TT | CCACTCAACTTTAACCCG |
| **9** | GTCCCTGCCTCTATATCT | TT | GGAGCGATAGCAGCGGGACTCCATA | CTATTACCAGACATCAGATCCGATA | TT | CCACTCAACTTTAACCCG |
| **10** | GTCCCTGCCTCTATATCT | TT | TAACTGAAGAACATAATGGATTCCC | ATGTGTTGGCACTTGAATTGTTTGA | TT | CCACTCAACTTTAACCCG |
| **11** | GTCCCTGCCTCTATATCT | TT | TGACGGCGGCTTGTATGGAGAATCT | GTTACCGCTCGATGTTATGTTTGGC | TT | CCACTCAACTTTAACCCG |
| **12** | GTCCCTGCCTCTATATCT | TT | TTTGTGGCACTACTAGTCCCGTTTT | GGACTCTTGGCCTTCTTCGGCCTCG | TT | CCACTCAACTTTAACCCG |
| **13** | GTCCCTGCCTCTATATCT | TT | CTCTTCTATTTTTAAACCAAACCTG | GTTGTTGGGCCTGCTGGCGACACTT | TT | CCACTCAACTTTAACCCG |
| **14** | GTCCCTGCCTCTATATCT | TT | TTTTGCTGAACAATGCTTCCAAGAC | CTCTCATAAATATATCAGGATATCT | TT | CCACTCAACTTTAACCCG |
| **15** | GTCCCTGCCTCTATATCT | TT | TTCTCTTCTTTGTTTTCTTGGCGGC | GAGTTGTGCTCTGGTGAAAGTAGTC | TT | CCACTCAACTTTAACCCG |
| **16** | GTCCCTGCCTCTATATCT | TT | AAAAACATATCAGGATATCCAACAC | CTCGGACCGTTCGGCGAAATGGCCC | TT | CCACTCAACTTTAACCCG |
| **17** | GTCCCTGCCTCTATATCT | TT | TGTGATGCCTGCGATCCCATTCATT | ATGCAAGAGATCGACGGCACCGGGT | TT | CCACTCAACTTTAACCCG |
| **18** | GTCCCTGCCTCTATATCT | TT | GGACGTTGGCTGAAAAGCCAGTGTT | GCCAGAAGCGAATGGATTGCAAGAC | TT | CCACTCAACTTTAACCCG |
| **19** | GTCCCTGCCTCTATATCT | TT | GATGATGGCGGAGACGAAGGCGATG | ACTCCTGATGACGATATGTTTCTGA | TT | CCACTCAACTTTAACCCG |
| **20** | GTCCCTGCCTCTATATCT | TT | AAGTACCTCCCGTTGGAGTCGAATT | TATTGCACCCACTCAGAGATGTTGT | TT | CCACTCAACTTTAACCCG |
| **21** | GTCCCTGCCTCTATATCT | TT | TTCCTCTTCGGATACTGAGGAGTAC | CCCAGGACTACCGACTTGATGCAGC | TT | CCACTCAACTTTAACCCG |
| **22** | GTCCCTGCCTCTATATCT | TT | TTGTTGGATGTGTTGTGTTTTGATG | TGATGTGCGGATAATAGTTGTGATG | TT | CCACTCAACTTTAACCCG |
| **23** | GTCCCTGCCTCTATATCT | TT | CGTTGCGGAGATATTGCAGGAAAAA | GGAATCCGTGTAATGACCGAATCGT | TT | CCACTCAACTTTAACCCG |
| **24** | GTCCCTGCCTCTATATCT | TT | GTTAACTTCTAGCATGTTTTTGCCT | CGACAACGTTGAGCTGTATAACTGT | TT | CCACTCAACTTTAACCCG |
| **25** | GTCCCTGCCTCTATATCT | TT | CTATCTCTGTCTCTAAAACTTATGC | AGAGATGTGCACTTATAGAGATGGT | TT | CCACTCAACTTTAACCCG |
| **26** | GTCCCTGCCTCTATATCT | TT | GACAATTAAGCTTTGATTATTCGGC | TCAACTCACAGACAAAACACTCGTG | TT | CCACTCAACTTTAACCCG |
| **27** | GTCCCTGCCTCTATATCT | TT | TTGAGCTTTGGTTTTGGTCACATTG | TCAAAGTTTAGTTCAACAGTTGAGA | TT | CCACTCAACTTTAACCCG |
| **28** | GTCCCTGCCTCTATATCT | TT | TCTGACGAATCCATCATTAATTGAC | ACTTACCCGTCAGTATTTGAATCAG | TT | CCACTCAACTTTAACCCG |
| **29** | GTCCCTGCCTCTATATCT | TT | CTATTTCGGAGATAACATTCGCAGG | CGATTGAATGACAATCAAAATTATT | TT | CCACTCAACTTTAACCCG |
| **30** | GTCCCTGCCTCTATATCT | TT | TGTTCACATCAGTTAACTAGAGATC | ACTTTGACTGTGCTTTCAAACAGTC | TT | CCACTCAACTTTAACCCG |

**Table S11:** Probe pairs designed for *Al-peropsin* (B3 initiator).

| Pair | Initiator | Spacer | Hybridzation | Hybridzation | Spacer | Initiator |
|---|---|---|---|---|---|---|
| **1** | GTCCCTGCCTCTATATCT | TT | TCTCTAAACTGTTGGCTTATCCGAC | ATTTCACTTTCGATTTTCTGCTAAT | TT | CCACTCAACTTTAACCCG |
| **2** | GTCCCTGCCTCTATATCT | TT | AACATCCACAAAGCGAACAACTCTT | TTGAGTTAAGCTACTCTGGTCTTTG | TT | CCACTCAACTTTAACCCG |
| **3** | GTCCCTGCCTCTATATCT | TT | AAGTCGATTATTGGTTAGGTAGTAG | AAAAACGGATGATAAGAAAGCTCCT | TT | CCACTCAACTTTAACCCG |
| **4** | GTCCCTGCCTCTATATCT | TT | GCAAATAGTGGCGGAACCAAAGTGA | ATTGGATTAAACAGTGTAGAGGTTT | TT | CCACTCAACTTTAACCCG |
| **5** | GTCCCTGCCTCTATATCT | TT | ATACTGTCCACAAGCAGAGTATCGC | AAAGAGGAACTGTTTTAGGATCACC | TT | CCACTCAACTTTAACCCG |
| **6** | GTCCCTGCCTCTATATCT | TT | AACTATGATAGTCGACATTATAGTT | GGGAGACCACGAAAACACAAATACA | TT | CCACTCAACTTTAACCCG |
| **7** | GTCCCTGCCTCTATATCT | TT | ATTAATTCCAGCGTTTCCTCGCTTA | TTTTTCCCTAACCCAAATGTCTTCT | TT | CCACTCAACTTTAACCCG |
| **8** | GTCCCTGCCTCTATATCT | TT | GCATAATAGCAGTAGAAAATAACGC | GAACCATTCCTGACTTTACTAACAA | TT | CCACTCAACTTTAACCCG |
| **9** | GTCCCTGCCTCTATATCT | TT | TTATGTCTCCAATCTATAGTACACG | ATTATGAATGATTTATAAGCCGCGT | TT | CCACTCAACTTTAACCCG |
| **10** | GTCCCTGCCTCTATATCT | TT | CCCAACCTATTAATGGCATTAATGC | TTATACTTGAATCCAAGCCATATCT | TT | CCACTCAACTTTAACCCG |
| **11** | GTCCCTGCCTCTATATCT | TT | ATTTGCAGTAATTGTATTTATCCCG | AAGAGTACAACCAAACAGCGTTTAT | TT | CCACTCAACTTTAACCCG |
| **12** | GTCCCTGCCTCTATATCT | TT | CAAAGCCAGTAATGTTAAAGTGCCG | CTTTTGGCAAGAAATATTATATCTG | TT | CCACTCAACTTTAACCCG |
| **13** | GTCCCTGCCTCTATATCT | TT | AAAGCATATGCTTGACAGCCATCAT | TGTGCAGAACCTGCCAGAAACCCCA | TT | CCACTCAACTTTAACCCG |

**Table S12:** Probe pairs designed for *Al-rhodopsin* HCRs (B3 initiator)

| Pair | Initiator | Spacer | Hybridzation | Hybridzation | Spacer | Initiator |
|---|---|---|---|---|---|---|
| **1** | GTCCCTGCCTCTATATCT | TT | GTTTTATTAAAATGGGTAAACTTTA | CCTACACGACGCTCTTCCGATCTCA | TT | CCACTCAACTTTAACCCG |
| **2** | GTCCCTGCCTCTATATCT | TT | CTGATTGTTGTTAACCTCAGATTCG | TTTAACTCATAAAAGTGTGTGTTGT | TT | CCACTCAACTTTAACCCG |
| **3** | GTCCCTGCCTCTATATCT | TT | AGCCAACATATGACACGAATAGAAC | AAACGGATTATAGCAGATTGAACTC | TT | CCACTCAACTTTAACCCG |
| **4** | GTCCCTGCCTCTATATCT | TT | AATATGAAAGCAATTCAAAGGCAAC | GATGACTTCTGGACTAAAATCGGCA | TT | CCACTCAACTTTAACCCG |
| **5** | GTCCCTGCCTCTATATCT | TT | CTTTTCTGGATATCATTCTCTCCCG | ATAATCTTTCGATCAGATCTTTTGA | TT | CCACTCAACTTTAACCCG |
| **6** | GTCCCTGCCTCTATATCT | TT | TCTCACATAAGCGACTGTTGTTATC | ATATTTTCTTCTTATCTGACACCCG | TT | CCACTCAACTTTAACCCG |
| **7** | GTCCCTGCCTCTATATCT | TT | TCTGGATATATAGCTCTGCATCGAG | ATTATTTGTCGAGATCTATTTTGTG | TT | CCACTCAACTTTAACCCG |
| **8** | GTCCCTGCCTCTATATCT | TT | GTATCTATCGATAGCTATTACGGCC | TTGATAAAACCTAGAGTTTATTGCT | TT | CCACTCAACTTTAACCCG |
| **9** | GTCCCTGCCTCTATATCT | TT | ACCTGCATTGAAGGCAATAGTTTGC | GTGAACGTTGAAACATATACAAAAG | TT | CCACTCAACTTTAACCCG |
| **10** | GTCCCTGCCTCTATATCT | TT | GCAATATCCTCGCGATAATGAACGG | AAATCACACCAAAAGGCCAGTTATT | TT | CCACTCAACTTTAACCCG |
| **11** | GTCCCTGCCTCTATATCT | TT | TGGTTTGATCGGAAAAATTGAAACC | ACTTGAGTTTTGCACTGAATCCATC | TT | CCACTCAACTTTAACCCG |

**Table S13:** Probe pairs designed for *Al-Six3* HCRs (B2 initiator)

| Pair | Initiator | Spacer | Hybridzation | Hybridzation | Spacer | Initiator |
|---|---|---|---|---|---|---|
| **1** | CCTCGTAAATCCTCATCA | AA | CTGGAAAACAGACTTTGTATTGAAG | AAAACACAGGTCATGCTAAACATCG | AA | ATCATCCAGTAAACCGCC |
| **2** | CCTCGTAAATCCTCATCA | AA | TATGTGCGTGCCACCTCACAATTTC | AATCAGTTATTAATTGTTTGCTTGA | AA | ATCATCCAGTAAACCGCC |
| **3** | CCTCGTAAATCCTCATCA | AA | CTTCAAGCACCGAGGCTTCATCATT | CTTTTGGTCACACGACACGCCCCTT | AA | ATCATCCAGTAAACCGCC |
| **4** | CCTCGTAAATCCTCATCA | AA | CCAACATCTACATCATCGTCATCAT | GTCGAGACGGCTGTCGGGCTTAGGG | AA | ATCATCCAGTAAACCGCC |
| **5** | CCTCGTAAATCCTCATCA | AA | AGCACTGTTGGCGGTACCGGCTAAT | ATTGAGAAGATGCGTTTTCTGGTTA | AA | ATCATCCAGTAAACCGCC |
| **6** | CCTCGTAAATCCTCATCA | AA | AATGTCCGCCACTTGAACTACGGTT | TCGAACCATTCCTGCCACTACTGCC | AA | ATCATCCAGTAAACCGCC |
| **7** | CCTCGTAAATCCTCATCA | AA | TTTCCCACTTGTGTTGGTGTTAATC | TCACGTTGTCTTCTGTTTTTAAACC | AA | ATCATCCAGTAAACCGCC |
| **8** | CCTCGTAAATCCTCATCA | AA | TTCTTTAAAACAGTGCGTCTTCTGT | CCACTCTCTTAGTAGAGATCTGGTT | AA | ATCATCCAGTAAACCGCC |
| **9** | CCTCGTAAATCCTCATCA | AA | CATCGCTTGTAATTTCGAATGTGAT | AGCTTCTTGATAGTGTGCCTCCAAC | AA | ATCATCCAGTAAACCGCC |
| **10** | CCTCGTAAATCCTCATCA | AA | GTATGAAAAGCCACTAAAGCTCTGG | ATATGATATAGTTCTCTGAAATTTC | AA | ATCATCCAGTAAACCGCC |
| **11** | CCTCGTAAATCCTCATCA | AA | CAGCACAGTTTGGATGAGCCACTGG | TTAGAACAGATTCATTTTTATTAAG | AA | ATCATCCAGTAAACCGCC |
| **12** | CCTCGTAAATCCTCATCA | AA | AATATCTCCACTTTCTTCTAGTGTC | TGACCACAGAAACCTTCCTAGTCTT | AA | ATCATCCAGTAAACCGCC |
| **13** | CCTCGTAAATCCTCATCA | AA | AAGTTTAAAGTCGGTAACATAAACA | CAAACAGCGGCCACTTGACTCACAG | AA | ATCATCCAGTAAACCGCC |
| **14** | CCTCGTAAATCCTCATCA | AA | TGGACTGGCTGAAGAACCAGTTGAT | AGCTGCCGATGCAGTGGCAGCTGAT | AA | ATCATCCAGTAAACCGCC |
| **15** | CCTCGTAAATCCTCATCA | AA | CCAGATGTAGGGATTGGCTGATGCA | GCTGTGCCCAAACCCAATGCCGTCA | AA | ATCATCCAGTAAACCGCC |
| **16** | CCTCGTAAATCCTCATCA | AA | ACGATCGAGACGCTATTGTGTTATT | AAGGCAAAGGCAGCGCCGACAATGT | AA | ATCATCCAGTAAACCGCC |
| **17** | CCTCGTAAATCCTCATCA | AA | GATGAACAGCTAGATGATGATGAGG | AATGATAAATATGCTGCAGAATTGG | AA | ATCATCCAGTAAACCGCC |
| **18** | CCTCGTAAATCCTCATCA | AA | AAGCACCACAACTACTAATATCCGT | AAGATGAGCCGTCACCAGGATGAAG | AA | ATCATCCAGTAAACCGCC |
| **19** | CCTCGTAAATCCTCATCA | AA | AGAAGCTGTCTCACCAGAAATGCCG | ATATATATTACTAATAACCAGATAA | AA | ATCATCCAGTAAACCGCC |
| **20** | CCTCGTAAATCCTCATCA | AA | TGTAAACTATCACTAGAGGATTGCG | ATGCAATAGTGATATCAAGAACACA | AA | ATCATCCAGTAAACCGCC |

**Table S14:** Probe pairs designed for *Al-sine oculis* (B3 initiator)

| Pair | Initiator | Spacer | Hybridzation | Hybridzation | Spacer | Initiator |
|---|---|---|---|---|---|---|
| **1** | GTCCCTGCCTCTATATCT | TT | CATTCATACTCAGCAATAGGAATAG | CATTCATACTAACTTTACGCTTAAG | TT | CCACTCAACTTTAACCCG |
| **2** | GTCCCTGCCTCTATATCT | TT | TCGTGTCTCTCTCTCTCTCTCTCTC | AAAATTCACTACCAAACATCAAAAT | TT | CCACTCAACTTTAACCCG |
| **3** | GTCCCTGCCTCTATATCT | TT | TTTCTCTCTCATAGCTCTCTATCCT | GCTCTCTGTCTCTCTCTCACACCTC | TT | CCACTCAACTTTAACCCG |
| **4** | GTCCCTGCCTCTATATCT | TT | AATTGAGTCTCTGAGATAATGGTGG | ATTAAGTGACCAAAACTTGACAGAC | TT | CCACTCAACTTTAACCCG |
| **5** | GTCCCTGCCTCTATATCT | TT | TTTATCGTACAGAAAGTCTGGTTTG | ATAAGTAATAACTGGCAGTTTGTCT | TT | CCACTCAACTTTAACCCG |
| **6** | GTCCCTGCCTCTATATCT | TT | CACAGTTAATGGTCTCTCACAGTTG | TCCTAATAATTGCTGGAATGTGTAA | TT | CCACTCAACTTTAACCCG |
| **7** | GTCCCTGCCTCTATATCT | TT | ATCAACACCAGAAGCAGCCGCCACT | ATGTGATCCTCCGATAGTAGATCCG | TT | CCACTCAACTTTAACCCG |
| **8** | GTCCCTGCCTCTATATCT | TT | GCCAAATGGTGGGAATGATGAGGAT | ATGACTAGCGCAGACGGCGCCACAT | TT | CCACTCAACTTTAACCCG |
| **9** | GTCCCTGCCTCTATATCT | TT | ATCGAAATCCGTGATACATCGCCGA | ATGTCGATCCGATGTGGTGATGATT | TT | CCACTCAACTTTAACCCG |
| **10** | GTCCCTGCCTCTATATCT | TT | TGTCACTGAACTCAGCTGTTGATGA | CGCAGTAGCCGCTGATTGACTGGCC | TT | CCACTCAACTTTAACCCG |
| **11** | GTCCCTGCCTCTATATCT | TT | TTACCCATGTTGTGAACTAAATGGT | TGACCTGTATTCGACTGATTGTGAT | TT | CCACTCAACTTTAACCCG |
| **12** | GTCCCTGCCTCTATATCT | TT | CCTCATTGGAAGAGTCATCGCCGCT | AACTACTACTCGTTCGAAGTGATGT | TT | CCACTCAACTTTAACCCG |
| **13** | GTCCCTGCCTCTATATCT | TT | TGGTTTAGCACCAGATTTAATACTA | GTTGCCAGAGTTGGAGGAATCATTG | TT | CCACTCAACTTTAACCCG |
| **14** | GTCCCTGCCTCTATATCT | TT | CTATCTTTAGCTTCAGCGGCTCTAT | CCAGAATTGGTTTTATCGGAAGGAT | TT | CCACTCAACTTTAACCCG |
| **15** | GTCCCTGCCTCTATATCT | TT | TCACTTGTGTTGTCGTAAGTCCTGT | TCTGTCGCCTATTTTTGAACCAATT | TT | CCACTCAACTTTAACCCG |
| **16** | GTCCCTGCCTCTATATCT | TT | CGAAGGATAAGGATTGTGAGCATAC | TTCGGCCAGTTCTCTCTTTTCTCGA | TT | CCACTCAACTTTAACCCG |
| **17** | GTCCCTGCCTCTATATCT | TT | TTGAAACAATAAGATGTCTCTTCTC | TCGCGAAGAACATTTCGCGATTTTT | TT | CCACTCAACTTTAACCCG |
| **18** | GTCCCTGCCTCTATATCT | TT | TGCGTCTTATGCGGTATTTGCCCAC | CCCATATAGTACGAGGTAATGGAAA | TT | CCACTCAACTTTAACCCG |
| **19** | GTCCCTGCCTCTATATCT | TT | TGCTTCTATATAATGTGCTTTCAGC | ACCTAATGGACGTCCGCGAAGACGC | TT | CCACTCAACTTTAACCCG |
| **20** | GTCCCTGCCTCTATATCT | TT | AAAATTGTGACTTTCTAGGATTCGG | TTGTAGTTTTGGATGAGATGATGGA | TT | CCACTCAACTTTAACCCG |
| **21** | GTCCCTGCCTCTATATCT | TT | CTACCAAAGCTTTAGCTTTAAGTAC | GTTCTTTGAAATTGCCCCTGTTAAA | TT | CCACTCAACTTTAACCCG |
| **22** | GTCCCTGCCTCTATATCT | TT | AGGCAATGACCATAAAAATCGTCCC | TTCATTCTTTTGTAACTGTTCACAT | TT | CCACTCAACTTTAACCCG |
| **23** | GTCCCTGCCTCTATATCT | TT | ACTTCACAAACACATGCAACTTGTT | CTTTCAATATTGCCCGACTGTTGAA | TT | CCACTCAACTTTAACCCG |
| **24** | GTCCCTGCCTCTATATCT | TT | CTGGCGAAGTAGGGTTGGGACATCC | GAGTGAATCCAAACGATGGTAGAAG | TT | CCACTCAACTTTAACCCG |
| **25** | GTCCCTGCCTCTATATCT | TT | GTACAGCACTGGCCATTCCTGGTAT | GAGGACCACCAATATGCCCACTGAG | TT | CCACTCAACTTTAACCCG |
| **26** | GTCCCTGCCTCTATATCT | TT | TCCGACAATCATTCCACTTATTAAG | AATAGCGTTGGACGCATCCGTCATA | TT | CCACTCAACTTTAACCCG |
| **27** | GTCCCTGCCTCTATATCT | TT | TCAGATTTGAGGTAAGGCTCTGAAG | CAATTCATTAAAACCCGAACAAATT | TT | CCACTCAACTTTAACCCG |
| **28** | GTCCCTGCCTCTATATCT | TT | TGATTAGACTTTGCGCACGTTTTGC | AGTATTGTCTGTCGAGCAAAACAGC | TT | CCACTCAACTTTAACCCG |
| **29** | GTCCCTGCCTCTATATCT | TT | TTTACTCAGTTTGAGTTCTGTTCTG | TCTGCCGACAGATTCTCACTGAATG | TT | CCACTCAACTTTAACCCG |
| **30** | GTCCCTGCCTCTATATCT | TT | AATTCAATCTAAAACCCTTGAGGCC | AGTATTCACAGACACACAAATATCT | TT | CCACTCAACTTTAACCCG |
| **31** | GTCCCTGCCTCTATATCT | TT | GCCGGGAGAACAAAATCAACTGCTT | AGAAGCGTTGCCTGTATTAGATGTT | TT | CCACTCAACTTTAACCCG |
| **32** | GTCCCTGCCTCTATATCT | TT | AAGAAGTCTCTCTGAAGAAGCTCTC | TGCTCAAGACTGAAGAAGTCCTCTC | TT | CCACTCAACTTTAACCCG |
| **33** | GTCCCTGCCTCTATATCT | TT | TGACTGAAGTGGCAACTCGAGACTC | GAAGAAATTCTCTGTGAAGAAAAAT | TT | CCACTCAACTTTAACCCG |
| **34** | GTCCCTGCCTCTATATCT | TT | TAGCCAAAAGTCAAGCTTAATCTGT | GCATGAGATTTGGAAGAACCGAAAC | TT | CCACTCAACTTTAACCCG |
| **35** | GTCCCTGCCTCTATATCT | TT | ACTGCGAAACTGCAATACTTTCCGC | AGCGCAAGGCATGCATTACATTATG | TT | CCACTCAACTTTAACCCG |
| **36** | GTCCCTGCCTCTATATCT | TT | CAATGGCTGCTCAATACCAACTCTC | TCACAGACTAGTCGTTCCCAAACGA | TT | CCACTCAACTTTAACCCG |
| **37** | GTCCCTGCCTCTATATCT | TT | AACCTTGCAATCACCGCCCGTTTTA | TTGCTATTGGCTCAAATTTTTCTGC | TT | CCACTCAACTTTAACCCG |
| **38** | GTCCCTGCCTCTATATCT | TT | TCTTTCATTTCTTTCCATCCCATCT | TCTTCATCATACTGTAGTGGAAGTA | TT | CCACTCAACTTTAACCCG |

**Table S15:** Probe pairs designed for *Al-toy* HCRs (B1 initiator)

| Pair | Initiator | Spacer | Hybridzation | Hybridzation | Spacer | Initiator |
|---|---|---|---|---|---|---|
| 1 | GAGGAGGGCAGCAAACGG | AA | CGATTGTGGATTTGGTTGAGCAGAC | TATAAAATGATTGGAGTAATTGTGA | TA | GAAGAGTCTTCCTTTACG |
| 2 | GAGGAGGGCAGCAAACGG | AA | AAATAAGGTTGAGATGATTGAGGCG | TCGAACGGATCCACAGAATGGCCCA | TA | GAAGAGTCTTCCTTTACG |
| 3 | GAGGAGGGCAGCAAACGG | AA | TGTTGAAGTTTTGGATTGAATTACT | GCGGAGGAGGCGTTAGGGCCGAGAA | TA | GAAGAGTCTTCCTTTACG |
| 4 | GAGGAGGGCAGCAAACGG | AA | GTTGACGAAATTTCGGTTGTTTTCC | CGATGAGAGTGTGGCCAAAGAGGAA | TA | GAAGAGTCTTCCTTTACG |
| 5 | GAGGAGGGCAGCAAACGG | AA | ATTTGGTTATCGGCAGGAGTGTTAC | TTTGAACTCTTGTTAGTATTGCTAT | TA | GAAGAGTCTTCCTTTACG |
| 6 | GAGGAGGGCAGCAAACGG | AA | TGGCTCTTCTGTTTGAAAACCAAAC | TTCTCAATTTTTCTTCTCGTCTCCA | TA | GAAGAGTCTTCCTTTACG |
| 7 | GAGGAGGGCAGCAAACGG | AA | ATCAGCCAGTTTCTCGCGAGCAAAC | AATTCTAGCTTCCGGTAAACTTATT | TA | GAAGAGTCTTCCTTTACG |
| 8 | GAGGAGGGCAGCAAACGG | AA | TTCGCTGTAATCTTCTTTTGAGTCG | TTTGTTCATCAGTAAAAGCAGTCCT | TA | GAAGAGTCTTCCTTTACG |
| 9 | GAGGAGGGCAGCAAACGG | AA | CCATCTGATGAATAATTATTTTCGG | CTCAATTGCGACTCTTCATCTGTAG | TA | GAAGAGTCTTCCTTTACG |
| 10 | GAGGAGGGCAGCAAACGG | AA | GCCGAAGACAGCCTTCATGAGCACT | TGACATCACAAGTGAGCTTATCTGT | TA | GAAGAGTCTTCCTTTACG |
| 11 | GAGGAGGGCAGCAAACGG | AA | TACCGGAGGGCTACCACTGGATGTT | TGTCACATCTCCACAATTAGTTAGA | TA | GAAGAGTCTTCCTTTACG |
| 12 | GAGGAGGGCAGCAAACGG | AA | GGTGAATGCGGTGATGAGGAAGGGG | CTACCATTATTTTGGCCAAAGTGCT | TA | GAAGAGTCTTCCTTTACG |
| 13 | GAGGAGGGCAGCAAACGG | AA | GATGTGCTGAAATCATGCGAAATTT | ATGATGTGGAATGATACCATTGCCA | TA | GAAGAGTCTTCCTTTACG |
| 14 | GAGGAGGGCAGCAAACGG | AA | TTTCTGTGCGGCCAAATTTCTTAAT | CAAATTATTGCCACTGACTTGTTGT | TA | GAAGAGTCTTCCTTTACG |
| 15 | GAGGAGGGCAGCAAACGG | AA | GTATCGGTGTTGCAAACGTTTTCAG | CTGTTTATTGATGAGACACTTGGAA | TA | GAAGAGTCTTCCTTTACG |
| 16 | GAGGAGGGCAGCAAACGG | AA | AAATGGACGGACACTCGCGTTTGTA | ACAATCTATCACGTATTTCCCACGC | TA | GAAGAGTCTTCCTTTACG |
| 17 | GAGGAGGGCAGCAAACGG | AA | GGCAACACGTGGTTTACTTCCGCCT | AGTAATCGCGTGCACTACTATCGAA | TA | GAAGAGTCTTCCTTTACG |
| 18 | GAGGAGGGCAGCAAACGG | AA | TAATAGCGGCCAAGAATCTTCGAGA | GCTCTTGGTTTGATAGACCCAGTTT | TA | GAAGAGTCTTCCTTTACG |
| 19 | GAGGAGGGCAGCAAACGG | AA | ATATATCACAAGGTCGGGCACCGCT | AGCCGTTCGATATTTGTAGCATTCG | TA | GAAGAGTCTTCCTTTACG |
| 20 | GAGGAGGGCAGCAAACGG | AA | TGAGTCCGGCAGTGGCCGACCGTTC | TGCCAAATCTATGATTTTCTGTCGG | TA | GAAGAGTCTTCCTTTACG |
| 21 | GAGGAGGGCAGCAAACGG | AA | CTTTGACCCTTATGATGCATGTCTG | TAGACTCCACCCAATTGATTGATTC | TA | GAAGAGTCTTCCTTTACG |

**Table S16:** Probe pairs designed for *Al-toy* HCRs (B2 initiator).

| Pair | Initiator | Spacer | Hybridzation | Hybridzation | Spacer | Initiator |
|---|---|---|---|---|---|---|
| 1 | CCTCGTAAATCCTCATCA | AA | CGATTGTGGATTTGGTTGAGCAGAC | TATAAAATGATTGGAGTAATTGTGA | AA | ATCATCCAGTAAACCGCC |
| 2 | CCTCGTAAATCCTCATCA | AA | AAATAAGGTTGAGATGATTGAGGCG | TCGAACGGATCCACAGAATGGCCCA | AA | ATCATCCAGTAAACCGCC |
| 3 | CCTCGTAAATCCTCATCA | AA | TGTTGAAGTTTTGGATTGAATTACT | GCGGAGGAGGCGTTAGGGCCGAGAA | AA | ATCATCCAGTAAACCGCC |
| 4 | CCTCGTAAATCCTCATCA | AA | GTTGACGAAATTTCGGTTGTTTTCC | CGATGAGAGTGTGGCCAAAGAGGAA | AA | ATCATCCAGTAAACCGCC |
| 5 | CCTCGTAAATCCTCATCA | AA | ATTTGGTTATCGGCAGGAGTGTTAC | TTTGAACTCTTGTTAGTATTGCTAT | AA | ATCATCCAGTAAACCGCC |
| 6 | CCTCGTAAATCCTCATCA | AA | TGGCTCTTCTGTTTGAAAACCAAAC | TTCTCAATTTTTCTTCTCGTCTCCA | AA | ATCATCCAGTAAACCGCC |
| 7 | CCTCGTAAATCCTCATCA | AA | ATCAGCCAGTTTCTCGCGAGCAAAC | AATTCTAGCTTCCGGTAAACTTATT | AA | ATCATCCAGTAAACCGCC |
| 8 | CCTCGTAAATCCTCATCA | AA | TTCGCTGTAATCTTCTTTTGAGTCG | TTTGTTCATCAGTAAAAGCAGTCCT | AA | ATCATCCAGTAAACCGCC |
| 9 | CCTCGTAAATCCTCATCA | AA | CCATCTGATGAATAATTATTTTCGG | CTCAATTGCGACTCTTCATCTGTAG | AA | ATCATCCAGTAAACCGCC |
| 10 | CCTCGTAAATCCTCATCA | AA | GCCGAAGACAGCCTTCATGAGCACT | TGACATCACAAGTGAGCTTATCTGT | AA | ATCATCCAGTAAACCGCC |
| 11 | CCTCGTAAATCCTCATCA | AA | TACCGGAGGGCTACCACTGGATGTT | TGTCACATCTCCACAATTAGTTAGA | AA | ATCATCCAGTAAACCGCC |
| 12 | CCTCGTAAATCCTCATCA | AA | GGTGAATGCGGTGATGAGGAAGGGG | CTACCATTATTTTGGCCAAAGTGCT | AA | ATCATCCAGTAAACCGCC |
| 13 | CCTCGTAAATCCTCATCA | AA | GATGTGCTGAAATCATGCGAAATTT | ATGATGTGGAATGATACCATTGCCA | AA | ATCATCCAGTAAACCGCC |
| 14 | CCTCGTAAATCCTCATCA | AA | TTTCTGTGCGGCCAAATTTCTTAAT | CAAATTATTGCCACTGACTTGTTGT | AA | ATCATCCAGTAAACCGCC |
| 15 | CCTCGTAAATCCTCATCA | AA | GTATCGGTGTTGCAAACGTTTTCAG | CTGTTTATTGATGAGACACTTGGAA | AA | ATCATCCAGTAAACCGCC |
| 16 | CCTCGTAAATCCTCATCA | AA | AAATGGACGGACACTCGCGTTTGTA | ACAATCTATCACGTATTTCCCACGC | AA | ATCATCCAGTAAACCGCC |
| 17 | CCTCGTAAATCCTCATCA | AA | GGCAACACGTGGTTTACTTCCGCCT | AGTAATCGCGTGCACTACTATCGAA | AA | ATCATCCAGTAAACCGCC |
| 18 | CCTCGTAAATCCTCATCA | AA | TAATAGCGGCCAAGAATCTTCGAGA | GCTCTTGGTTTGATAGACCCAGTTT | AA | ATCATCCAGTAAACCGCC |
| 19 | CCTCGTAAATCCTCATCA | AA | ATATATCACAAGGTCGGGCACCGCT | AGCCGTTCGATATTTGTAGCATTCG | AA | ATCATCCAGTAAACCGCC |
| 20 | CCTCGTAAATCCTCATCA | AA | TGAGTCCGGCAGTGGCCGACCGTTC | TGCCAAATCTATGATTTTCTGTCGG | AA | ATCATCCAGTAAACCGCC |
| 21 | CCTCGTAAATCCTCATCA | AA | CTTTGACCCTTATGATGCATGTCTG | TAGACTCCACCCAATTGATTGATTC | AA | ATCATCCAGTAAACCGCC |

**Table S17:** Probe pairs designed for *Al-wg* HCRs (B2 initiator).

| Pair | Initiator | Spacer | Hybridzation | Hybridzation | Spacer | Initiator |
|---|---|---|---|---|---|---|
| **1** | CCTCGTAAATCCTCATCA | AA | GATCGGTTTTAACCATTTCAACAGG | AGTGTTTACTCGTTTTCAACACCAT | AA | ATCATCCAGTAAACCGCC |
| **2** | CCTCGTAAATCCTCATCA | AA | GCTTTACAAACCTTGCATTTGACCT | CACAAACACGTATGAACTGTTTTTC | AA | ATCATCCAGTAAACCGCC |
| **3** | CCTCGTAAATCCTCATCA | AA | GCTCTAATTCCTCTCTGATTTCAGT | AACACCAATGAAAAGTACAATTACA | AA | ATCATCCAGTAAACCGCC |
| **4** | CCTCGTAAATCCTCATCA | AA | ACATCCTTCAACACCGATAGAAGTG | ATATCCTCGCCCACAACATAATAAA | AA | ATCATCCAGTAAACCGCC |
| **5** | CCTCGTAAATCCTCATCA | AA | ACTCCAAATTTAGGATTAGGAGTAC | TTACAAACTCTTCCTTTAGTCCCAA | AA | ATCATCCAGTAAACCGCC |
| **6** | CCTCGTAAATCCTCATCA | AA | TGCTGACCATTACCCTCGAAGCGCC | TTTTTCTTGTTAATCCTCTGTAGTC | AA | ATCATCCAGTAAACCGCC |
| **7** | CCTCGTAAATCCTCATCA | AA | GTCTCTGAATGGAGGCAACCGCATC | AAATCTATCTTTAAGATTATTGCCA | AA | ATCATCCAGTAAACCGCC |
| **8** | CCTCGTAAATCCTCATCA | AA | ATTCCATGACATTTGCATTCACGCC | CACGTGCGTACAGTACATGAACCTG | AA | ATCATCCAGTAAACCGCC |
| **9** | CCTCGTAAATCCTCATCA | AA | CTTCATTGTTGTGCAGATTCATTAT | TCTCATTTGAAACATGCGAACGACC | AA | ATCATCCAGTAAACCGCC |
| **10** | CCTCGTAAATCCTCATCA | AA | ATCAACAAAGGCTCGCGCAAACTTA | TCTGAGATCTCGTCCTCTTTCAGCT | AA | ATCATCCAGTAAACCGCC |
| **11** | CCTCGTAAATCCTCATCA | AA | CCCCACTCCCAGTCCAGACCATTTG | CCAAATTCGATATTATCCGAACAGC | AA | ATCATCCAGTAAACCGCC |
| **12** | CCTCGTAAATCCTCATCA | AA | ATGTCTCTATCAAACCTTCGCTGCA | TCCGATTATTACGATAATCACATGT | AA | ATCATCCAGTAAACCGCC |
| **13** | CCTCGTAAATCCTCATCA | AA | ACTCGTTATCGCATAAACGAACGCA | TCTAGAAATTGAATGCGCCACTGCC | AA | ATCATCCAGTAAACCGCC |
| **14** | CCTCGTAAATCCTCATCA | AA | CCAAAAATTCCTTTGCCTTTCATGT | TCTCTGCAACCCCTTTGTACAATTC | AA | ATCATCCAGTAAACCGCC |
| **15** | CCTCGTAAATCCTCATCA | AA | CTTTGAATTGGTTTTGGCATTCGTT | CGGTTGTGGGACAGTTCCATCGTCT | AA | ATCATCCAGTAAACCGCC |
| **16** | CCTCGTAAATCCTCATCA | AA | AATGAGAACTCCAGGATTGTCCCTG | TGCCATTTTCATGGCTTTGGCAACG | AA | ATCATCCAGTAAACCGCC |
| **17** | CCTCGTAAATCCTCATCA | AA | TGTGCCGCCAGTAATTGAATTAGTG | ACGTAAAGGAACATTAATACCGTTA | AA | ATCATCCAGTAAACCGCC |
| **18** | CCTCGTAAATCCTCATCA | AA | CACCCGAGTGTGCGATTGCCCACCA | GCGGGTCTAGTACTAGATTTGTTGG | AA | ATCATCCAGTAAACCGCC |
| **19** | CCTCGTAAATCCTCATCA | AA | TCTTGGCATCAATCTTGACGGGCAT | ATCCCTTTCCCCGACCGCCTTTGCC | AA | ATCATCCAGTAAACCGCC |
| **20** | CCTCGTAAATCCTCATCA | AA | TATAATAGCACAAACACCGACTCTA | AATGATCAGAGACATGATCAGCATC | AA | ATCATCCAGTAAACCGCC |
| **21** | CCTCGTAAATCCTCATCA | AA | TTGTTACATGTCCTCAAACGCATGG | GTCCAAATGGCACTTAGAGGCACCC | AA | ATCATCCAGTAAACCGCC |
| **22** | CCTCGTAAATCCTCATCA | AA | GCCCGATCGGCCGACACGGCCGCAC | AAACAGCAAACACTAATTTGATGTT | AA | ATCATCCAGTAAACCGCC |
| **23** | CCTCGTAAATCCTCATCA | AA | TGAGTCGACACGTGCGCTCTTATCA | AAAAAGTTTCCAAATCTGCGTTCAG | AA | ATCATCCAGTAAACCGCC |
| **24** | CCTCGTAAATCCTCATCA | AA | TCGTATTTCATGCAGAGGCAAGTCC | CTTTGGTCTTGAAATGCCAGAACGC | AA | ATCATCCAGTAAACCGCC |
| **25** | CCTCGTAAATCCTCATCA | AA | TATTTACCAGTGCTTTACTGCTCGA | GTTCTGAGCCCACGCCCGCTTCAAC | AA | ATCATCCAGTAAACCGCC |
| **26** | CCTCGTAAATCCTCATCA | AA | AGAAAAAGTAAAGTTTCACAAACGC | TCAAAACCAGCCAACAAAGAGATGA | AA | ATCATCCAGTAAACCGCC |
| **27** | CCTCGTAAATCCTCATCA | AA | AGACTTTCAGTTGGATAGACTTGGC | TCAATCAACTCTATCTATGAATTGA | AA | ATCATCCAGTAAACCGCC |
| **28** | CCTCGTAAATCCTCATCA | AA | CGATGTCTGGTTAAGAATCAGAACT | TGTCGTTTCTCCGATTCAGAGAAGT | AA | ATCATCCAGTAAACCGCC |
| **29** | CCTCGTAAATCCTCATCA | AA | AGGGACCCAGAATTGGGTGGTCTCC | GAATAAATATTCATTAGTATAATTT | AA | ATCATCCAGTAAACCGCC |
| **30** | CCTCGTAAATCCTCATCA | AA | TTAGATAGTAGACCATTGATTGGAG | AAATAGAAAATTCGCGTCATTGTGA | AA | ATCATCCAGTAAACCGCC |
| **31** | CCTCGTAAATCCTCATCA | AA | TTTGCGAGCCGTGAAGTTTCTAATT | AGGACAAAAGAATGTGTGAACTTAT | AA | ATCATCCAGTAAACCGCC |
| **32** | CCTCGTAAATCCTCATCA | AA | TTCTTAAGTCACGAAATATACCTGA | TCTTTGGACAAGGAACACGCCTTTT | AA | ATCATCCAGTAAACCGCC |
| **33** | CCTCGTAAATCCTCATCA | AA | TTGCTCACACTATATTCCGAAGAGT | TCCTTACGCTATATTCCTGACACTT | AA | ATCATCCAGTAAACCGCC |
| **34** | CCTCGTAAATCCTCATCA | AA | TGCTTATACACTGTTGCTTACACTC | CTGAACATGCTATTGCTTATACTCT | AA | ATCATCCAGTAAACCGCC |
| **35** | CCTCGTAAATCCTCATCA | AA | TTCCTTACACTCTGTTCCTCACACT | TCCTTATACACTATTGCTTGCACTC | AA | ATCATCCAGTAAACCGCC |
| **36** | CCTCGTAAATCCTCATCA | AA | ATCCCTCATACTCTATTGCCAACAC | TTCCTCACACTCTATTCCTTATACT | AA | ATCATCCAGTAAACCGCC |
| **37** | CCTCGTAAATCCTCATCA | AA | CAAATTGAATGAAATTCATTCCTTG | CCCTGAAGGCAGATATTATAGTGGA | AA | ATCATCCAGTAAACCGCC |
| **38** | CCTCGTAAATCCTCATCA | AA | AGGCATAAAAACAACTTAATCCTAT | GTCGTCATTCGTGTTGCGCTATAGA | AA | ATCATCCAGTAAACCGCC |
| **39** | CCTCGTAAATCCTCATCA | AA | GAACCATTTGATGCTTCCCTAACAC | AGCAAAGACTGAACCATGAGATGAA | AA | ATCATCCAGTAAACCGCC |
