## Supplementary figures and images for "The Expression of *Pax6* Genes in an Eyeless Arachnid Suggests Their Ancestral Role in Arachnid Head Development"

### Figure S1

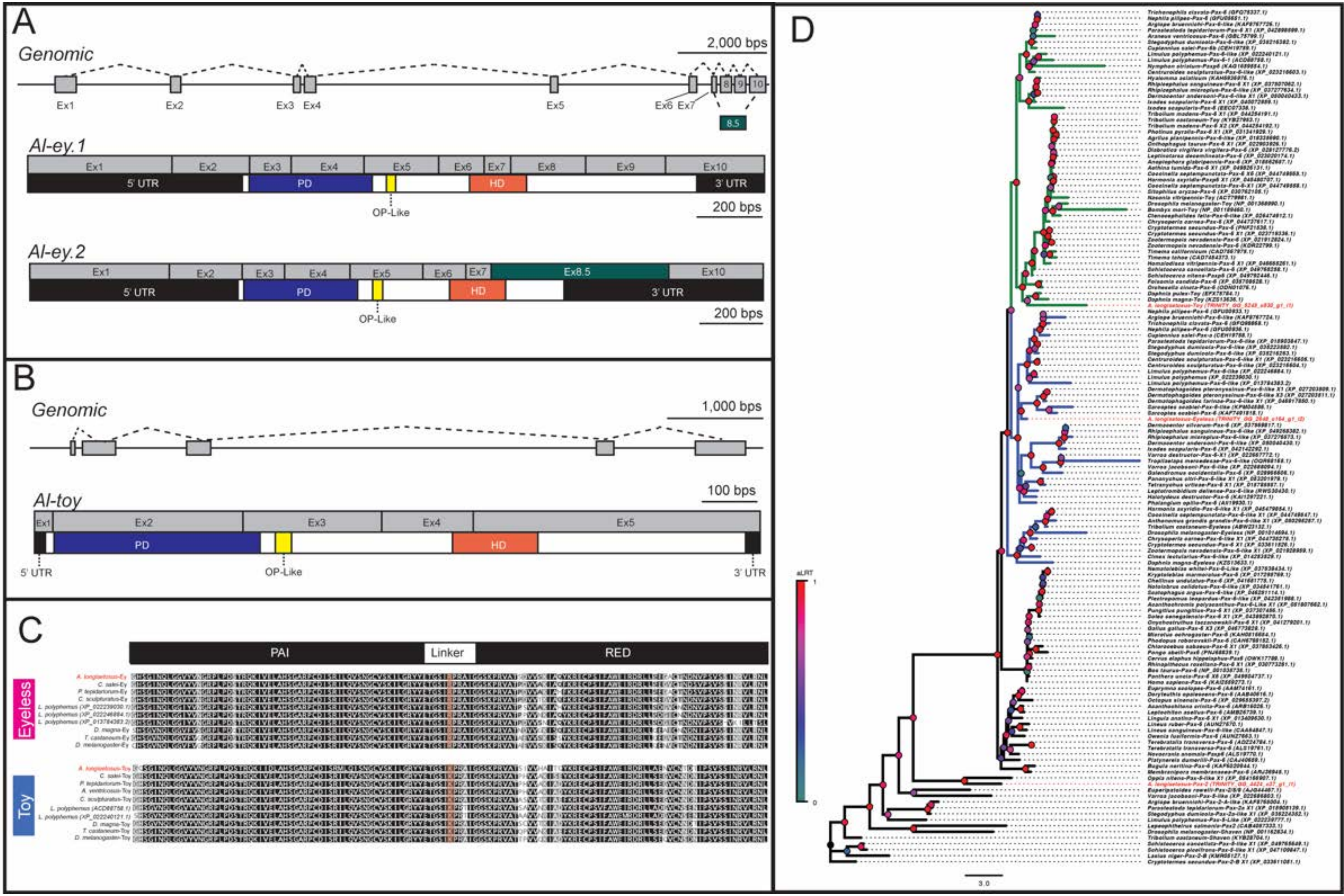

### Figure S2

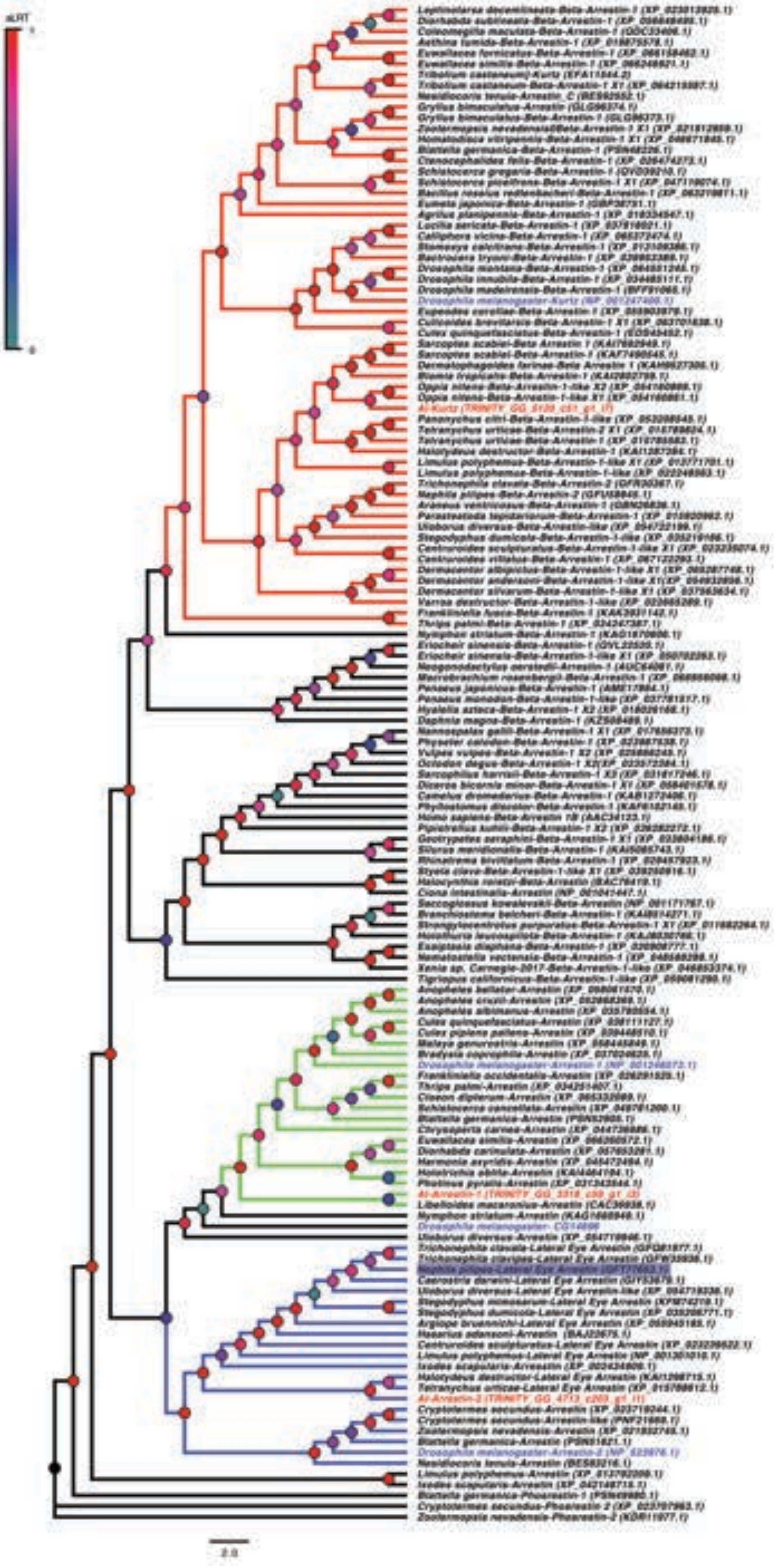
